# A CXCR4 partial agonist improves immunotherapy by targeting polymorphonuclear myeloid-derived suppressor cells and cancer-driven granulopoiesis

**DOI:** 10.1101/2024.10.09.617228

**Authors:** Jin Qian, Chenkai Ma, Quin T. Waterbury, Xiaofei Zhi, Christine S. Moon, Ruhong Tu, Hiroki Kobayashi, Feijing Wu, Biyun Zheng, Yi Zeng, Hualong Zheng, Yosuke Ochiai, Ruth A. White, David W. Harle, Jonathan S. LaBella, Leah B. Zamechek, Lucas ZhongMing Hu, Ryan H. Moy, Arnold S. Han, Bruce Daugherty, Seth Lederman, Timothy C. Wang

## Abstract

Polymorphonuclear myeloid-derived suppressor cells (PMN-MDSCs) are pathologically activated neutrophils that potently impair immunotherapy responses. The chemokine receptor CXCR4, a central regulator of hematopoiesis, represents an attractive PMN-MDSC target1. Here, we fused a secreted CXCR4 partial agonist TFF2 to mouse serum albumin (MSA) and demonstrated that TFF2-MSA peptide synergized with anti-PD-1 to induce tumor regression or eradication, inhibited distant metastases, and prolonged survival in multiple gastric cancer (GC) models. Using histidine decarboxylase (Hdc)-GFP transgenic mice to track PMN-MDSC *in vivo*, we found TFF2-MSA selectively reduced the immunosuppressive Hdc-GFP^+^ CXCR4^hi^ tumor PMN-MDSCs while preserving proinflammatory neutrophils, thereby boosting CD8^+^ T cell-mediated anti-tumor response together with anti-PD-1. Furthermore, TFF2-MSA systemically reduced PMN-MDSCs and bone marrow granulopoiesis. In contrast, CXCR4 antagonism plus anti-PD-1 failed to provide a similar therapeutic benefit. In GC patients, expanded PMN-MDSCs containing a prominent CXCR4^+^LOX-1^+^ subset are inversely correlated with the TFF2 level and CD8^+^ T cells in circulation. Collectively, our studies introduce a strategy of using CXCR4 partial agonism to restore anti-PD-1 sensitivity in GC by targeting PMN-MDSCs and granulopoiesis.

## Introduction

Polymorphonuclear myeloid-derived suppressor cells (PMN-MDSCs) are a major cellular component in advanced solid tumors that hinders anti-tumor immunity. These cells comprise a heterogeneous group of predominantly immature neutrophils pathologically programmed by cancer-derived inflammatory signals^2^. Despite being short-lived, their continuous replenishment from aberrant myelopoiesis in the bone marrow (BM) sustains their large number and potent immunosuppression in the tumor microenvironment (TME)^3^.

Emerging evidence has delineated distinct development trajectories and transcriptional features of PMN-MDSCs compared with classical neutrophils (PMNs)^3–6^. However, specific markers for distinguishing PMN-MDSCs from the anti-tumor neutrophils, especially in murine models, remain lacking^2,7^. This has complicated the preclinical evaluation of therapies against PMN-MDSCs7, despite recent studies highlighting the crucial role of a subset of tumor-associated neutrophils for immunotherapy success^8–12^. Existing therapeutic approaches such as neutrophil depletion mainly target circulating neutrophils in the periphery, often leading to compensatory overproduction from BM progenitors, and thus limiting treatment efficacy^13–16^. Histidine decarboxylase (Hdc) is a unique enzyme that catalyzes the synthesis of histamine, and our previous work established it as a marker for a subpopulation of immature neutrophils that predominates in cancer and thus potentially a useful PMN-MDSC marker^17^. Here, we utilized a transgenic mouse model as the tumor host, enabling the tracking and targeting of PMN-MDSCs throughout their development *in vivo*^17–19^.

The chemokine receptor CXCR4, controlled by its agonist ligand SDF-1 (stromal cell-derived factor-1, also named CXCL12), plays pleiotropic functions in diverse physiological processes such as hematopoiesis, myelopoiesis, immune cell development, survival, and trafficking^1,20^. However, the SDF-1-CXCR4 axis can be exploited by many cancers to orchestrate an immunosuppressive microenvironment through PMN-MDSC recruitment and T cell exclusion^21–23^. Hence, CXCR4 antagonists, clinically useful for hematopoietic stem cell mobilization, have been repurposed in clinical trials to treat cancer in combination with immune checkpoint inhibitors^24,25^. However, to date, these attempts have shown limited therapeutic benefit^16,26^, likely owing to the essential role of CXCR4 in maintaining BM homeostasis and normal immune function^27–29^. Interestingly, SDF-1 overexpression or supplementation can also inhibit tumor progression, suggesting that some level of CXCR4 signaling may be beneficial for anti-tumor immunity^30–33^.

Partial CXCR4 agonists, yet to be tested in this setting, have the potential to block excessive action by full agonists while sustaining a moderate level of receptor activation, thus avoiding the detrimental effects of complete receptor blockade. Trefoil Factor 2 (TFF2), a small secreted peptide of the trefoil factor family, has been shown to act as a partial agonist of CXCR4, dampening SDF-1-mediated intense signaling while eliciting weak CXCR4 activation^34–36^.

Interestingly, the *Tff2* gene is methylated and epigenetically silenced during gastric preneoplasia, contributing to the development of gastric cancer^37^. *Tff2* deficiency in mice also accelerates colon cancer growth, whereas its overexpression minimizes tumor burden by suppressing the development of MDSCs^38,39^. Our previous studies have shown that gastric cancer treatment with immune checkpoint therapy is enhanced by the co-administration of drugs that suppress MDSC^40^. Here, we fused TFF2 with murine serum albumin (MSA) to develop a novel peptide TFF2-MSA (**Fig. 1A**) with enhanced stability (data not shown). In combination with anti-PD-1 immunotherapy, it shows tremendous efficacy in treating advanced gastric cancer by selectively and systemically targeting PMN-MDSCs.

**Fig. 1.**
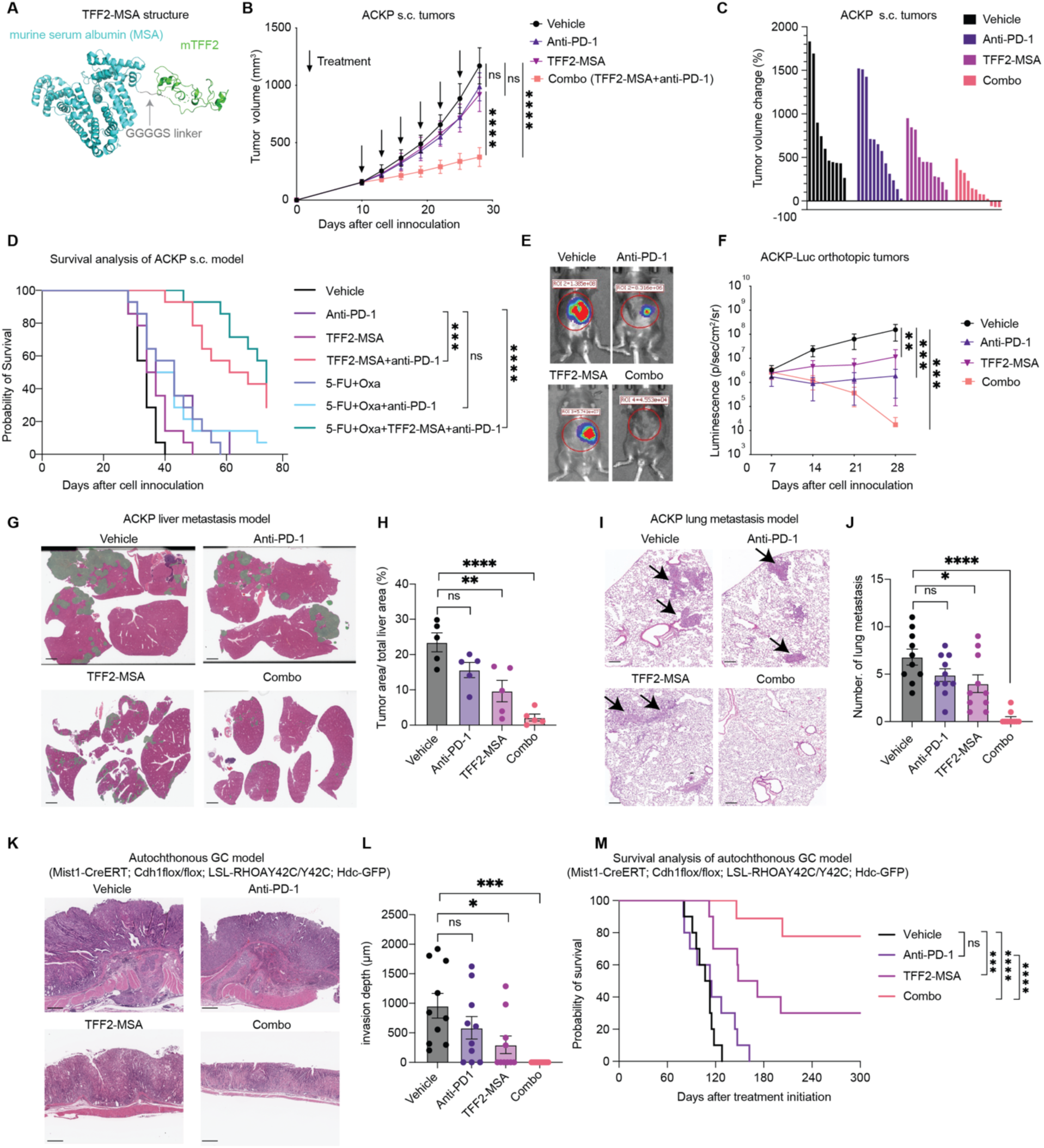
TFF2-MSA elicits robust tumor inhibition in combination with anti-PD-1 in immunocompetent mouse models of GC. **A.** Artistic representation of fusion structure of TFF2-MSA. **B.** Subcutaneous ACKP tumor growth curve in Hdc-GFP host mice that received the treatments every three days as indicated by black arrows (n=10-12). **C.** Tumor volume changes relative to the initial volume of each tumor. **D**. Survival curve of ACKP tumor-bearing mouse survival in response to the indicated treatments (n=14). 5-FU, 5-fluorouracil. Oxa, oxaliplatin. **E, F.** Representative bioluminescence images and luminescence quantification showing orthotopic stomach tumors from luciferase-expressing ACKP cells following treatments (n=5). **G, H.** Representative images of H&E staining and quantification of the liver metastases (colored in green) from portal vein injected ACKP cells following treatments (n=5). Scale bar, 2mm. **I, J.** Representative images of H&E staining and quantification of the spontaneous lung micrometastases following surgical resection of ACKP tumors and treatments (n=10 per group). Arrows indicate lung metastases. Scale bar, 100μm. **K, L.** Representative images of H&E staining and quantification of tumor invasion depth following treatments (n=10). Representative macroscopic images of autochthonous stomach tumors in Mist1-CreERT; Cdh1^flox/flox^; LSL-RHOA^Y42C/Y42C^; Hdc-GFP mice at 10 months post tamoxifen induction following the indicated treatments (n=10 per group). **M.** Survival curve of autochthonous GC-bearing Mist1-CreERT; Cdh1^flox/flox^; LSL-RHOA^Y42C/Y42C^; Hdc-GFP mice receiving the indicated treatments (n= 10 per group). Data is presented as mean ± SEM, and P values are calculated by two-way ANOVA (**B, F**), one-way ANOVA (**H, J, L**) and log-rank test (**D, M**). *P < 0.05, **P < 0.01, ***P < 0.001, ****P < 0.0001, ns, not significant.

## Results

### TFF2-MSA acts synergistically with anti-PD-1 to inhibit tumor growth and metastasis and prolong mouse survival

TFF2-MSA, both alone and in combination with anti-PD-1 antibody, was first studied in syngeneic mouse models of GC. The ACKP cancer cell line, derived from a genetically engineered mouse (Atp4b-Cre;Cdh1^-/-^;LSL-Kras^G12D/+^;Trp53^-/-^) that develops mixed-type GC with widespread metastases,^41^ was injected subcutaneously into Hdc-GFP transgenic mice. When the tumors reached an average volume of 150-250 mm^3^, treatments were initiated with intraperitoneal injections of the vehicle, TFF2-MSA, anti-PD1 antibody, or TFF2-MSA plus anti-PD-1 antibody (Combo therapy) every three days (**Fig. S1A-B**). Although ACKP tumors were refractory to either anti-PD-1 or TFF2-MSA alone, the combination therapy displayed robust synergy in tumor inhibition (**Fig. 1B**), resulting in tumor regression in a quarter of the mice (3 of 12) (**Fig. 1C**). Additionally, splenic enlargement in tumor-bearing mice, characteristic of cancer-driven myelopoiesis^38^, was markedly alleviated with TFF2-MSA or combination treatments (**Fig. S1C**).

Standard chemotherapy (5-FU+oxaliplatin), used as first-line GC patient treatment^40^, was further incorporated into treatment comparisons^40^. Of all tested treatments, only the TFF2-MSA plus anti-PD-1 combination, with or without chemotherapy, significantly suppressed tumor growth (**Fig. S1D**), and extended median survival of mice (64, and 73 days respectively, vs. 34 days in vehicle, **Fig. 1D**). In contrast, the immune-chemotherapy combination offered limited survival benefit (median survival 38.5 days, **Fig. 1D, Fig. S1D**) in this aggressive GC model. The combination of TFF2-MSA plus anti-PD-1 caused no weight loss as seen in chemotherapy or its combination with anti-PD-1 (**Fig. S1E**). Considering the possible difference of TME between subcutaneous and orthotopic tumor models, we next developed a syngeneic orthotopic model by injecting luciferase-expressing ACKP cells into the mouse stomach submucosa layer and starting treatments after 7 days. Weekly bioluminescence imaging of stomach tumors revealed only the combo regimen eradicated 80% of orthotopic stomach tumors compared to 0% in either monotherapy, as confirmed by histopathology, leading to long-term survival in the cured mice (**Fig. 1E-F, Fig. S1F-I**). Cancer-induced splenomegaly was simultaneously ameliorated by TFF2-MSA and Combo therapy (**Fig. S1J**).

Advanced GC frequently metastasizes to the liver or lung, resulting in a dismal prognosis^42^. Tumor-induced PMN-MDSCs facilitate such cancer metastases^43–45^. In a model of liver metastases resulting from portal vein injection of ACKP cells, TFF2-MSA inhibited liver metastases by 59%, while the combo regimen achieved a more than 90% reduction in metastatic lesions and a nearly 2-fold increase in median survival (**Fig. 1G-H, Fig. S1K-L**). In a model of spontaneous lung metastases following surgical resection of subcutaneous established ACKP tumor, TFF2-MSA reduced micro-metastatic nodules while the combination of TFF2-MSA plus anti-PD-1 led to near complete elimination of lung metastases, resulting in significantly extended survival (**Fig.1I-J**). In addition to the ACKP model, a new syngeneic PC (Trp53^-/-^; CCNE1) organoid line representing intestinal-type GC was generated and transplanted into the Hdc-GFP mice for treatment evaluations (**Fig. S2A**). Although the PC tumors were relatively sensitive to anti-PD-1, the addition of TFF2-MSA improved PC tumor inhibition and mouse survival compared to anti-PD-1 antibody alone (**Fig. S2B**).

Next, we generated a novel autochthonous mouse GC model with inducible RhoA^Y42C^ mutation and CDH1 loss (Mist1-CreERT; Cdh1^flox/flox^; LSL-RHOA^Y42C/Y42C^; Hdc-GFP) to model the diffuse-type GC patients with the worst survival among all GC subtypes^42^. Eight months after tamoxifen induction, mice developed poorly differentiated diffuse-type GC interspersed with signet ring cells, confirmed by an expert pathologist (**Fig. S2C**). Treatment was then initiated with TFF2-MSA, anti-PD-1, or their combination for 2 months (**Fig. S2D**). While GC invaded the stomach wall resulting in its thickening resembling “linitis plastica” in the vehicle or anti-PD-1 alone treated group, TFF2-MSA restrained the majority of the tumor lesions to the mucosal layer, and only atypical foci were present after Combo treatment (**Fig. 1K-L, Fig. S2E-G**). The total tumor area was simultaneously decreased with the Combo regimen, accompanied by reduced splenomegaly (**Fig. S2H-J**). Notably, 80% of the mice in combo therapy group survived beyond 300 days, remaining free of GC as confirmed by histology, whereas all mice treated with anti-PD-1 monotherapy succumbed within 6 months (**Fig. 1M)**.

Finally, TFF2-MSA also improved anti-PD-1 efficacy in other GI cancer models, including a subcutaneous CT26 colon cancer model (**Fig. S3A-C**), an orthotopic CT26-luciferase colon cancer model (**Fig. S3D-E**), and a subcutaneous Panc02 pancreatic cancer model (**Fig. S3F**). Taken together, these data from multiple immunocompetent models demonstrate that TFF2-MSA acts in robust synergy with anti-PD-1 to reduce tumor growth and extend mouse survival.

### TFF2-MSA selectively reduces immunosuppressive Hdc-GFP^+^ PMN-MDSCs in the tumor microenvironment

Next, we investigated whether TFF2-MSA resensitized ACKP tumors to immunotherapy by targeting tumor-associated PMN-MDSCs. First, in the Hdc-GFP host mice of our subcutaneous ACKP tumor model, we compared morphological and functional features of Hdc-GFP^+^ (Hdc^+^) and Hdc-GFP^-^ (Hdc^-^) subsets within the CD45^+^CD11b^+^LY6G^+^LY6C^-/low^ tumor-associated PMNs. DRAQ5 nuclear staining and unbiased nuclear lobe counting using imaging flow cytometry revealed that the Hdc^+^ PMNs exhibited an immature granulocyte morphology, with poor nuclear segmentation compared to their Hdc^-^ counterparts **(Fig. 2A-B, Fig. S4A**). Furthermore, Hdc^+^ tumor-associated PMNs potently suppressed T cell proliferation in coculture with T cells, contrasting to Hdc^-^ PMNs that promoted T cell proliferation (**Fig. 2C, Fig. S4B)**. The immunosuppressive function of Hdc^+^ tumor PMNs depends in part on arginase and iNOS (inducible nitric oxide synthase) (**Fig. S4C**). Using single-cell RNA sequencing (scRNA-seq) of isolated CD45^+^ cells from vehicle-treated ACKP tumors in Hdc-GFP mice (as detailed below), we found numerous reported PMN-MDSC or pro-tumor neutrophil markers (*CD14*^5^, *Cd300ld*^46^, *Jaml*^4^, *S100a8/9*^47^, *Entpd1*^48^, *Slfn4*^49^, etc.) and CXCR4 enriched in these Hdc^+^ PMNs compared with the Hdc^-^ PMN subset (**Fig. S4D, Table S1**), thus validating this Hdc^+^ LY6G^+^ PMN subset as *bona fide* PMN-MDSCs.

**Fig. 2.**
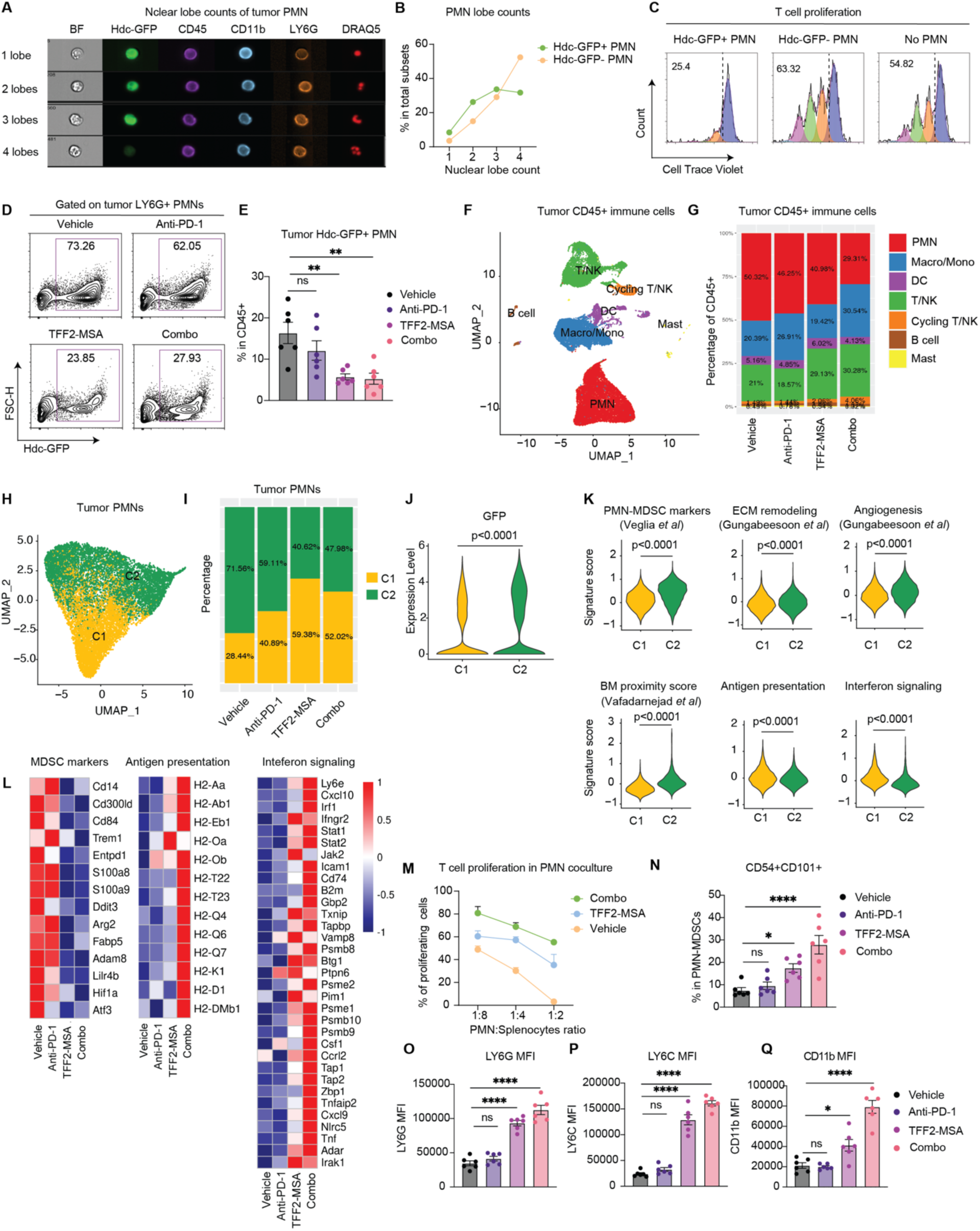
TFF2-MSA selectively reduced immunosuppressive PMN-MDSCs in the tumor microenvironment. **A.** Nuclear segmentation of Hdc-GFP^+^ and Hdc-GFP^+^ tumor PMNs (pooled from n=3 mice) analyzed at 60x by Imagestream imaging cytometer. Representative images of BF, PMN markers, and DRAQ5 nuclear staining are shown for the CD45^+^CD11b^+^LY6G^+^ PMNs scored as having 1-4 nuclear lobes. BF, bright field. **B**. Proportions of cells with 1-4 nuclear lobes within Hdc-GFP^+^ and Hdc-GFP^-^ PMN subsets. **C**. Representative flow cytometry plots of cell trace violet staining showing T cell proliferation after coculture with Hdc-GFP^+^ or Hdc-GFP^-^ tumor PMNs at a PMN to splenocyte ratio of 1:8, or without PMNs. **D**. Representative flow cytometry plots showing Hdc-GFP^+^ PMN-MDSCs within tumor LY6G^+^ PMNs following treatments. **E**. Percentage of Hdc-GFP^+^LY6G^+^ tumor PMN-MDSCs within tumor CD45^+^ cells following treatments (n = 6 mice). **F**. ScRNA-seq analysis of CD45^+^ cells isolated from ACKP tumors (pooled from n=3 per group) in Hdc-GFP mice treated with vehicle, anti-PD-1, TFF2-MSA, or their combination. UMAP dimensional reduction is colored by cell types. DC, dendritic cells; NK, natural killer cells; Macro, macrophages; Mono, monocytes. **G**. Relative frequency of each immune cell type within tumor CD45^+^ cells from the treatment groups. **H**. UMAP depicting 2 subclusters of tumor PMN. **I**. Relative frequency of 2 subclusters within tumor PMNs following treatments. **J**. Hdc-GFP expression in the 2 subclusters of tumor PMNs. **K**. Expression score of gene signature obtained from prior studies^2,8,50^ in tumor PMN subclusters. ECM, extracellular matrix. BM, bone marrow. **L**. Heatmap showing manually selected gene expressions in tumor PMNs following treatments. **M**. Representative flow cytometry plots of cell trace violet staining showing T cell proliferation after coculture with tumor PMNs sorted from different treatment groups. **N**. Frequency of CD54^+^CD101^+^ cells within tumor PMNs following treatments (n=3). **O-Q**. Expression of LY6G, LY6C, CD11b on tumor PMNs following treatments (n=3), expressed as mean fluorescence intensity (MFI). Data is presented as mean ± SEM, and P values are calculated by one-way ANOVA (**D, N-Q**) or Wilcoxon test (**J, K**). *P < 0.05, **P < 0.01, ***P < 0.001, ****P < 0.0001, ns, not significant.

While TFF2-MSA only moderately decreased total tumor PMNs (**Fig. S4E-F**), it profoundly reduced Hdc^+^ tumor PMN-MDSCs (by 65% in TFF2-MSA alone and 68% in Combo group, respectively) (**Fig. 2D-E**). To gain broader insights into the impact of TFF2-MSA on the immune microenvironment, we isolated total CD45^+^ immune cells for scRNA-seq from ACKP tumors that were treated with vehicle, anti-PD-1, TFF2-MSA or TFF2-MSA plus anti-PD-1 (see Method for details). Low-resolution clustering identified major clusters of PMNs, T cells and NK cells (T/NK), cycling T/NK cells, macrophages/monocytes, dendritic cells (DC), B cells, and mast cells (**Fig. 2F, Fig. S5A**). The most prominent change following TFF2-MSA treatment was PMN contraction accompanied by T/NK cell expansion, and Combo treatment further augmented the increased ratio of T/NK cells to PMNs (**Fig. 2G**).

Unsupervised clustering of the tumor PMN population identified 2 major subsets, C1 and C2. TFF2-MSA increased the relative ratio of C1 to C2 in the remaining tumor PMNs (**Fig. 2H-I**). C2 subset was enriched with GFP, PMN-MDSC markers, signatures of tumor angiogenesis^8^, ECM remodeling^8^, and BM proximity score suggesting neutrophil immaturity^50^ (**Fig. 2J-K**). Conversely, the C1 subset was enriched for antigen presentation and interferon-stimulated genes (ISGs, including prototypical *Cxcl10* and *Ly6e*), all neutrophil predictors for immunotherapy success^8,9,11,12^ (**Fig. 2K**). TFF2-MSA and Combo treatments rendered the total tumor PMNs to exhibit lower PMN-MDSC markers than those from vehicle or anti-PD-1 alone, as confirmed by flow cytometry showing downregulation of Arginase 1, and upregulation of MHC molecules in TFF2-MSA and Combo groups (**Fig. 2L, Fig. S6A-B**). Functionally, TFF2-MSA treatment abrogated the inhibitory effect of total tumor PMNs on T cell division, and Combo-treated PMNs further enhanced T cell proliferation, suggesting their switch to immunostimulatory PMNs (**Fig. 2M, Fig. S6C**). In concert with these data, a recently identified anti-tumor mature CD54^+^CD101^+^ neutrophil subset in GC^51^ was augmented by Combo treatment (**Fig. 2N**). PMNs from TFF2-MSA- and Combo-treated mice also displayed increased surface expression of LY6G, LY6C, CD11b, CD101, all suggestive of PMN maturation towards more functional status^8^ **(Fig. 2O-Q, Fig. S6D)**.

We asked whether TFF2-MSA also altered gene expression profiles within Hdc^+^ tumor PMN-MDSCs. Across all treatment groups, PMN-MDSC markers were consistently enriched in C2 compared to C1, while TFF2-MSA preferentially modulated their levels in the C2 subset (**Fig. S6E**). Decreased expression of the PMN-MDSC markers *Arg1* and *Cd14* and increased expression of neutrophil differentiation/maturation marker *Irf8*^52^ and *Cd101*^14^ were found in Hdc^+^ PMN-MDSCs from TFF2-MSA and Combo-treated mice (**Fig. S6F-G**). These observations demonstrated that TFF2-MSA not only reduced PMN-MDSC numbers but also modulated the PMN-MDSC phenotypes.

We wondered whether TFF2-MSA directly reprogrammed the Hdc^+^ C2 subset into the Hdc-GFP^-^ C1 subset. RNA velocity analysis suggested this was unlikely, as the C1 and C2 subpopulations showed distinct differentiation trajectories with separate progenitor origins (**Fig. S6H**). In support of this, sorted Hdc^+^ PMN from either tumor or matched spleen failed to switch to GFP^-^ during *in vitro* culture, even with the addition of TFF2-MSA **(Fig. S6I)**.

Interestingly, although the Hdc^+^ fraction of PMNs was rarely found in normal stomachs of healthy mice, this fraction significantly expanded in the stomachs of Mist1-CreERT; Cdh1^flox/flox^; LSL-RHOA^Y42C/Y42C^; Hdc-GFP mice with poorly differentiated GCs, a change that was reversed by TFF2-MSA treatment or its combination with anti-PD-1 (**Fig. S7A-C**). Overall, our data place Hdc^+^ PMN-MDSCs at the center of immunosuppression in the TME, with TFF2-MSA effectively alleviating this key immunotherapy barrier.

### TFF2-MSA functions as a CXCR4 partial agonist

We next studied whether TFF2-MSA effects on the TME could be attributed to its known target CXCR4^34^. Both single-cell transcriptomic and flow cytometric analysis revealed that Hdc^+^ PMN-MDSCs account for the majority of CXCR4^+^ immune cells in the TME and highly expressed CXCR4, although CXCR4 was also expressed at lower levels in other immune cells (**Fig. S8A-B**). To track TFF2-MSA effects on CXCR4^+^ cells *in vivo*, we employed a Cxcr4-GFP reporter mouse, as TFF2-MSA interferes with antibody binding to CXCR4^34^. Flow cytometric examination of ACKP tumors in Cxcr4-GFP mice confirmed the high level of CXCR4 on PMNs (**Fig. S8C**). TFF2-MSA treatment reduced the proportion of CXCR4^+^ cells within LY6G^+^ PMNs in both tumor and blood (**Fig. S8D)**.

To clarify the action of TFF2-MSA on CXCR4, we transfected the human myelogenous leukemia cell line K562, which lacks endogenous CXCR4 expression^53^, with a HiBiT-tagged Cxcr4 construct. Both TFF2-MSA and its human version TFF2-HSA induced HiBiT-tagged Cxcr4 internalization, leading to a reduced membrane-bound HiBiT signal in a dose-dependent manner. However, the effect was markedly weaker than that of SDF-1, confirming TFF2-MSA’s lesser activity on CXCR4 desensitization as a partial agonist (**Fig. S8E**). In chemotaxis assays, TFF2-MSA preferentially acted on Hdc^+^ PMNs with a bell-shaped concentration-response curve, behaving as a partial agonist, unlike SDF-1 which strongly attracts both Hdc^+^ and Hdc^-^ PMNs (**Fig. S8F-G**). Our previous study demonstrated TFF2 antagonized Jurkat cell chemotaxis in response to an SDF-1 gradient, consistent with its partial blockade of full agonist activity^34^. PI3K-Akt signaling is a major downstream pathway of CXCR4 activation^34^. We observed that TFF2-MSA impaired SDF-1-mediated Akt activation and weakly activated Akt signaling by itself, which was blocked by the CXCR4 antagonist AMD3100 (**Fig. S8H**), indicating its dependence on CXCR4. These data suggest the inhibition of SDF-1-mediated chemotaxis and signaling by TFF2-MSA may at least partially contribute to its selective reduction of Hdc^+^ PMNs *in vivo*.

### TFF2-MSA in conjunction with anti-PD-1 induces a profound cytotoxic CD8 T cell response that mediates tumor control

Having established TFF2-MSA’s inhibition of PMN-MDSCs, we next defined the effects of TFF2-MSA on T cells. Our flow cytometry revealed increased T cells with TFF2-MSA alone and with Combo therapy, with a much greater expansion in CD8^+^ T cells compared to CD4^+^ T cells or Tregs (**Fig. 3A-B, Fig. S9A-B**). Whole tumor section staining revealed a marked influx of CD8^+^ T cells in response to TFF2-MSA, with deep infiltration into the tumor core with Combo therapy, suggestive of active tumor killing (**Fig. 3C-D, Fig. S9C**). A significant phenotypic response in the intra-tumoral CD8^+^ T cells was observed following Combo therapy, characterized by an increased frequency of both Tim3^+^Granzyme B^+^ and Tim3^-^Granzyme B^+^ cytotoxic CD8^+^ T cells, alongside elevated levels of Ki67^+^ proliferative and CD69^+^ activated CD8^+^ T cells (**Fig. 3E-F, Fig. S9D**). Although TFF2-MSA monotherapy boosted the numbers of multifunctional IFNγ^+^TNFα^+^ CD8^+^ T cells, the combo regimen more effectively expanded this population (**Fig. 3G**).

**Fig. 3.**
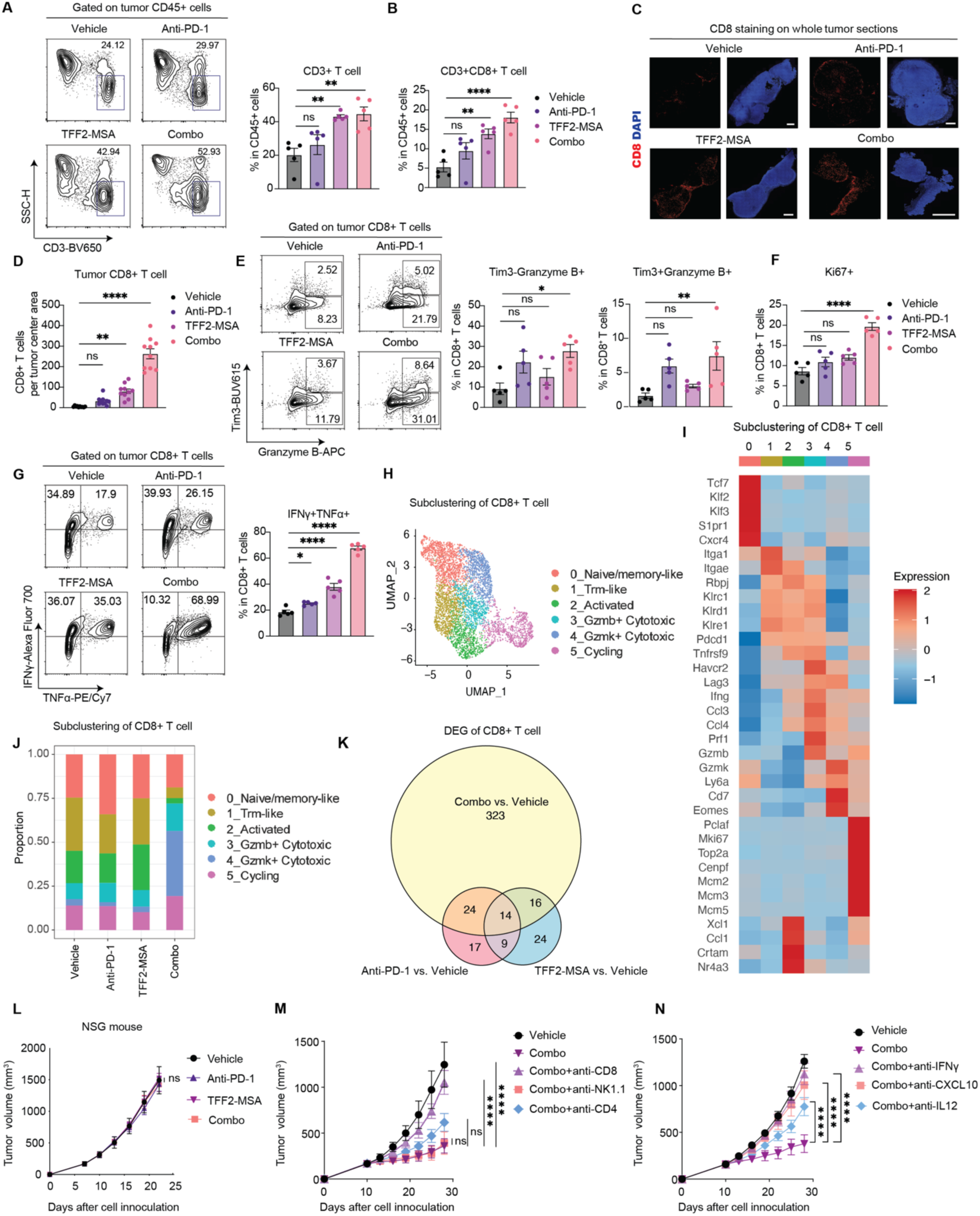
TFF2-MSA in combination with anti-PD-1 rejuvenates CD8 T cell-mediated anti-tumor immunity. **A.** Representative flow cytometry plots and quantification of CD3^+^ T cells in ACKP tumors following treatments (n=5). **B**. Quantification of CD8^+^ T cells in ACKP tumors following treatments (n=5). **C**. Representative images of CD8 (red) and DAPI (blue) staining in whole ACKP tumor sections following treatments (n=5). Scale bar, 100μm. **D**. Quantification of CD8 staining per field in the tumor center area (>1/10 diameter from the edge) in **C** (n=10 random fields). **E**. Representative flow cytometry plots and quantification of Tim3^-^Granzyme B^+^ and Tim3^+^Granzyme B^+^ cells within CD8^+^ cells from ACKP tumors following treatments (n=5). **F**. Quantification of Ki67^+^ cells within CD8^+^ cells from ACKP tumors following treatments (n=5). **G**. Representative flow cytometry plots and quantification of IFNγ^+^TNFα^+^ within CD8^+^ T cells from ACKP tumors following treatments (n=5). **H.** UMAP depicting subclusters of the intratumoral CD8^+^ T cells. **I.** Heatmap of manually selected marker genes used for defining CD8^+^ T cell subclusters. **J**. Relative frequency of intratumoral CD8^+^ T cell subclusters following treatments. **K.** Venn diagram depicting differentially expressed genes (DEGs) in intratumoral CD8^+^ T cells in the indicated treatments versus vehicle control. Numbers in the circles indicate the number of DEGs. L. Growth curve of subcutaneously implanted ACKP tumor in NOD.Cg-Prkdc^scid^ Il2r^gtm1Wjl^/SzJ (NSG) mice following treatments (n=6). **M**, **N.** Growth curve of subcutaneously implanted ACKP tumors following treatments (n=10). Data is presented as mean ± SEM, and P values are calculated by one-way ANOVA (**A-B, D-G**) or two-way ANOVA (**M, N**). *P < 0.05, **P < 0.01, ***P < 0.001, ****P < 0.0001, ns, not significant.

In our scRNA-seq, classification of all the T/NK clusters in **Fig. 2F** revealed a higher ratio of CD8^+^ T cells to Treg only in the Combo-treated tumors (**Fig. S9E-G)**. Within CD8^+^ T cells, we identified six subclusters of differential gene expression profiles (**Fig. 3H-I**). Cluster 3 and cluster 5 highly expressed cytotoxic (*Gzmb, Ifng, Prf1*) and exhaustion markers (*Havcr2, Lag3, Pdcd1*), while cluster 4 featured *Gzmk, Cd7, Ly6a, and Eomes*, markers suggesting a less differentiated cytotoxic subset (**Fig. 3I**). Remarkably, all of the cytotoxic subsets, including clusters 3, 4 and 5, were dramatically enriched in Combo-treated CD8^+^ T cells (**Fig. 3J**). This aligns with our flow cytometry data showing the expansion of both exhausted and progenitor-like cytotoxic CD8^+^ T cells in the Combo group, highlighting a synergistic effect in rejuvenating anti-tumor immunity (**Fig. 3K, Fig. S9H, Table S2**).

We next investigated whether the tumor suppression in combo therapy was attributable to the increased CD8 T cells. The efficacy of Combo therapy was completely lost in NSG (NOD.*Cg*-Prkdc^scid^ll2rg^tm1Wji^/SzJ) mice (**Fig. 3L)**, and Rag1^-/-^ mice lacking an adaptive immune system, excluding direct effects of TFF2-MSA on tumor cells as the contributor to treatment efficacy (**Fig. S9I)**. Unlike anti-NK1.1 and anti-CD4 which showed no or limited effect, anti-CD8 completely abrogated the difference in tumor growth between the Combo and vehicle groups (**Fig. 3M, Fig. S9J-K**). Together, these data revealed the critical role of CD8^+^ T cells for the synergistic efficacy of the Combo regimen.

A specific CCR7^+^ DC subset of mature immunoregulatory dendritic cells (mregDC) that frequently correlates positively with CD8^+^ immunotherapy responses, significantly increased with Combination treatments^54–57^ (**Fig. S10A-D**). Compared to monocyte-derived DC (moDC) and conventional DC (cDC), the CXCR4^+^ mregDC subset expressed higher levels of *IL12b*, co-stimulatory molecules and MHC class I genes, suggesting its enhanced interaction with T cells (**Fig. S10E-F**). Intriguingly, Combo therapy resulted in a synergistic change in gene signatures only in DCs, but not in macrophages/monocytes, favoring the contribution of DCs to the T cell response (**Fig. S10G-I**). We reasoned that the cooperation between IL12^+^ DC, IFNγ^+^ CD8^+^ Τ cells, and CXCL10^+^ neutrophils underlies the synergistic efficacy of TFF2-MSA and anti-PD-1 combination^8,58^. This was supported by two observations: First, the expression levels of *Ifng, Cxcl10* and *Il12b* were all significantly upregulated in combo-treated tumors (**Fig. 2L, Fig. S9H, S10C-D**). Secondly, neutralizing antibodies against anti-IFNγ, anti-IL12 or anti-CXCL10 reversed the sensitivity of ACKP tumors to the combo therapy (**Fig. 3N**). These findings indicate that the combination therapy engaged multiple components of innate and adaptive immunity, thereby unleashing the full-fledged activity of CD8^+^ T cells.

### TFF2-MSA systemically restrains cancer-induced PMN-MDSCs and granulopoiesis

The lack of direct polarization effect of TFF2-MSA on tumor PMNs, along with mitigated splenomegaly in TFF2-MSA treated mice, suggests that TFF2-MSA may restrain PMN-MDSCs early in their generation or differentiation (**Fig. S6I, S1C, K**). We examined the blood and spleen of Hdc-GFP mice with ACKP subcutaneous tumors for the effects of TFF2-MSA, and observed a selective reduction of Hdc^+^ PMN-MDSCs, resembling their decreased accumulation in the TME (**Fig. 4A, D, Fig. S11A-B**). Selective reduction of Hdc^+^ PMN-MDSCs in circulation was similarly observed in the autochthonous mouse GC model (**Fig. S11C-E**). Following TFF2-MSA treatment, CD3^+^ and CD8^+^ T cells were more abundant in blood and spleen, consistent with the T cell influx seen in TFF2-MSA-treated tumors, indicating a systemic anti-tumor immune response (**Fig. 4B-C, E-F**).

**Fig. 4.**
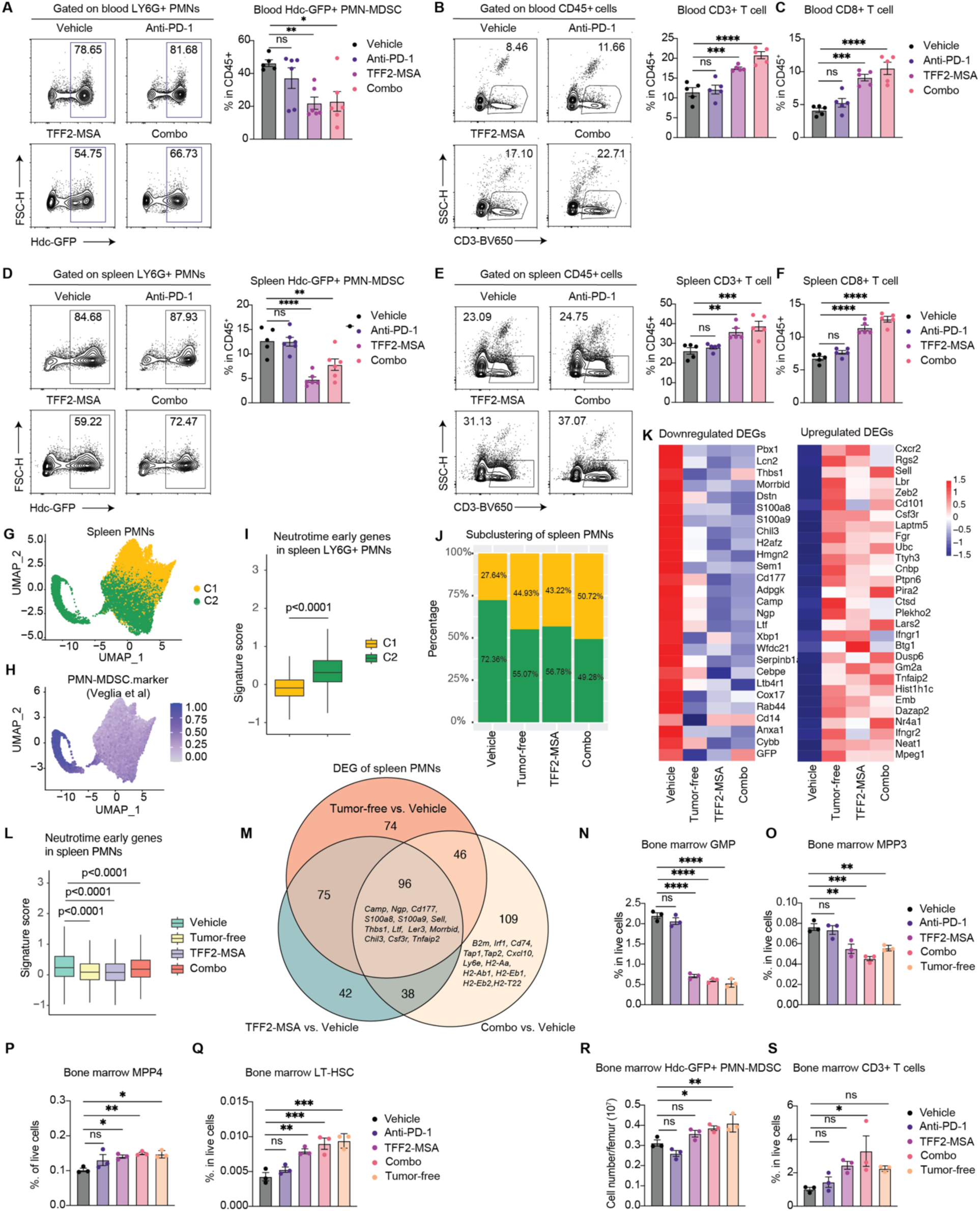
TFF2-MSA systemically restrains cancer-induced PMN-MDSCs and granulopoiesis. **A.** Representative flow cytometry plots of Hdc-GFP^+^ PMN-MDSCs within LY6G^+^ PMNs, and their quantification in the blood of ACKP model following treatments (n=5-6). **B.** Representative flow cytometry plots and quantification of CD3^+^ T cells in the blood of ACKP model following treatments (n=5). **C**. Quantification of CD8^+^ T cells in the blood of ACKP model following treatments (n=5). **D.** Representative flow cytometry plots of Hdc-GFP^+^ PMN-MDSCs within LY6G^+^ PMNs, and their quantification in the spleen of ACKP model following treatments (n=5-6). **E**. Representative flow cytometry plots and quantification of CD3^+^ T cells in the spleen of ACKP model following treatments (n=5). **F.** Quantification of CD8^+^ T cells in the spleen of ACKP model following treatments (n=5 per group). **G**. UMAP dimensional reduction depicting subclusters of the LY6G^+^ spleen PMNs isolated from tumor-free mice and ACKP tumor-bearing Hdc-GFP mice following treatments (pooled from n=3 per group). **H**. UMAP depicting the distribution of PMN-MDSC marker expressions from Veglia *et al.*^5^ in spleen PMNs. **I.** Expression of neutrotime early genes^59^ in spleen PMN subclusters. **J.** Relative frequency of spleen PMN subclusters following treatments. **K.** Heatmap of manually selected downregulated and upregulated genes in spleen PMNs from the treatment groups. DEG, differentially expressed genes. **L.** Expression of neutrotime early genes in spleen PMNs isolated from the treatment groups. **M.** Venn diagram depicting differentially expressed genes (DEGs) in spleen PMNs in the indicated group versus vehicle-treated ACKP tumor-bearing mice. Numbers in the circles indicate the number of DEGs. **N.** Quantification of GMPs in the bone marrow of tumor-free mice and ACKP tumor-bearing mice following treatments (n=3). **O, P, Q.** Quantification of MPP3, MPP4, and LT-HSC in the bone marrow of tumor-free mice and ACKP tumor-bearing mice following treatments (n=3). MPP3, multipotent progenitor 3. LT-HSC, long-term hematopoietic stem cell. **R.** Quantification of cell number of Hdc-GFP^+^ PMN-MDSCs in the bone marrow of tumor-free mice and ACKP tumor-bearing mice following treatments (n=3). **S.** Quantification of CD3^+^ T cell in the bone marrow of tumor-free mice and ACKP tumor-bearing mice following treatments (n=3). Data is presented as mean ± SEM, and P values are calculated by one-way ANOVA (**A-F, N-O, Q-S**) or Wilcoxon test (**I, L**). *P < 0.05, **P < 0.01, ***P < 0.001, ****P < 0.0001, ns, not significant.

The spleen is a major site for PMN-MDSC expansion and differentiation in murine cancer models^4^. Hence, LY6G^+^ PMNs from the spleen of ACKP-tumor-bearing mice treated with vehicle, TFF2-MSA, or Combo therapy were isolated for scRNA-seq analysis. An additional spleen sample from tumor-free mice was included to identify tumor-specific signatures in splenic PMNs (**Fig. 4G**). Consistent with previous studies^5^, tumor PMNs featured pathologically activated programs compared to spleen PMNs of the same ACKP tumor-bearing mice **(Fig. S12A, Table S3)**. Upon comparing splenic PMNs from different groups, vehicle-treated ACKP tumor-bearing mice exhibited an expanded C2 subset that preferentially expressed early neutrophil development (*Ngp, Camp, Lcn2, Ltf, Cd177*, BM proximity score, and Neutrotime early genes) and known PMN-MDSC markers (*S100a8, S100a9, Xbp1, Cd14*)^5,12,50,59^, compared to tumor-free mice (**Fig. 4H-J, Fig. S12B-C)**. This suggests tumor-driven accelerated mobilization of immature neutrophils which acquired some extent of immunosuppressive features. Importantly, TFF2-MSA treatment reduced the C2 subset to a percentage comparable to that of tumor-free mice, while Combo treatment resulted in an even smaller C2 subset than tumor-free mice, suggesting early inhibition of PMN-MDSC generation (**Fig. 4J**). Downregulation of PMN-MDSC markers (including GFP) and immaturity scores, along with upregulation of neutrophil maturation/differentiation genes (*Cd101, Csf3r, Cxcr2, Gm2a*, etc.), were uniformly observed in splenic PMNs of tumor-free, TFF2-MSA, and Combo-treated mice^14,60^ (**Fig. 4K-L, Fig. S12D**). Analyses of differentially upregulated genes (DEGs) showed a significant overlap between PMNs from TFF2-MSA treated mice and tumor-free mice compared to vehicle-treated mice, while Combo therapy induced additional upregulation of ISGs and antigen presentation genes in PMNs, recapitulating the emergence of immunostimulatory PMNs in the tumor (**Fig. 4M, Fig. S12E, Table S4**). This suggests possible early interactions of proinflammatory PMNs and T cells in the spleen. Together, these data indicate that TFF2-MSA systemically restrained tumor-driven PMN-MDSCs and restored effective anti-tumor immunity.

Solid tumors alter BM hematopoiesis by favoring myelopoiesis^61,62^, skewing hematopoietic stem cell (HSC) differentiation towards myeloid-biased multipotent progenitor 3 (MPP3) at the expense of lymphoid-biased MPP4 and long-term HSCs (LT-HSCs)^18^. MPP3 then generates myeloid committed progenitors such as granulocyte-macrophage progenitors (GMPs) to produce tumor-demanded PMN-MDSCs (**Fig. S16**)^63^. Given the effect of TFF2-MSA on reducing systemically PMN-MDSCs, we postulated that TFF2-MSA may inhibit cancer-induced myelopoiesis in the BM.

TFF2-MSA alone and Combo treatment significantly reduced the GMPs to a level similar to that in tumor-free mice (**Fig. 4N, Fig. S13A-B**). This was accompanied by contraction of the myeloid-biased MPP3 and restoration of the lymphoid-biased MPP4 and LT-HSC pool, indicating normalized hematopoiesis following TFF2-MSA treatment (**Fig. 4O-Q)**. A similar reduction of myeloid progenitors was observed in the mouse model of autochthonous GCs (**Fig. S13C**). We asked whether TFF2-MSA acts on granulopoiesis directly or indirectly via our previously defined negative feedback loop, whereby BM Hdc^+^ myeloid cells restrain myelopoiesis from hematopoietic stem cells (HSCs)^18^. Flow cytometry showed that TFF2-MSA treatment restored the BM Hdc^+^ PMN number to levels observed in tumor-free mice following, along with an increased percentage of CD3^+^ T cells (**Fig. 4R-S**). On the other hand, in BM of Cxcr4-GFP reporter mice, we found the highest expression of CXCR4 in GMPs among stem cell and myeloid progenitors, and its expression was predominantly influenced by TFF2-MSA in GMPs, nominating GMPs as the most possible drug target (**Fig. S13D-E**). In the culture of sorted myeloid progenitors (MPs) in media designed to promote myeloid differentiation^61^, TFF2-MSA inhibited GMPs and Hdc^+^ PMN production, both in the absence and presence of SDF-1 (**Fig. S13F-G**). In summary, these data demonstrated that TFF2-MSA reduced myeloid progenitors in the BM, likely via both direct and indirect mechanisms, thus systemically normalizing PMN production.

### Superior efficacy of anti-PD-1 plus TFF2-MSA compared to combinations with anti-LY6G or CXCR4 antagonist AMD3100

Given the marked efficacy of the partial CXCR4 agonist in combination with anti-PD-1, we compared TFF2-MSA with existing PMN-targeted strategies, such as anti-LY6G and the clinically available CXCR4 antagonist, AMD3100. Despite using an optimized anti-LY6G-based neutrophil elimination strategy^13^, anti-LY6G only slightly improved tumor responses when combined with anti-PD-1 in the ACKP model (**Fig. S14A-B**). We further reasoned that deleting total PMNs with anti-LY6G might diminish the TFF2-MSA therapeutic effect if the two agents were used together. Indeed, adding anti-LY6G to TFF2-MSA plus anti-PD-1 reduced the efficacy of the Combo regimen, indicating a role for some LY6G^+^ PMNs in the anti-tumorigenic response (**Fig. S14C**). In a side-by-side comparison of TFF2-MSA to the CXCR4 antagonist AMD3100, we found that AMD3100 failed to enhance anti-PD-1 immunotherapy response in our model. Indeed, TFF2-MSA plus anti-PD-1 markedly outperformed AMD3100 plus anti-PD-1 in tumor inhibition (**Fig. S14D**). Flow cytometric analysis revealed that AMD3100 alone or its combination with anti-PD-1 failed to reduce Hdc^+^ tumor PMN-MDSCs in the TME and the blood, and bone marrow GMPs of tumor-bearing mice (**Fig. S14E-H**). This aligns with reports that AMD3100 induces an accumulation of immature neutrophils in the TME^16^, which is likely due to its mobilization of BM neutrophils leading to compensatory granulopoiesis. Consistent with this notion, automated white blood cell (WBC) counts also revealed a distinct mode of action of TFF2-MSA compared to AMD3100 in tumor-free mice (**Fig. S14I**). Notably, TFF2-MSA did not profoundly elevate the neutrophil percentage in WBC within 6 hours post-administration, unlike AMD3100 (**Fig. S14J**), suggesting the superiority of TFF2-MSA in maintaining BM homeostasis, likely due to its milder effect on desensitizing CXCR4 (**Fig. S7E**).

### TFF2 is reduced and negatively correlates with elevated CXCR4^+^LOX-1^+^ PMN-MDSCs in GC patients

In patients with cancer, low-density PMN-MDSCs are found among the peripheral blood mononuclear cells (PBMCs) in circulation, unlike the conventional mature high-density neutrophils (HDN)^12^. However, some mature neutrophils undergoing phenotypic changes may also accumulate in the PBMC layer, complicating the phenotyping of PMN-MDSCs^64^. To better characterize PMN-MDSCs, we obtained fresh PBMCs from untreated GC and gastroesophageal junction cancer (GEJ) patients for flow cytometric characterizations (**Table S5**). Compared with age-matched healthy donors, both GC and GEJ patients showed significantly elevated levels of total MDSCs (CD45^+^Lin^-^HLA-DR^-^CD11b^+^CD33^+^) in PBMCs (**Fig. 5A**). We further categorized total MDSCs into three subsets: granulocytic MDSC (PMN-MDSC, CD45^+^Lin^-^HLA-DR^-^ CD11b^+^CD33^+^CD14^-^CD66b^+^), monocytic MDSC (M-MDSC, CD45^+^Lin^-^HLA-DR^-^ CD11b^+^CD33^+^CD14^+^CD66b^-^), and early MDSC (eMDSC, CD45^+^Lin^-^HLA-DR^-^CD11b^+^CD33^+^CD14^-^CD66b^-^)^65^ (**Fig. S15A**). Among all three subsets, we found the greatest expansion of PMN-MDSC, in GC and GEJ patients by roughly 9.5 fold and 44.3 fold respectively compared to healthy donors (**Fig. 5B, Fig. S15B**). The identified PMN-MDSCs expressed high levels of CD15 and moderate levels of CD33 as reported^65^ (**Fig. S15C-D**). Deletion of this arginase 1^+^ PMN-MDSC fraction from PBMCs by CD66b-specific beads restored autologous T cell proliferation *in vitro* (**Fig. S15E-F**), further confirming their identity. Lectin-type oxidized LDL receptor 1 (LOX-1), a specific marker for human immature PMN-MDSCs^66^, showed exclusive expression in low-density PMN-MDSCs, rather than HDNs (**Fig. S15G**). LOX-1^+^ cell frequency in PMN-MDSCs showed a trend of increase in gastric and GEJ cancer patients (**Fig. S15H**). When we included the neutrophil maturation marker CD16 in our analysis^67^, the predominant phenotype of PMN-MDSCs switched from CD16^+^LOX-1^-^ to CD16^-^ LOX-1^+^ in GC and GEJ patients (**Fig. S15I-J**), a finding that we confirmed with another mature neutrophil marker CD10 (data not shown)^67^. These data indicate a profound expansion in GC and GEJ patients of immature PMN-MDSC.

**Fig. 5.**
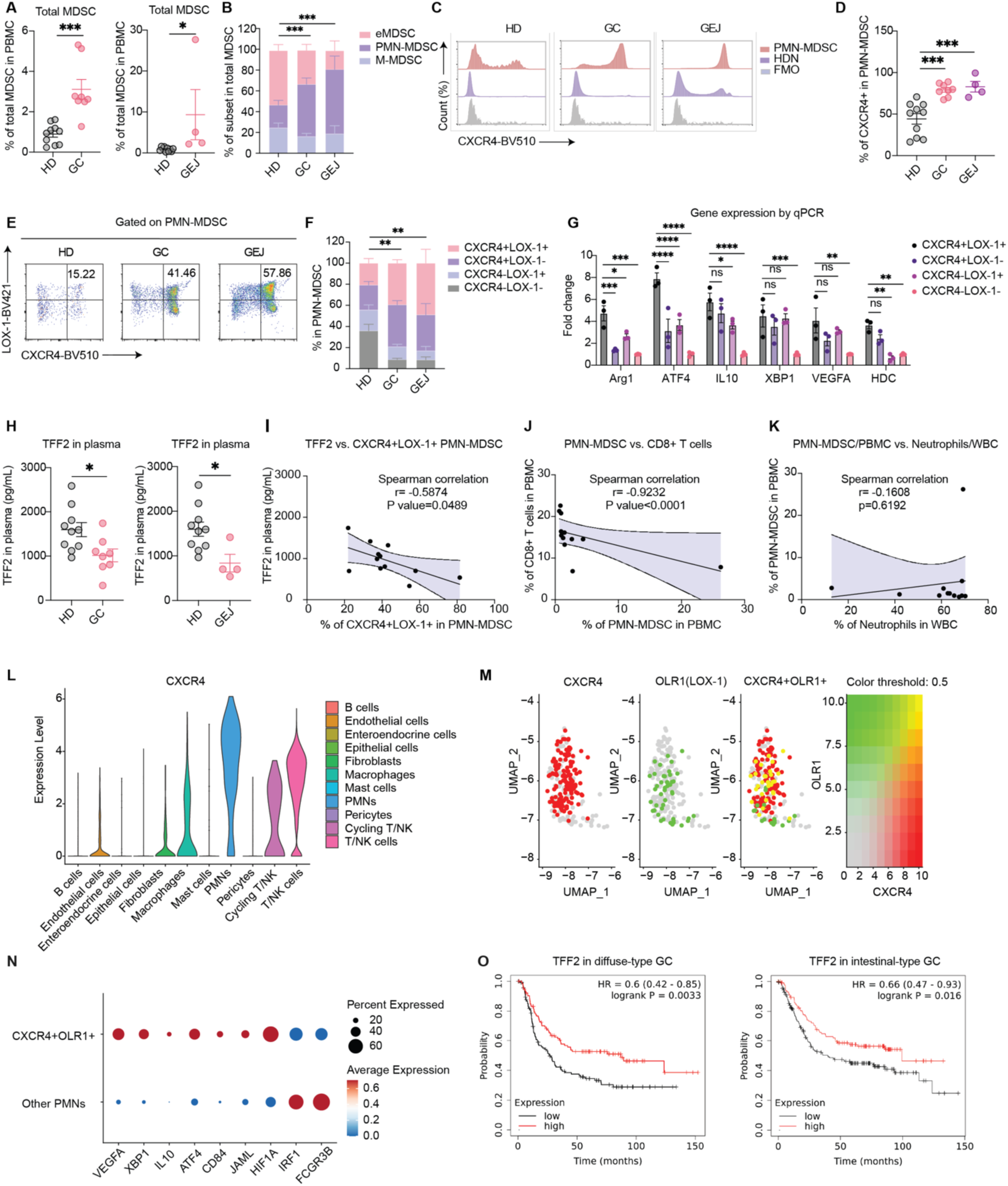
TFF2 is reduced and negatively correlates with elevated CXCR4^+^LOX-1^+^ PMN-MDSCs in GC patients. **A.** Total MDSCs in PBMCs of HD, GC and GEJ patients. HD (healthy donor), n=10. GC, n=8. GEJ (gastroesophageal junction cancer), n=4. **B.** Relative frequency of the three MDSC subsets within total MDSCs in HD, GC and GEJ patients. Statistical significance was calculated for PMN-MDSC percentage. **C.** Representative histogram of CXCR4 expression on PMN-MDSCs and HDN shown as the normalized count percentage of the total population. HDN, high density neutrophil. FMO, fluorescence minus one control. **D.** Percentage of CXCR4^+^ cells in PMN-MDSCs in HD, GC, and GEJ patients (n=10, 8, 4). **E.** Representative flow cytometry plots of CXCR4^+^ LOX-1^+^ PMN-MDSCs in HD, GC, and GEJ patients. **F.** Relative frequency of subsets defined by CXCR4 and LOX-1 within PMN-MDSCs of HD, GC and GEJ patients (n=10, 8, 4). Statistical significance was calculated for CXCR4^+^LOX-1^+^ PMN-MDSCs. **G.** Gene expressions of known PMN-MDSC markers (*ARG1, ATF4, IL10, XBP1, VEGFA, HDC*) in the PMN-MDSC subsets from GC patients (n=3) detected by RT-qPCR. **H.** TFF2 serum levels in the blood of HD, GC, and GEJ patients (n=10, 8, 4). **I.** Correlation of TFF2 serum levels with CXCR4^+^LOX-1^+^ subset frequency within PMN-MDSCs of GC and GEJ patients (n=8, 4). **J.** Correlation of CD8^+^ T cell with PMN-MDSC frequency in PBMCs of GC and GEJ patients (n=8, 4). **K**. Correlation of neutrophils percentage in WBCs with PMN-MDSC percentage in human GC and GEJ PBMCs (n=8, 4). **L.** CXCR4 expression in different cell types of human gastric cancer tissues by scRNA-seq analysis (n=2). **M.** Overlap of *CXCR4* and *OLR1*(gene name of LOX-1) within PMNs of human gastric cancer tissues (n=2). *CXCR4^+^OLR1^+^* PMNs are highlighted in yellow. **N.** Pro-tumoral and anti-tumoral markers in *CXCR4^+^OLR1^+^*and the remaining portion of PMNs in human gastric cancer tissues (n=2). **O.** Kaplan-Meier curves of overall survival in human diffuse-type (n=241) and intestinal-type GC patients (n=320) according to *TFF2* median values. Data is presented as mean ± SEM, and P values are calculated by one-way ANOVA (**B, D, F, G**), two-sided unpaired Student’s t-test (**A, H**), or log-rank test (**O**). For correlation analysis, The Spearman correlation coefficient was calculated and indicated (**I, J**). *P < 0.05, **P < 0.01, ***P < 0.001, ****P < 0.0001, ns, not significant.

CXCR4 expression was restricted to low-density PMN-MDSCs rather than HDNs in all blood samples of HD, GC, and GEJ (**Fig. 5C, Fig. S15K**). Notably, a significant CXCR4^+^LOX-1^+^ subset predominated in PMN-MDSCs from GC and GEJ patients (**Fig. 5D-F**). This subpopulation exhibited the highest levels among PMN-MDSCs of many immunosuppressive gene transcripts (*ARG1, ATF4, IL10, XBP1, VEGFA*) and *HDC*, as well as Arginase 1 at the protein level, underscoring its immunosuppressive feature (**Fig. 5G, Fig. S15L**). Additionally, TFF2 level in human plasma decreased in GC and GEJ patients compared to healthy controls (**Fig. 5H**), and it inversely correlated with both PMN-MDSC frequency in PBMCs, and the CXCR4^+^LOX-1^+^ percentage within PMN-MDSCs (**Fig. 5I, Fig. S15M**). These observations support the notion that TFF2 may be an important negative regulator of PMN-MDSCs in human GC. In keeping with their role in suppressing cytotoxic T cells, circulating levels of PMN-MDSC inversely correlated with total CD8^+^ T cell and granzyme B^+^ CD8 T cell numbers (**Fig. 5J, Fig. S15N**).

Interestingly, neutrophil counts from complete blood count (CBC) with differential in patient blood display no correlation with PMN-MDSC abundance, suggesting that routine clinical neutrophil counts may not adequately reflect PMN-MDSC levels (**Fig. 5K**). Indeed, we noticed a trend of elevated PMN-MDSC and its CXCR4^+^LOX-1^+^ subset in GC patients with distant metastasis compared to those without metastasis, while this trend was not seen with neutrophil counts or their frequency in WBCs (**Fig. S15O**).

Lastly, we reanalyzed our previously published scRNA-seq data including 1 diffuse-type and 1 intestinal-type human GC sample (**Table S6**)^68^. Despite the limited PMN number identified due to their low RNA content (**Fig. S16A-B**), reclassification showed the highest expression of CXCR4 in PMNs among all cell types in human GC tissues (**Fig. 5L**). A CXCR4^+^LOX-1^+^ subset was detected within tumor PMNs, enriching for immunosuppressive markers including *VEGFA, XBP1, IL10, ATF4, CD84, JAML, and HIF1A* (**Fig. 5M-N**). Stratification of patients with diffuse-type or intestinal-type GC, based on median TFF2 levels in GC tissues pooled from six published Gene Expression Omnibus (GEO) datasets, revealed that low *TFF2* expression was significantly associated with poor overall survival in both groups (**Fig. 5O**). These human data underscore our mouse model’s fidelity and our studies’ translational potential.

## Discussion

Accumulating evidence has positioned the interaction between PMN-MDSCs and CD8 T cells at the core of the immunosuppressive TME^69^. While previous efforts aimed at blocking or depleting PMN-MDSCs^2,7^, major challenges have impeded the development of effective PMN-MDSC-targeted therapies: First, most of these preclinical studies rely merely on the granulocyte marker LY6G for PMN-MDSC phenotyping which also identifies pro-inflammatory or anti-tumorigenic neutrophils^3,65^. Secondly, their compensatory overproduction from cancer-expanded myeloid progenitors has rendered most treatment effects transient and limited^13–16^. Optimal treatment strategies appear to require systemic targeting of PMN-MDSC production, differentiation, and recruitment to cancer sites. Our novel CXCR4 partial agonist, TFF2-MSA peptide, showed remarkable synergy in GC treatment with anti-PD-1, as the combination led to tumor regression or eradication, significant metastasis reduction, and mouse survival extension. By using Hdc-GFP to identify bona fide tumor PMN-MDSCs, we found TFF2-MSA selectively reduced Hdc-GFP^+^ PMN-MDSCs among the LY6G^+^ population along their development trajectory in the spleen, blood, and tumor. Furthermore, TFF2-MSA directly restrained cancer-induced granulopoiesis by reducing the expansion of myeloid progenitors (e.g. GMP). Consequently, combination therapy switched the TME from immunosuppressive to immunostimulatory, leading to profound activation of an anti-tumor cytotoxic T cell response. Our findings thus support a new combination strategy, using CXCR4 partial agonism to overcome anti-PD-1 resistance in GC and other PMN-MDSC enriched GI cancers (**Fig. S17**). Notably, in our aggressive GC model, only this regimen demonstrated efficacy, whereas combinations of chemo-immunotherapy that were effective against early-stage GC failed^40^. Monotherapy with TFF2-MSA demonstrated better control of metastatic lesions over primary cancers, underscoring the significance of CXCR4 signaling in advanced stages of GC^1,26^.

TFF2-MSA exhibited clear advantages over other PMN-MDSC targeted therapies in that it selectively targeted PMN-MDSCs while preserving the immunostimulatory PMNs. While the role of PMNs in mediating immunotherapy resistance is well established, only recently have studies identified an anti-tumoral PMN subset that potentiates immunotherapy response, featured by interferon response^8,12^, antigen presentation functions^9,11^, or cytotoxicity^8,10^. These reports have highlighted the need to discriminate carefully between pro- and anti-tumorigenic PMN subtypes when designing combinatorial immunotherapies. Here, we utilized the Hdc-GFP model to track and isolate the immunosuppressive PMN-MDSCs^17–19^. which was validated by our scRNA-seq analysis. Unbiased clustering revealed two tumor PMN subsets, C1 and C2, with distinct morphology, transcriptomics, and function in our scRNA-seq analysis. The C2 subset was mostly Hdc-GFP^+^ and CXCR4^high^, and represented the T-cell inhibiting PMN-MDSCs, and was selectively reduced by TFF2-MSA in tumor, blood, and spleen. Our trajectory analysis as well as phenotyping of the cultured subpopulations, supported that the subsets have undergone imprinting before reaching the tumor. The selective action of TFF2-MSA on Hdc-GFP^+^ PMN-MDSCs with high CXCR4 was further supported by our chemotaxis assay, which showed TFF2 preferential modulation of Hdc-GFP^+^ subset chemotaxis and partial blockade of SDF-1-induced downstream signaling. Given the abundant SDF-1 chemotaxis signal emanating from the TME, the CXCR4 partial agonist TFF2-MSA may disorient the Hdc-GFP^+^ PMN-MDSCs away from the tumor-derived SDF-1 signal, redirecting them to physiological functions at their normal sites, similar to the findings that forced SDF-1 overexpression in cancer cell itself impede cancer cell metastasis^33^. Consistent with this notion, we found TFF2-MSA normalized the abundance of Hdc^+^ PMN-MDSCs in the blood, spleen, and especially bone marrow to levels similar to those in the tumor-free mice.

CXCR4 is best known for its role as a master regulator of hematopoietic homeostasis in the bone marrow^1^, as evidenced by the impaired hematopoiesis and embryonic lethality in mice lacking either CXCR4 or SDF-1^70^. The SDF-1-CXCR4 axis was initially identified as the key retention signal for hematopoietic stem and progenitor cells (HSPCs) and neutrophils in the bone marrow. CXCR4 is the only chemokine receptor with a gain-of-function mutation that leads to a human hereditary disorder as its impaired desensitization causes circulating neutropenia^70^. Thus, CXCR4 antagonist plerixafor (AMD3100) is used clinically for stem cell mobilization and transplantation. Subsequent studies have expanded the role of CXCR4 in hematopoiesis, including its promotion of HSPC survival and quiescence, and myelopoiesis^1,70,71,72^. CXCR4 also plays numerous roles in the immune system, including contributing to the development and homing of T cells^73,74^, DCs^27,75^, B cells^76^, and the formation of immune synapses ^28^. However, in various cancers including GC^77^, SDF-1-CXCR4 signaling is hyperactivated in the TME, contributing to the immunosuppression by recruiting MDSCs and Tregs and repelling T cells^29^. Although it was predicted that CXCR4 inhibition could potentiate immunotherapy, clinical trials of CXCR4 antagonists have generally shown modest efficacy^26,29^. Reminiscent of the granulocytosis of immature neutrophils observed in CXCR4^-/-^ mice, a recent phase 2 trial of plerixafor (AMD3100) revealed it caused premature release of immature neutrophils from bone marrow and their abnormal accumulation in the TME, compromising combinatorial efficacy with anti-PD-1^16^. This suggested the need for fine-tuning CXCR4 signaling to preserve some of its physiological functions^78,79^. Indeed, our study demonstrated that CXCR4 partial agonism, achieved with TFF2-MSA, achieved superior tumor control compared to anti-LY6G or CXCR4 antagonist AMD3100 when combined with anti-PD-1. Unlike these therapeutics, TFF2-MSA did not result in a rebound of myeloid production or robust mobilization of neutrophils^13,16,80^. Instead, TFF2-MSA reduced overactivated myeloid-based MPP3 and myeloid-committed GMPs, while restoring LT-HSCs and lymphoid-biased MPP4 in tumor-bearing mice. This myeloid/lymphoid rebalancing may be due to the restored retention of Hdc^+^ PMNs in the marrow, which suppresses HSPC myeloid bias through an established paracrine feedback loop^18^. Furthermore, TFF2-MSA directly inhibited GMPs and the overproduction of Hdc^+^ PMNs. Nevertheless, given the broad hematopoietic expression patterns of CXCR4, TFF2-MSA likely influences granulopoiesis at multiple levels^71^. Consistent with these findings, studies showed that CXCR4 loss on neutrophils alone, or its heterozygous deletion, was sufficient to reduce tumor burden^81,82^, suggesting that a more refined CXCR4-targeting approach, such as partial agonism, could be a viable alternative as a cancer treatment.

Intratumoral CD8^+^ T cells increased during TFF2-MSA monotherapy, likely due to an influx from circulation rather than local proliferation, as their numbers also rose in circulation, spleen and bone marrow. This increase may be attributed to systemic relief of T cell inhibition by PMN-MDSCs, or a rebalancing of the myeloid-to-lymphoid ratio in hematopoiesis. However, emergence of cytotoxic subsets among CD8^+^ T cells and significant tumor regression were observed only with the combination regimen, when innate and adaptive immunity were co-targeted, consistent with other studies^8,11,83^. The combo therapy appeared to simultaneously induce a CCR7^+^ DC subset, which potentially possesses antigen-presenting functions^56,57^ and cooperated with CD8^+^ T cells and neutrophils in mediating its efficacy. Whether TFF2-MSA directly affects DCs or other immune cells remains to be investigated.

Lastly, our patient analysis supported our findings in mouse models. In the blood of GC and GEJ patients, we observed a deficiency of TFF2 plasma levels, accompanied by an elevation of the lower density PMN-MDSCs in blood PBMCs, which functionally inhibited T cell proliferation and negatively correlated with CD8^+^ T cell numbers. Among PMN-MDSCs, a prominent CXCR4^+^ LOX-1^+^ subset showed the greatest expansion with the highest levels of immunosuppressive genes in GC and GEJ. Lower TFF2 plasma levels correlated with higher numbers of PMN-MDSCs and a larger fraction of CXCR4^+^LOX-1^+^ subpopulation, supporting a model where TFF2 serves as a negative regulator of PMN-MDSCs. In the GC tumor samples, PMNs harbored the highest CXCR4 expression similar to our observations in mice, and low levels of TFF2 in GC tissues predicted poor survival of patients. These findings corroborated previous observations suggesting that TFF2 may restrain gastric tumor progression in humans^37^. Notably, the frequency of PMN-MDSC is not reflected by routine neutrophil counts in our small GC cohort, raising the question of whether the CXCR4^+^LOX-1^+^ PMN-MDSCs or TFF2 level might serve as better predictors of clinical resistance to immunotherapy. Studies have demonstrated not only a significant accumulation of the CXCR-high tumor PMNs but also CXCR4-high granulocytic-biased hematopoietic progenitors in human tumors, further supporting the utility of CXCR4 modulators in reducing the source of human PMN-MDSCs^84^. Collectively, our data demonstrate that CXCR4 partial agonism by restoring TFF2 may represent an attractive approach to overcome resistance to anti-PD-1 immunotherapy, in the treatment of GC and other tumors.

## Acknowledgments

This research was supported by the NIH/NCI Cancer Center Support Grant P30CA013696; grants from the NIH/NCI, including the NCI Outstanding Investigator Award R35CA210088, and funding from Tonix Pharmaceuticals (Tonix CU21-3958) to T.C.W.; the NCI Outstanding Investigator Award (R35 CA197745) to A.C. and T32 support (T32DK007647) to Q.T.W. Both T.C.W. and R.A.W. received support from the Department of Defense W81XWH-21-10901 grant. This study was supported by the NIH/NIDDK Columbia University Digestive and Liver Disease Research Center grant P30DK132710 and used their BioImaging, BioInformatics, Organoid and Clinical Biospecimen Cores. This publication was supported by the National Center for Advancing Translational Sciences, National Institutes of Health, through Grant Number UL1TR001873. This work also used Cancer Stem Cell Initiative Flow Core Facility, Molecular Pathology/MPSR, Columbia Genome Center Single Cell Analysis Core, Oncology Precision Therapeutics and Imaging Core, Columbia Center for Translational Immunology, Genomics and High Throughput Screening, and Columbia Database Shared Resource. We thank Dr. Sandra Ryeom (Columbia University Irving Medical Center) and her mentee Dr. Chi-Lee Ho for providing ACKP cell line; Dr. Adam Bass (Novartis Institute for Biomedical Research) and his team Dr. Sho Hangai and Varun Sahu for providing PC organoids; Dr. Michael Kissner and Cancer Stem Cell Initiative Flow Core Facility for expertise and help during this project; Erin Bush for her expertise in scRNA-seq. We deeply appreciate the patients and donors who participated in this study, and Emma K. Blystone, and Dr. Joel T. Gabre and other clinician and nurses from Columbia Database Shared Resource (DBSR) and Adriana Prada Rey from Columbia Center for Translational Immunology (CCTI) for collecting the clinical samples.

## Author contributions

JQ and TCW conceived, designed the study, and wrote and edited the manuscript. TCW supervised the study and secured the funding. JQ performed most of the experiments and analysis and assisted with the interpretation of scRNA-seq analysis. CM performed the scRNA-seq analysis of PMNs and DCs from tumors, and splenic PMNs, and contributed to data interpretation. QW performed the bone marrow sampling, contributed to bone marrow analysis, and the development of other experimental protocols. XZ performed the orthotopic cell injection and contributed to the development of liver and lung metastasis models. CSM performed the scRNA-seq analysis of T cells and data interpretation and assisted with PMN scRNA-seq analysis. RT performed the scRNA-seq analysis of human GC samples. HK designed the vector, contributed to the protocol for HiBiT experiment, and performed the independent histopathological analysis. FW, BZ, YZ, HZ, YO, RAW, and DWH assisted with experimental protocols or participated in project discussions. LBZ and JSL maintained mouse colonies. LZM assisted with the scRNA-seq data visualization. RHM obtained clinical samples. AH supervised the scRNA-seq analysis of T cells. BD and SL provided the TFF2 peptides and performed the experiments of all colorectal and pancreatic cancer mouse models. All authors reviewed the manuscript before submission.

## STAR Methods

### Key resources table

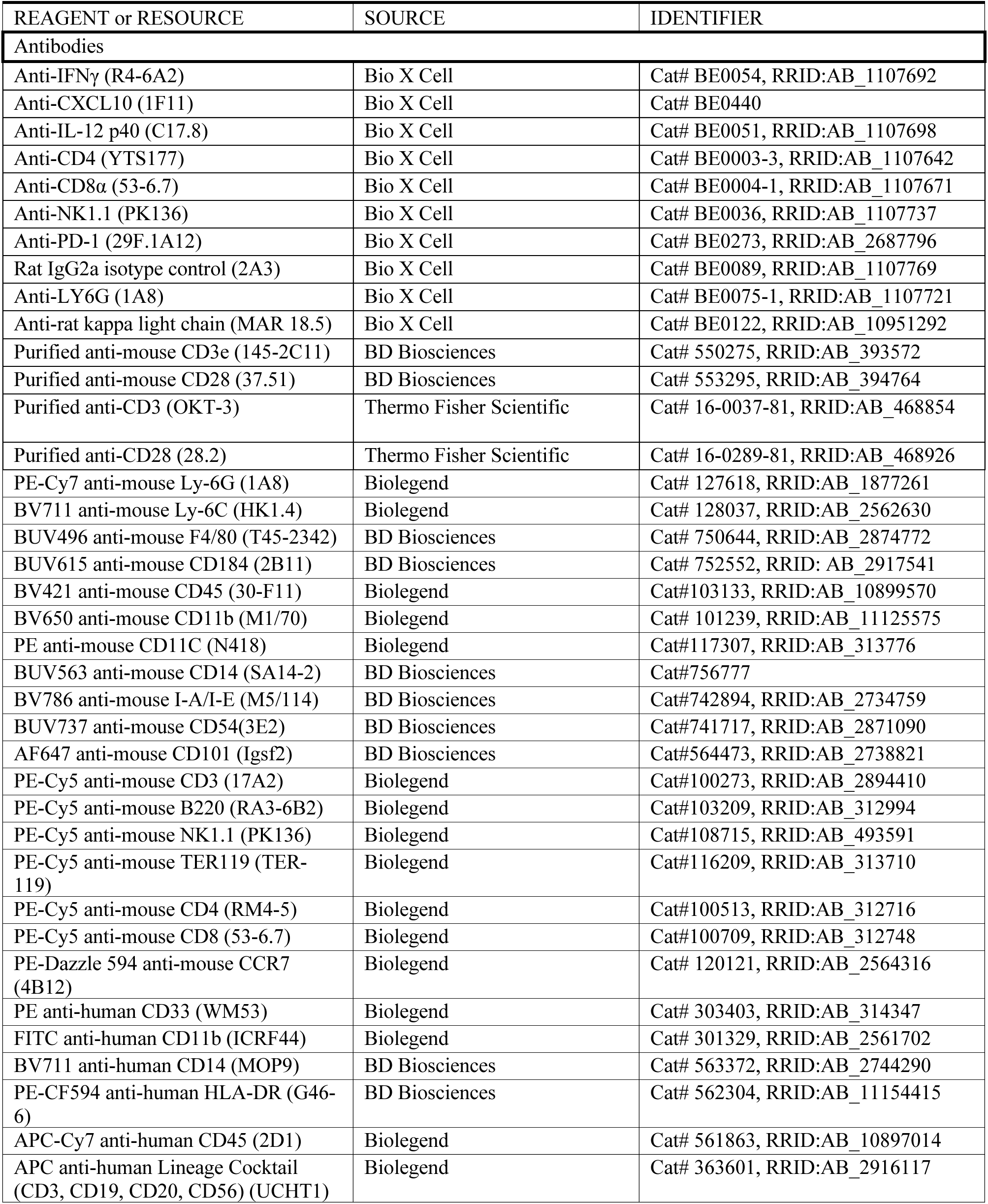
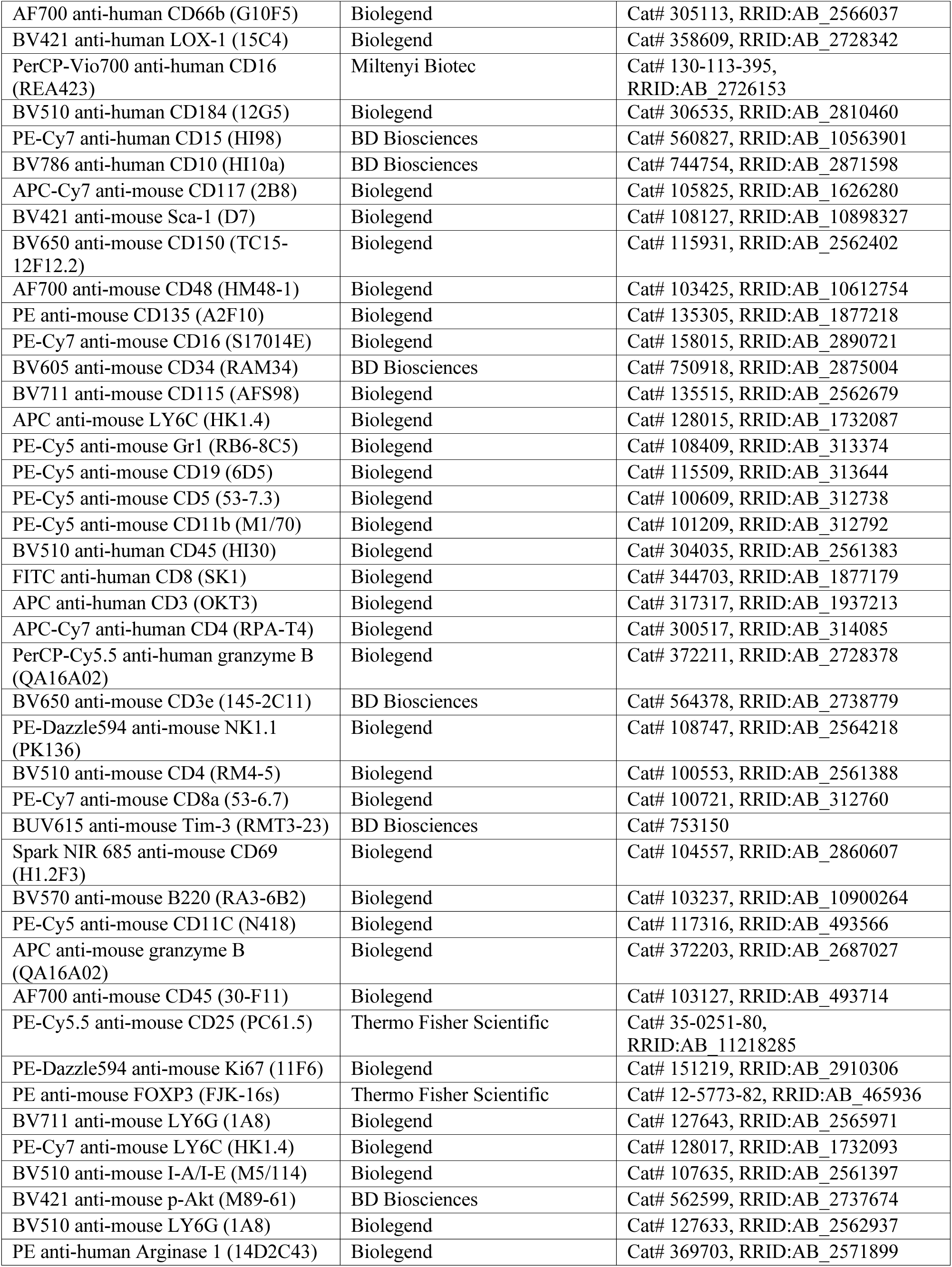
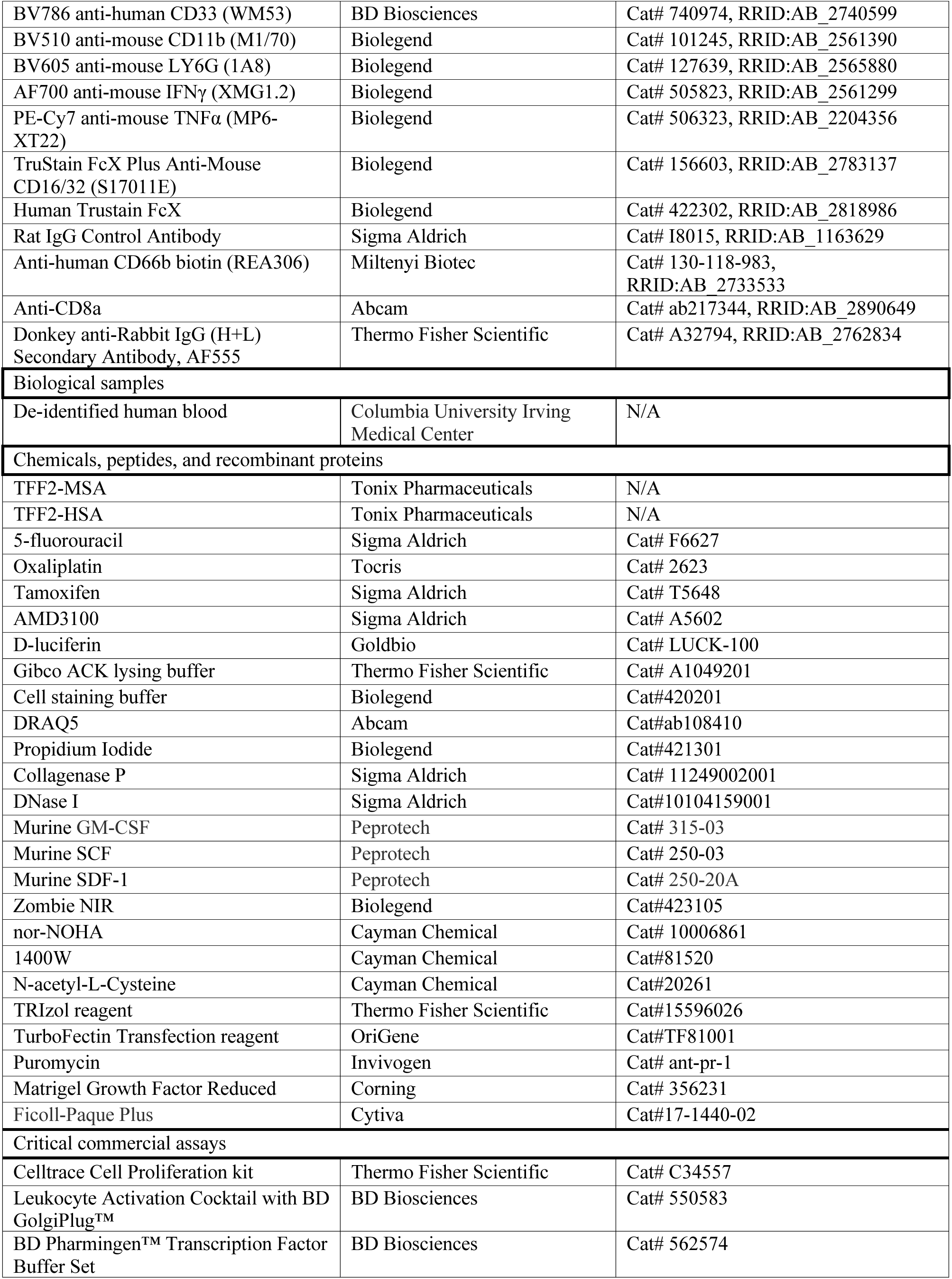
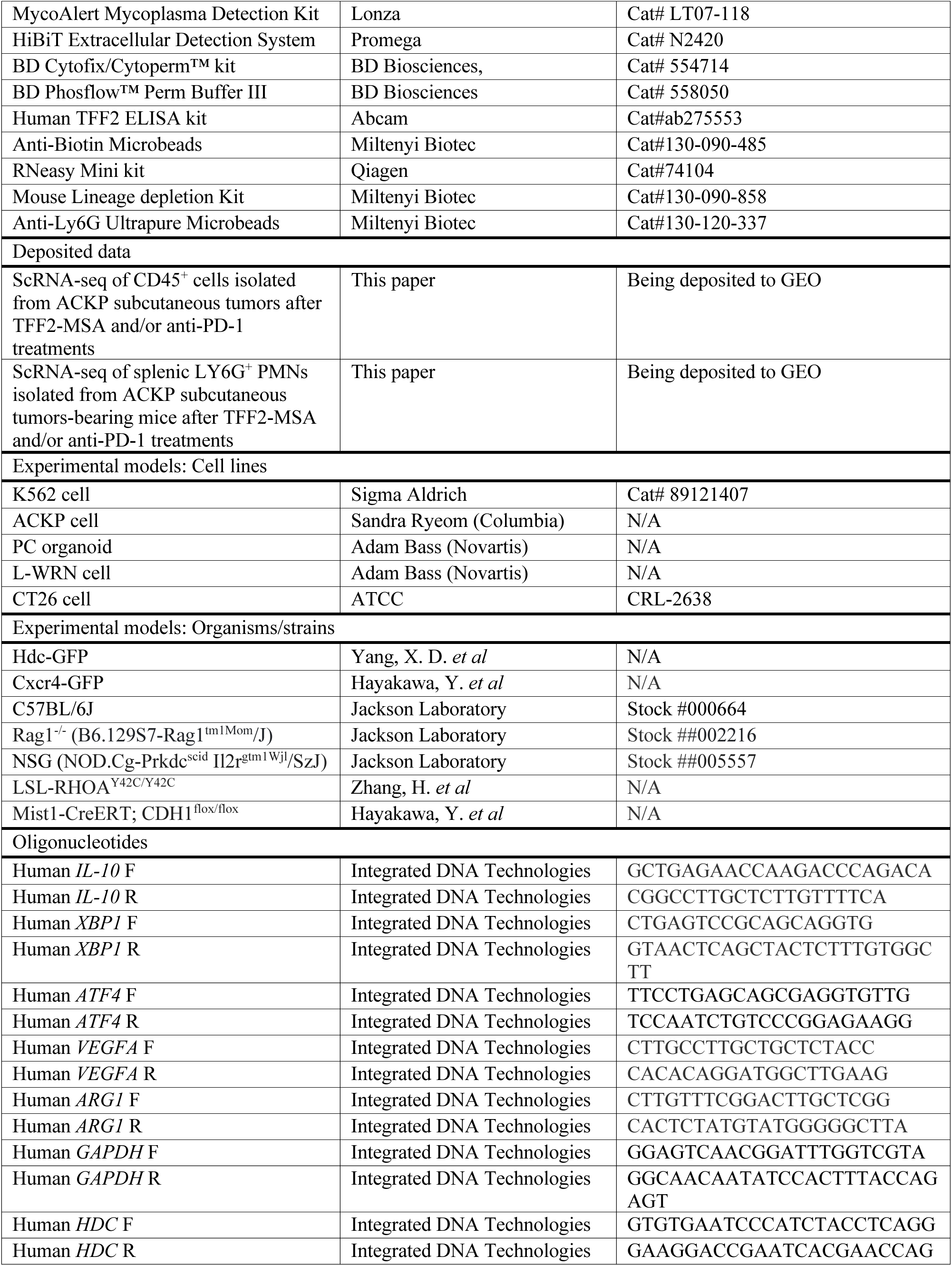
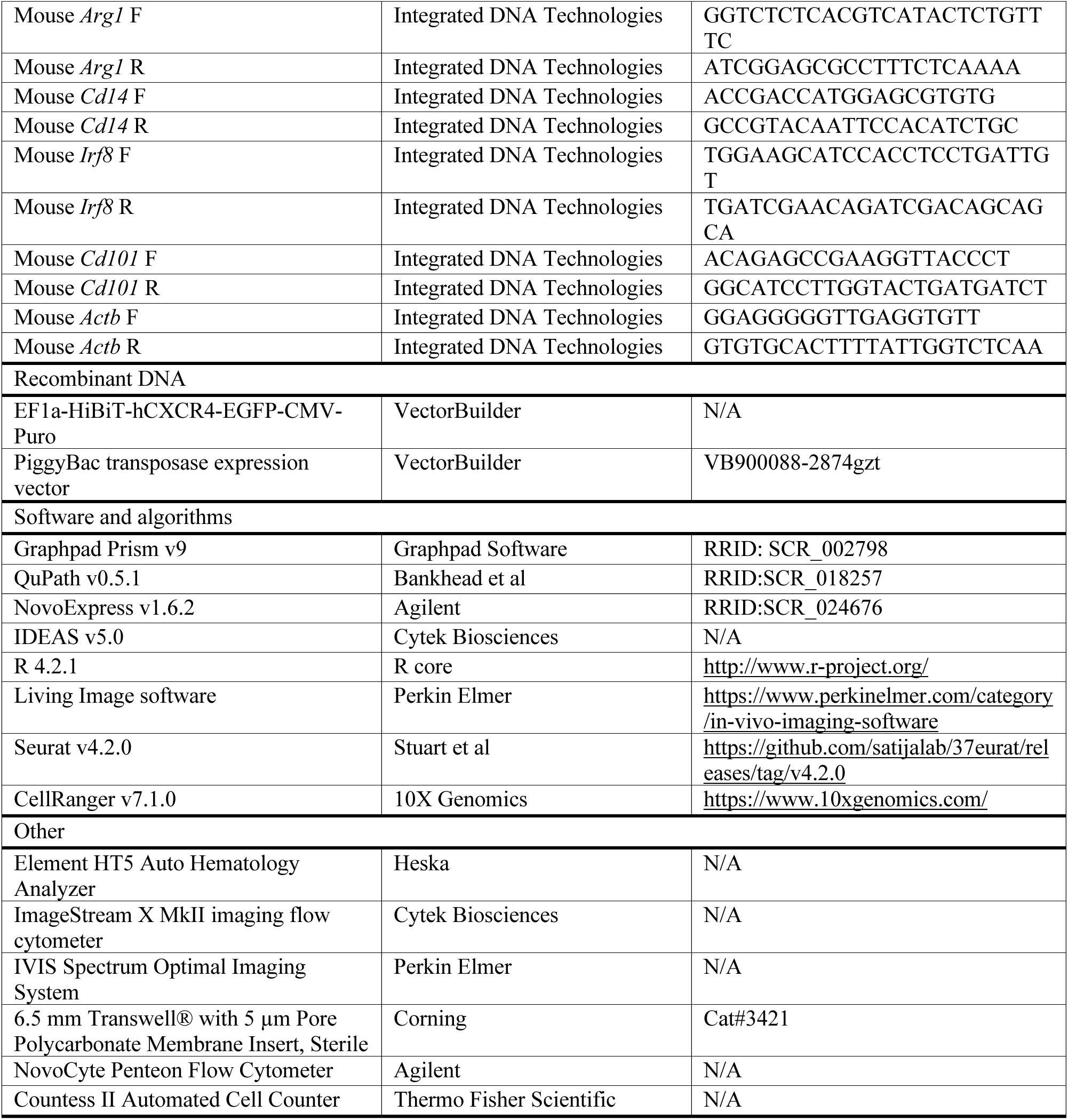

### Resource availability

#### Lead contact

Requests for further information and resources should be directed to and will be fulfilled by the lead contact, Timothy C. Wang.

#### Materials availability

All unique/stable reagents generated in this study are available from the lead contact without restriction.

#### Data and code availability

Single-cell RNA-seq data is being uploaded and deposited at Gene Expression Omnibus (GEO), and will soon be publicly available. This paper does not report original code. Any additional information required to reanalyze the data reported in this paper is available from the lead contact upon request.

### Experimental model and subject details

#### Human samples

Peripheral blood from gastric adenocarcinoma patients was collected at Columbia University Irving Medical Center, as approved by Columbia University IRB (AAAT8778). A total of 8 patients with pretreatment gastric cancer and 4 patients with pretreatment GEJ cancer were enrolled. Peripheral blood from healthy volunteers was collected with informed consent, as approved by the Columbia University IRB (AAAF0548). Patient characteristics for the blood study were obtained from Columbia Database Shared Resource (DBSR), and are detailed in Table S5. Patient characteristics reused for scRNA-seq from our previous study are detailed in Table S6.

#### Animals

All animal experiments were conducted in accordance with the National Institute of Health guidelines for animal research and approved by the Institutional Animal Care and Use Committee of Columbia University. Mice were housed in a specific pathogen-free facility. NSG (NOD.Cg-Prkdc^scid^ Il2r^gtm1Wjl^/SzJ, #005557) and Rag1^-/-^ (B6.129S7-Rag1^tm1Mom^/J, #002216) mice were purchased from the Jackson Laboratory. Hdc-GFP transgenic mice were described previously^17^. and Cxcr4-GFP mice were described previously^85^. Mist1-CreERT; CDH1^flox/flox^ mice were described previously^85^. LSL-RHOA^Y42C/Y42C^ mice were described previously^86^.

#### Mouse gastric cancer models

ACKP (Atp4b-Cre;Cdh1^-/-^;LSL-Kras^G12D/+^;Trp53^-/-^) gastric cancer cell line, previously described as Tcon cell line, was developed by Dr. Sandra Ryeom, Columbia University^41^. PC (Trp53^-/-^; CCNE1) gastric cancer organoid line was developed by Dr. Adam Bass (Novartis Institute for Biomedical Research). Briefly, normal gastric organoids were isolated from Trp53^flox/flox^ mice, and then Adenovirus-Cre was transduced to the organoids *in vitro* to knock out the Trp53 gene. Nutlin-3a treatment was performed to select Trp53-deficient cells. Then, the organoids were transduced with a lentivirus CCNE1 vector to overexpress CCNE1. P53 deficiency and CCNE1 overexpression were validated in the organoids by western blotting before they were used for *in vivo* experiments.

6-8 week-old male Hdc-GFP mice, C57BL/6J mice or Cxcr4-GFP mice were implanted subcutaneously with 1.5×10^6^ ACKP or 2 x10^6^ PC cells in 50% Matrigel solution (Corning) with PBS. After 10 days when the tumors reached an average volume of 150-250 mm^3^, these mice were randomized into each group with the same average tumor volume, and subjected to respective treatments (see Fig. S1B). Anti-PD-1 (10 mg/kg, Clone 29F.1A12, Bioxcell, Cat# BE0273) and TFF2-MSA (22.5 mg/kg, Tonix Pharmaceuticals) were intraperitoneally injected every 3 days to the indicated groups. For comparison, the vehicle group was injected with an isotype control antibody (10 mg/kg, Clone 2A3, Bioxcell). For treatment groups involving chemotherapy, 30 mg/kg 5-fluorouracil plus 5mg/kg oxaliplatin was given intraperitoneally every 3 days. Tumor size was monitored using digital calipers on the same day of the treatments. Tumor volume was calculated using the formula: volume =1/2 (Width^2^ × Length).

For the orthotopic tumor model, luciferase-expressing ACKP cells were used. Mice were anesthetized with 2–4% isoflurane and placed on a heat pad. A middle incision was performed, and the anterior wall of stomach was exposed. ACKP cells (1.5×10^6^ cells in 20 μL PBS) were drawn up into a sterile 31 G × 355 5/8” syringe, and injected slowly into the middle of the anterior wall without dissemination. Mice were randomly assigned to treatment groups according to the luminescence signal at day 7 post cell inoculation, and subjected to respective therapies.

For *in vivo* imaging, mice were anesthetized with isoflurane and i.p. injected with 100 μL of a 30 mg/mL D-luciferin solution (Goldbio). After 10 minutes, the mice were imaged using an IVIS Spectrum Optimal Imaging System (Perkin Elmer). Photon flux was measured with Living Image software (Perkin Elmer). Tumor samples were then collected after 4 weeks.

For the liver metastasis model, 1.5×10^6^ ACKP cells were injected via the portal vein, and treatment was initiated 3 days after the cell injection. Mice were euthanized and the liver was excised at 2 weeks post cell inoculation.

For the spontaneous lung metastasis model, the primary subcutaneous ACKP tumors were surgically removed when the tumor volume reached 500-600 mm^3^ to allow tumor metastasis to occur as previously described^41^. The respective treatment was started the second day after the surgery. 6 weeks after tumor resection, the mice were euthanized and lung tissue was extracted for analysis.

For survival analysis of the subcutaneous ACKP tumor model, the mice were deemed dead when the tumors reached 2 cm in diameter or when clinical signs arose, and were euthanized.

For the autochthonous model, Mist1-CreERT; Cdh1^flox/flox^; LSL-RHOA^Y42C/Y42C^; Hdc-GFP mice were administered 6 mg tamoxifen in 200 μL corn oil by gavage for 2 times^85^. Stomach samples were collected at 10 months post tamoxifen induction (see Fig. S2D). Stomach weight was measured after opening the stomach with scissors and thoroughly washing out its contents in PBS.

For antibody-mediated deletion of T and NK cells, the following antibodies: Anti-CD4 antibody (20 mg/kg, Clone YTS 177, Bioxcell, Cat# BE0003-3); Anti-CD8α antibody (20 mg/kg, Clone 53-6.7, Bioxcell, Cat# BE0004-1); Anti-NK1.1 (30 mg/kg, Clone PK136, Bioxcell, Cat# BE0036) were given intraperitoneally every 3 days starting 9 days post cell inoculation.

For antibody-mediated neutralization of IFNγ, CXCL10, and IL12 *in vivo*, the following antibodies: Anti-IFNγ antibody (10 mg/kg, Clone R4-6A2, Bioxcell, Cat# BE0054) ; Anti-CXCL10 (10 mg/kg, Clone 1F11, Bioxcell, Cat# BE0440); Anti-IL12 (10 mg/kg, Clone C17.8, Bioxcell, Cat# BE0051) were given intraperitoneally every 3 days starting 9 days post cell inoculation.

For anti-LY6G mediated total PMN deletion, we adopted a recently improved and verified protocol^13,83^ by administrating the “combo” regimen of 200 μg/mouse anti-Ly6G (Clone 1A8, Bioxcell, Cat# BE0075-1) and 50 μg/mouse anti-rat kappa light chain (Clone MAR 18.5, Bioxcell, Cat# BE0122) every other day starting from 10 days post cell inoculation. For AMD3100 comparison, AMD3100 (Sigma, Cat# A5602) was injected intraperitoneally every 3 days at 10mg/kg.

Sample sizes were determined based on pilot experiments, and power analysis using the G*Power software (Version. 3.1.9.7).

#### Cell and organoid culture

ACKP cells were cultured as previously described in DMEM supplemented with 10% FBS, 100 IU/ml penicillin/streptomycin. PC organoids were cultured using the conditioned medium from L-WRN cells mixed with fresh medium (1:1 ratio)^87^. The medium was replaced every other day. All cells used in this study were routinely screened for mycoplasma contamination using the MycoAlert Mycoplasma Detection Kit (Lonza, Cat# LT07-118).

#### Histological analysis

Tumor histology was analyzed independently by an experienced pathologist in a blinded manner. Quantification of histopathological images was performed using QuPath.v0.5.1.

#### Imaging flow cytometry

After dissociating tumor samples into single cell suspensions and blockade of nonspecific staining using TruStain FcX Plus (BioLegend, Cat# 156603), the following antibodies were used for surface marker staining of samples at 1:200 dilution for 25 minutes on ice: BV510 anti-mouse CD11b, Clone M1/70, Biolegend Cat# 101245; BV605 anti-mouse LY6G, Clone 1A8, Biolegend Cat# 127639; BV421 anti-mouse CD45, Clone 30-F11, Biolegend Cat# 103133; PE-Cy7 anti-mouse LY6C, Clone HK1.4, Biolegend Cat# 128017. 1×10^6^ cells stained with surface markers and DRAQ5 (Biolegend, Cat# 424101, 1:1000) were then re-suspended in 50 μl PBS (Biolegend) for sample acquisition. Imaging flow cytometry and nuclear lobe counting were performed on an ImageStream X MkII imaging flow cytometer (Cytek Biosciences) as previously described^88,89^. Briefly speaking, cells were imaged using the 60 × objective.

Classifiers were set on brightfield to eliminate debris and on fluorescence channels to eliminate saturated images. Focused images of single cells were analyzed using the lobe count feature on the morphology mask of the DRAQ5 signal to compare nuclear segmentation of Hdc-GFP^+^ and Hdc-GFP^-^ cells within the gate of live CD45^+^CD11b^+^LY6G^+^LY6C^-/low^ population. Data was analyzed using IDEAS 5.0 software (Cytek Biosciences). LY6C channel is not shown in figures.

#### T cell proliferation in coculture

Splenocytes from C57BL/6J mice were labeled with Cell Trace Violet 5 μM (Thermo Scientific, Cat# C34557) for 20 minutes at 37°C, and then seeded to flat bottom 96-well pre-coated with anti-CD3 (Clone 145–2C11, BD Biosciences, 1ug/mL)/anti-CD28 (Clone 37.51, Fisher Scientific, Cat# 553295, 5ug/mL into solution) at a density of 2×10^5^/well. PMN-MDSCs were isolated from ACKP tumor cell suspensions using anti-Ly6G Ultrapure Microbeads (Miltenyi Biotec, Cat# 130-120-337) on a MACS separator (Miltenyi Biotec) according to the manufacturer’s protocol, and added to the labeled splenocytes at serial ratios. The mixture was then cocultured in complete T cell medium (RPMI, 10% FBS, 2mM l-glutamine, 100 units/mL penicillin and streptomycin, 1% MEM nonessential amino acids, 1 mmol/L sodium pyruvate, 0.02 mmol/L 2-mercaptoethanol) for 3 days before flow cytometry analysis of T cells. For some experiments, the following agents were added to the coculture: nor-NOHA (Nω-hydroxy-nor-Arginine, arginase inhibitor, Cayman Chemical, Cat# 10006861) at 200μM; NAC (N-acetyl-L-Cysteine, ROS inhibitor, Cayman Chemical, Cat#20261) at 1mM; 1400W (iNOS inhibitor, Cayman Chemical, Cat#81520) at 1μM.

#### Flow cytometry of mouse tissues

Spleen was mashed through a 70 μm cell strainer with the plunger end of a 5 ml syringe into single cell suspension. Blood was obtained from the descending aorta and placed in an EDTA tube on ice. Tumor tissues were minced carefully using scissors 1-2 mm diameter pieces in cell staining buffer (Biolegend, Cat# 420201), and placed in digestion media (10ml HBSS, HEPES 100ul, Collagenase P 10mg, BSA 100mg, and DNase I 10mg per tumor) inside a 50ml tube and incubated on a rotator at 37°C for 25 minutes. The digestion was then stopped by adding 10 ml HBSS including 10% FBS. The cell suspension was passed through a 40μm cell strainer, and removed red blood cells by resuspending in 5 ml of ACK red blood lysis buffer (Gibco) for 5 minutes. Zombie NIR staining was performed using the Zombie NIR Fixable Viability kit (Biolegend, Cat# 423105) according to its instructions for 20 minutes at room temperature. After Zombie NIR staining, cells were incubated with TruStain FcX Plus (BioLegend, Cat# 156603) for 10 minutes on ice, followed by staining with the flow cytometry antibodies all at 1:200 dilution for 25 minutes on ice. Cells are then washed thoroughly and resuspended for flow cytometry analysis.

For myeloid staining the following antibodies were used: PE/Cy7 anti-mouse Ly-6G (Biolegend, Cat# 127618), BV711 anti-mouse Ly-6C (Biolegend, Cat# 128037), BUV496 anti-mouse F4/80 (BD Biosciences, Cat# 750644), BUV615 anti-mouse CD184 (BD Biosciences, Cat# 752552), BV421 anti-mouse CD45 (Biolegend, Cat# 103133), BV650 anti-mouse CD11b (Biolegend, Cat# 101239), PE anti-mouse CD11C (Biolegend, Cat#117307), BV786 anti-mouse I-A/I-E (BD Biosciences, Cat#742894), PE/Dazzle 594 anti-mouse CCR7 (Biolegend, Cat#120121). In some experiments, BUV737 anti-mouse CD54 (BD Biosciences, Cat#741717) and Alexa Fluor 647 anti-mouse CD101 (BD Biosciences, Cat#564473) are further included.

For staining of phosphorylated protein, the sorted cells were stimulated for 20 minutes and fixed with BD Phosflow™ Lyse/Fix Buffer (BD Biosciences, Cat# 558049) for 10 minutes at 37°C water bath at a sample-to-buffer ratio of 1:20. After washing with HBSS, the cells were permeabilized with BD Phosflow™ Perm Buffer III (BD Biosciences, Cat# 558050) for 30 minutes on ice and stained with BV421 anti-mouse p-Akt (1:100, Clone M89-61, BD Biosciences, Cat#562599).

For lymphoid staining, the following antibodies were used: BV421 anti-mouse CD45 (Biolegend, Cat# 103133), BV650 anti-mouse CD3e (BD Biosciences, Cat# 564378), PE/Dazzle594 anti-mouse NK1.1 (Biolegend, Cat# 108747), BV510 anti-mouse CD4 (Biolegend, Cat# 100553), PE/Cy7 anti-mouse CD8a (Biolegend, Cat# 100721), BUV615 anti-mouse Tim-3 (BD Biosciences, Cat# 753150), Spark/NIR685 anti-mouse CD69 (Biolegend, Cat# 104557), BV570 anti-mouse B220 (Biolegend, Cat# 103237). After cell surface marker staining, samples were then fixed with BD Cytofix/Cytoperm™ kit (BD Biosciences, Cat# 554714) for 20 minutes at 4°C according to the manufacturer’s protocol and stained with APC anti-mouse Granzyme B (Biolegend, Cat# 372203) for 30 minutes on ice. For cytokine staining, to stimulate cytokine production, the cell culture medium was added with Leukocyte Activation Cocktail with BD GolgiPlug™ (BD Biosciences, Cat# 550583) and placed in the 37°C cell incubator for 6 hours before flow cytometry staining with Alexa Fluor 700 anti-mouse IFNγ (Biolegend, Cat# 505823) and PE/Cy7 anti-mouse TNFα (Biolegend, Cat# 506323). In this staining, PE/Cy5 CD8a (Biolegend, Cat#100709) was used for compatibility in staining.

For Treg and proliferating T cell staining, cells were first stained with antibodies of surface markers including: Alexa Fluor 700 anti-mouse CD45 (Biolegend, Cat# 103127), BV650 anti-mouse CD3e (BD Biosciences, Cat# 564378), BV510 anti-mouse CD4 (Biolegend, Cat# 100553), PE/Cy7 anti-mouse CD8a (Biolegend, Cat# 100721), PE/Cy5.5 anti-mouse CD25 (Thermo Fisher Scientific, Cat# 35-0251-80). The cells were then fixed with BD Pharmingen™ Transcription Factor Buffer Set (BD Biosciences, Cat# 562574) for 40 minutes at 4°C according to the manufacturer’s protocol and stained with antibodies for 40 minutes on ice including PE anti-mouse FOXP3 (Thermo Fisher Scientific, Cat# 12-5773-82), and PE/Dazzle594 anti-mouse Ki67 (Biolegend, Cat# 151219).

For bone marrow HSPC staining, BM cells were obtained by crushing the femur, tibiae, and pelvic bones in a buffer comprised of Hank’s buffered saline solution (HBSS) containing 3% heat-inactivated fetal bovine serum. After lysis of red blood cells as above, unfractionated BM cells were blocked using purified rat IgG (Sigma-Aldrich, Cat# I8015) diluted 1:100 in staining buffer for 10 minutes on ice. Cells were then incubated with the following lineage antibodies conjugated to PE/Cy5: B220 (1:800, Biolegend, 103209), CD19 (1:800; Biolegend, Cat# 115509), Gr-1 (1:800; Biolegend, Cat# 108409), CD11b (1:800; Biolegend, Cat# 101209), TER119 (1:400; Biolegend, Cat# 116209), CD5 (1:800; Biolegend, Cat# 100609), CD4 (1:800; Biolegend, Cat# 100409), CD8a (1:800; Biolegend, Cat# 100709), together with the following antibodies of stem cell and progenitor markers: APC/Cy7 anti-mouse c-Kit (1:800; Biolegend, Cat# 105825), BV421 anti-mouse Sca-1 (1:400; Biolegend, Cat# 108127), Alexa Fluor 700 anti-mouse CD48 (1:400; Biolegend, Cat# 103425), BV650 anti-mouse CD150 (1:200; Biolegend, Cat# 115931), PE anti-mouse CD135 (1:100; Biolegend, Cat# 135305), PE/Cy7 anti-mouse CD16 (1:800; Biolegend, Cat# 158015), BV605 anti-mouse CD34 (1:50, BD Biosciences, Cat# 750918), BV711 anti-mouse CD115 (1:400, Biolegend, Cat# 135515), and APC anti-mouse Ly6C (1:800, Biolegend, Cat# 128015). Right before analysis, stained cells were resuspended in 1:1000 diluted Propidium Iodide (Biolegend, Cat#421301) solution for dead cell exclusion.

All data recording was performed on a Novocyte Quanteon or Novocyte Penteon machine. All cell sorting using FACS was performed on a Sony MA900 machine. Data collection and analysis were performed using NovoExpress v1.6.2.

#### RT-qPCR (Real-time quantitative PCR)

Total RNA was extracted from the tissue and cells with TRIzol reagent (Invitrogen) and was reverse transcribed into cDNA with qScript cDNA SuperMix (QuantaBio). Expression levels of indicated genes were quantified by qPCR assays using a 7500 Real-time PCR system (Applied Biosystems) with a SYBR Green FastMix Reaction Mix (QuantaBio). Predesigned sequences from Integrated DNA Technologies (IDT) were used in this experiment as shown in the Key resources table.

#### scRNA-seq

For scRNA-seq of tumor CD45^+^ cells, CD45^+^ cells were isolated from ACKP subcutaneous tumors grown in Hdc-GFP mice after 3 cycles of treatments (pooled from 3 mice in each group). To increase the visibility of tumor PMNs in scRNA-seq, given their low RNA content, we employed a strategy of mixing live CD45^+^CD11b^+^LY6G^+^LY6C^-/low^ tumor PMNs with total CD45^+^ cells isolated from the same tumors at a 1:2 ratio during sample preparation for scRNA-seq analysis. For scRNA-seq of splenic PMNs, live CD45^+^CD11b^+^LY6G^+^LY6C^-/low^ population is sorted by FACS from ACKP subcutaneous tumor-bearing Hdc-GFP mice after 3 cycles of treatments (pooled from 3 mice in each group).

Library preparation and sequencing were performed by the JP Sulzberger Columbia Genome Center. Immediately after sorting, cells were counted using an automated cell counter (ThermoFisher Countess II FL) to check viability and processed following the manufacturer’s recommendations for Chromium Single Cell 3’ Library and Gel Bead Kit v.2 (10x Genomics). 10,000 total cells were loaded for each group. Samples were sequenced in an Illumina NovaSeq6000, and CellRanger (v7.1.0) pipeline was used for the genome mapping (mm10) with the default settings and the filtered matrix was used for the data input. After reading the gene expression matrix into R, we filtered out the cells with low quality (nFeature_RNA < 500 and percent.mt > 5). Potential doublets were removed by “DoubletFinder”. Therefore, we recovered 29104 and 22844 single cells from the tumor and spleen, respectively, for downstream analyses. PCA was run by computing 30 principal components and the first 20 principal components were utilized for the computing of Uniform Manifold Approximation and Projection (UMAP) embedding of the cells. Cell clusters were explored using a range of different resolution settings of 0.1/0.3/0.5/0.7. Cell cluster marker genes were conducted by “FindAllMarkers” with these settings: min.pct = 0.2, logfc.threshold = 0.25. By manually checking these cluster markers, we removed two small populations of likely stromal cells and epithelial cell contamination and annotated the rest of the cell clusters with 7 major cell types (PMN, Macro/Mono, T/NK, DC, Cycling T/NK, Mast, B cell). Top markers for each cell type were visualized in the dot plots in Fig. S5. For PMN, we subset cells with PMN labels from all samples and then applied the resolution setting of 0.3 for PMN clustering. Signature scores were calculated using “AddModuleScore” from Seurat with a setting of ctrl = 100 and then scaled. The average gene expression of each group was calculated using “AverageExpression”. Differentially expressed genes (DEGs) for each group were also generated by “FindAllMarkers” and these genes with average logFC of larger than 0 were considered as DEGs. Venn diagram was used to highlight these DEGs were shared and unique to the groups.

For RNA velocity, we first generated loom files from fastq files for each sample using velocyte package. After merging all loom files, we then only select the cells with PMN labels from Seurat objects for RNA velocity calculation. RNA velocity was estimated by Scvelo using the default settings and was visualized in the UMAP embedding.

For human scRNA-seq analysis, the raw sequencing data were aligned and quantified against the human reference genome (hg19) using CellRanger Single Cell Software Suite (version 3.1). The Seurat R package (version 5.1.0) was used to import and process the gene expression matrices.

#### HiBiT assay

The human K562 cell was selected for HiBiT assay as it lacks endogenous CXCR4 expression^53^. To introduce HiBiT-tagged CXCR4 into K562 cells, the following sequence was subcloned into a PiggyBAC transposon vector plasmid: EF1a promoter-HiBiT-3 x GGGS linker-hCXCR4 transcript variant 2 (NM_003467.3)-3 x GGGS linker-EGFP-CMV promoter-Puromycin resistant gene. Gene synthesis was performed by VectorBuilder (San Diego, CA, USA). This CXCR4 vector was co-transfected with the PiggyBac transposase expression vector (Cat# VB900088-2874gzt, VectorBuilder) using TurboFectin Transfection Reagent (Origene). TurboFectin transfection was performed according to the manufacturer’s protocol. From 3 days after transfection, the cell medium was added with 2 ug/mL puromycin to select for CXCR4-expressing cells. Medium with puromycin was replaced every two days to maintain selection. Right before HiBiT assay, CXCR4-expressing K562 was stimulated with SDF-1 or modified TFF2 at serial concentrations for 30 minutes. HiBiT assay was immediately performed as instructed using a HiBiT Extracellular Detection System (Promega, Cat# N2420) that enables accurate quantification of ligand-induced GRCR (G protein-coupled receptors) internalization^90^. Untransfected K562 cell was used for background reduction. TFF2-HSA in this assay was obtained from Tonix Pharmaceuticals.

#### Chemotaxis Assay

Chemotaxis was performed using 6.5 mm Transwell® with 5 µm Pore Polycarbonate Membrane Insert, Sterile (Corning, Cat# 3421). Splenocytes were isolated for CD11b^+^ myeloid cells. After 2 hours of serum starvation (0.1% BSA), 2×10^5^ cells from the same cell suspension were loaded onto the top chamber in 100μl RPMI medium with 0.1% BSA. 600μl Medium supplemented with attractants, BSA, or medium alone was simultaneously added to the bottom chamber and the cells were allowed to migrate towards the gradient for 5 hours. Then the top insert was carefully removed, and cells in the bottom chamber were harvested for flow cytometry analysis.

#### Immunofluorescence

Tissues were fixed overnight in 10% formalin at 4°C and embedded into paraffin blocks. The slides were processed by standard histological methods. Following antigen retrieval, the tissues were blocked with 10% BSA for 1 hour at RT and then stained with the primary antibody overnight. The following primary antibodies were used: anti-CD8 (Abcam, Cat# ab217344, dilution at 1:200). The following day the slides were stained with secondary antibodies and examined under microscope.

#### *In vitro* MP cell culture

MPs (Lin^-^C-kit^+^Sca-1^-^) cells were isolated from mouse bone marrow using lineage depletion kit (Miltenyi Biotec, Cat# 130-090-858) and fluorescence-activated cell sorting (FACS), and seeded in flat-bottom 96-well culture plates (0.5×10^5^ cells/well) and cultured in RPMI-1640 medium containing 10% FBS. The indicated cytokines were added at the following concentration: GM-CSF (Peprotech, Cat# 315-03) at 50 ng/mL, SDF-1 (Peprotech, Cat# 250-20A) at 100 ng/mL, TFF2-MSA at 1μM. SCF was added to all conditions at 20ng/mL. Media was refreshed every 2 days until harvesting for flow cytometric analysis at 5 days post cell seeding.

#### Mouse blood test

Peripheral blood was obtained before sacrificing mice in EDTA-anti-coagulated tubes, and CBCs differentials were determined immediately using Heska Element HT5 Auto Hematology Analyzer.

#### Flow cytometry of human blood

Blood was processed within 4 hours after blood sampling and analyzed using a method modified from previously described^91^. 5-10 ml of fresh human blood was collected in an EDTA tube. Blood was then admixed with the same volume of phosphate-buffered saline (PBS), and carefully overlayed onto 1.077 g/mL separation medium (Ficoll-Paque, Cytiva, Cat#17-1440-02). Density centrifugation was then performed at room temperature 400xg, 30 min without acceleration and brake. Afterward, plasma was aspirated and kept frozen in -80 until analyzed for TFF2 ELISA as below. Interface PBMCs were carefully collected using a pipette for flow cytometry analysis. The bottom layer was used for flow cytometry analysis of HDN. PBMCs were lysed with 5 ml of ACK red blood lysis buffer (Gibco) 2 times, and washed with cell staining buffer (Biolegend, Cat# 420201), followed by 575V cell viability stain (BD Biosciences, Cat# 565694) at RT for 15 minutes. Unspecific staining was then blocked with Human Trustain FcX (Biolegend, Cat# 422302) 5 ul in 100 ul cell staining buffer for 10 min at RT. The samples were followed by staining in antibodies all at 1:200 dilution for 25 minutes at 4°C.

For myeloid staining, the following antibodies were used : PE anti-human CD33 (Biolegend, Cat# 303403), FITC anti-human CD11b (Biolegend, Cat# 301329), BV711 anti-human CD14 (BD Biosciences, Cat# 563372), PE/CF594 anti-human HLA-DR (BD Biosciences, Cat# 563372), APC/Cy7 anti-human CD45 (Biolegend, Cat# 561863), APC anti-human Lineage Cocktail (CD3, CD19, CD20, CD56) (Biolegend, Cat# 363601), Alexa Fluor 700 anti-human CD66b (Biolegend, Cat# 305113), BV421 anti-human LOX-1 (Biolegend, Cat# 358609), PerCP/Vio700 anti-human CD16 (Miltenyi Biotec, Cat# 130-113-395), BV510 anti-human CD184 (Biolegend, Cat# 306535), PE/Cy7 anti-human CD15 (BD Biosciences, Cat# 560827), BV786 anti-human CD10 (BD Biosciences, Cat# 744754). For Arginase 1 staining, the above panel was modified by adding PE anti-human Arginase 1 (Biolegend, Cat# 369703), and replacing the CD10 antibody with BV786 anti-human CD33 (BD Biosciences, Cat# 740974). Human total MDSCs were defined as CD45^+^CD11b^+^CD33^+^LIN^-^HLA-DR^-^, human PMN-MDSCs were defined as CD45^+^CD11b^+^CD33^+^LIN^-^HLA-DR^-^CD14^-^CD66b^+^, human M-MDSCs were defined as CD45^+^CD11b^+^CD33^+^LIN^-^HLA-DR^-^CD14^+^CD66b^-^, human eMDSCs were defined as CD45^+^CD11b^+^CD33^+^LIN^-^HLA-DR^-^CD14^-^CD66b^-^.

For lymphoid staining, a small aliquot of blood was stained with the following antibodies: BV510 anti-human CD45 (Biolegend, Cat# 304035), FITC anti-human CD8 (Biolegend, Cat# 344703), APC anti-human CD3 (Biolegend, Cat# 317317), APC/Cy7 anti-human CD4 (Biolegend, Cat# 300517). After cell surface marker staining, samples were then fixed with BD Cytofix/Cytoperm™ kit (BD Biosciences, Cat# 554714) for 20 minutes at 4°C according to the manufacturer’s protocol and stained with PerCP/Cy5.51 anti-human granzyme B (Biolegend, Cat# 372211) for 30 minutes on ice. All data recording was performed on a Novocyte Penteon machine. Data collection and analysis were performed using NovoExpress v1.6.2.

#### Human PMN-MDSC functional assay

After blood processing, PBMCs were depleted of CD66b^+^ cells using anti-human CD66b biotin antibody (Clone REA306, Miltenyi Biotec, Cat# 130-118-983) and anti-biotin microbeads (Miltenyi Biotec, Cat#130-090-485) on a MACS separator (Miltenyi Biotec). Cell trace violet– labeled PBMCs or CD66b-depleted PBMCs were then stimulated with immobilized anti-CD3 (10 μg/mL, Clone OKT-3, Thermo Scientific, Cat# 16-0037-81) and soluble anti-CD28 (1 μg/mL, Clone 28.2, Thermo Scientific, Cat# 16-0289-81) and proliferation was examined by flow cytometry after 72 h, as previously described^92^.

#### ELISA

Plasma levels of TFF2 were measured with a commercially available human TFF2 ELISA kit (Abcam, Cat# ab275553) according to the manufacturer’s instructions.

#### Patient survival analysis

Kaplan-Meier analysis of the microarray data from human GC samples was analyzed and generated by the KM Plotter website (http://kmplot.com/analysis/)^93^. TFF2 was searched as a gene symbol, and the 214476_at Affymetrix ID was chosen. The median cutoff was selected to split patients. The analysis was performed using the following published human GC datasets: GSE14210, GSE15459, GSE22377, GSE29272, GSE51105, and GSE62254.

#### Statistics

Statistical analysis was performed with Graphpad prism 9. Methods for statistical analysis were used as noted in each figure legend. The n in figure legends indicates the numbers of independent biological replicates. P<0.05 was considered to indicate a statistically significant difference.

