## supplemental figures for "A CXCR4 partial agonist improves immunotherapy by targeting polymorphonuclear myeloid-derived suppressor cells and cancer-driven granulopoiesis"

**Supplemental Information**

Figures S1-17 and figure legends, Table S1-S6

**Fig. S1**

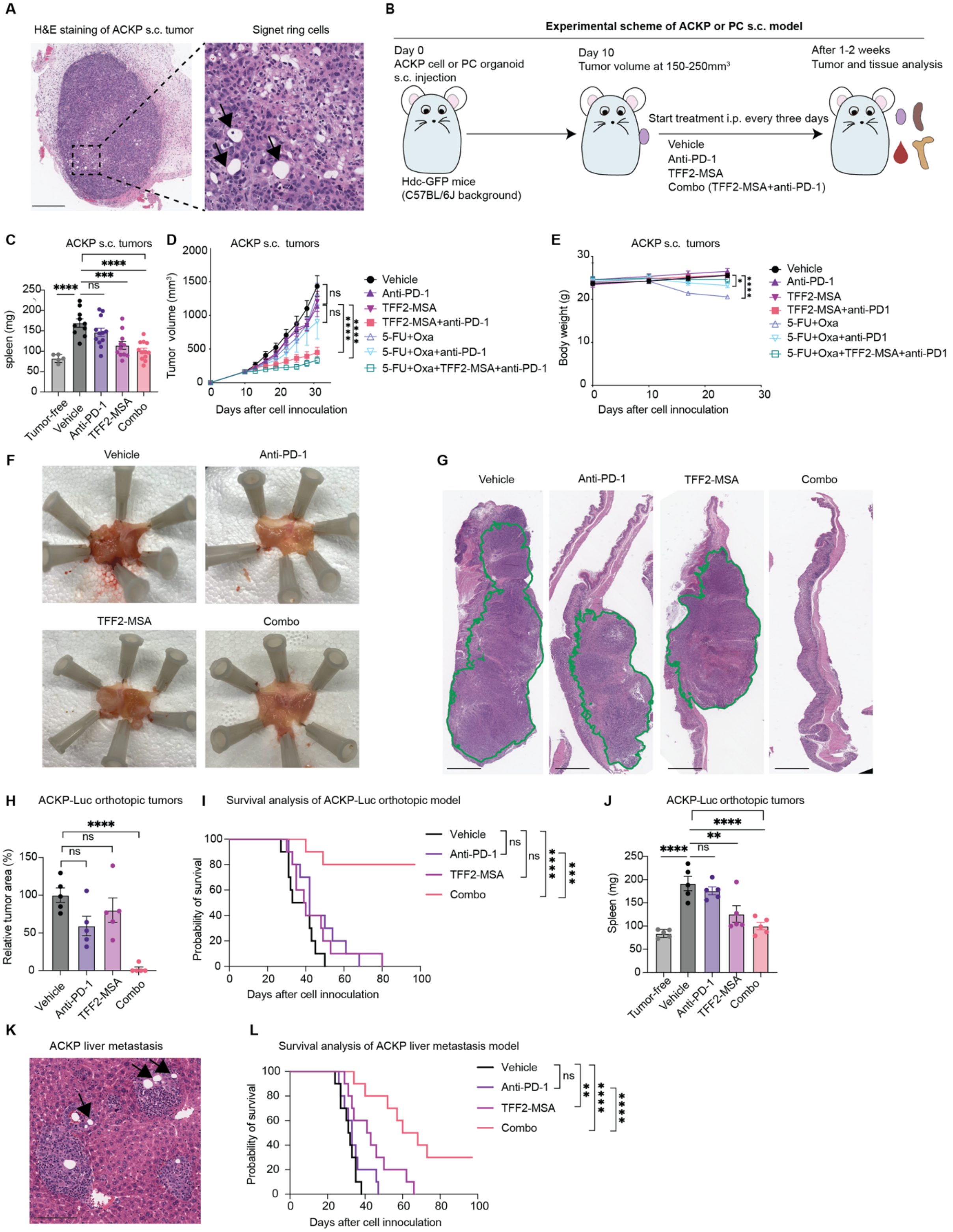
A
H&E staining of ACKP s.c. tumor
Signet ring cells
B
Experimental scheme of ACKP or PC s.c. model
Day 0
ACKP cell or PC organoid
s.c. injection
Hdc-GFP mice
(C57BL/6J background)
Day 10
Tumor volume at 150-250mm³
Start treatment i.p. every three days
Vehicle
Anti-PD-1
TFF2-MSA
Combo (TFF2-MSA+anti-PD-1)
After 1-2 weeks
Tumor and tissue analysis
C
ACKP s.c. tumors
spleen (mg)
Tumor-free
Vehicle
Anti-PD-1
TFF2-MSA
Combo
D
ACKP s.c. tumors
Tumor volume (mm³)
Days after cell innoculation
Vehicle
Anti-PD-1
TFF2-MSA
TFF2-MSA+anti-PD-1
5-FU+Oxa
5-FU+Oxa+anti-PD-1
5-FU+Oxa+TFF2-MSA+anti-PD-1
E
ACKP s.c. tumors
Body weight (g)
Days after cell innoculation
Vehicle
Anti-PD-1
TFF2-MSA
TFF2-MSA+anti-PD1
5-FU+Oxa
5-FU+Oxa+anti-PD1
5-FU+Oxa+TFF2-MSA+anti-PD1
F
Vehicle
Anti-PD-1
TFF2-MSA
Combo
G
Vehicle
Anti-PD-1
TFF2-MSA
Combo
H
ACKP-Luc orthotopic tumors
Relative tumor area (%)
Vehicle
Anti-PD-1
TFF2-MSA
Combo
I
Survival analysis of ACKP-Luc orthotopic model
Probability of survival
Days after cell innoculation
Vehicle
Anti-PD-1
TFF2-MSA
Combo
J
ACKP-Luc orthotopic tumors
Spleen (mg)
Tumor-free
Vehicle
Anti-PD-1
TFF2-MSA
Combo
K
ACKP liver metastasis
L
Survival analysis of ACKP liver metastasis model
Probability of survival
Days after cell innoculation
Vehicle
Anti-PD-1
TFF2-MSA
Combo

**Fig. S1. Related to Figure 1. TFF2-MSA improves anti-PD-1 efficacy in GC mouse models using ACKP cells. A.** Representative H&E staining image of syngeneic tumors established from subcutaneously injected ACKP cells. Arrows indicate signet ring cells. Scale bar: 400µm. **B.** Experimental scheme of syngeneic ACKP cell or PC organoid-derived s.c. tumor model. **C.** Spleen weight of the tumor-free mice and subcutaneous ACKP tumor-bearing mice receiving the indicated therapies (n=5 or 10-12 per group). **D.** Subcutaneous ACKP tumor growth curve in Hdc-GFP mice that received treatment of vehicle, anti-PD-1, TFF2-MSA, chemotherapy, or their combination (n=14 per group). 5-FU, 5-fluorouracil. Oxa, oxaliplatin. **E.** Continuous measurement of body weight of subcutaneous ACKP tumor-bearing mice subjected to the indicated treatments (n=14 per group). 5-FU, 5-fluorouracil. Oxa, oxaliplatin. **F.** Representative macroscopic images of orthotopic stomach tumors from injected luciferase-expressing ACKP cells subjected to the indicated treatments (n=5 per group). **G.** Representative H&E staining of orthotopic stomach tumors from injected luciferase-expressing ACKP cells subjected to the indicated treatments (n=5 per group). The tumor area is outlined. Scale bar: 2mm. **H.** Quantification of orthotopic ACKP tumor area in each group (n=5 per group), as normalized to the average tumor area of the vehicle-treated mice as 100%. **I.** Kaplan-Meier curve of ACKP orthotopic tumor-bearing mouse survival in response to the indicated treatments (n= 10 per group). **J.** Spleen weight of the tumor-free mice and orthotopic ACKP tumor-bearing mice receiving the indicated therapies (n=5 per group). **K.** Representative H&E staining of liver metastasis from portal vein injected ACKP cells. Arrows indicate signet ring cells. Scale bar, 100 µm. **L.** Kaplan-Meier curve of ACKP liver metastasis-bearing mouse survival in response to the indicated treatments (n= 10 per group). stomach tumors. Mean ± SEM are shown, and P values are calculated by one-way ANOVA (**C, H, J**), two-way ANOVA (**D, E**), or log-rank test (**I, L**).

*P < 0.05, **P < 0.01, ***P < 0.001, ****P < 0.0001, ns, not significant.

### Fig. S2

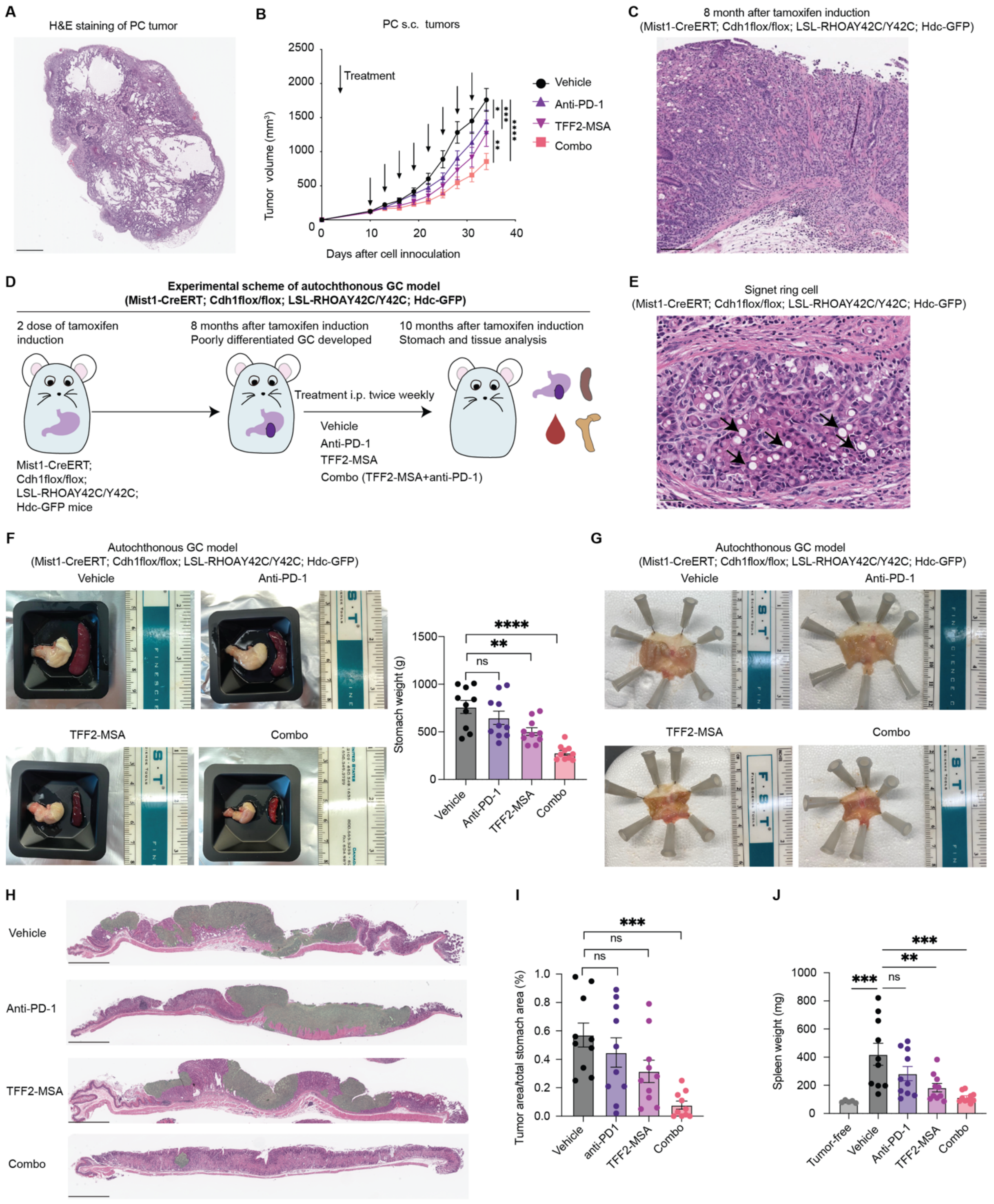

**Fig. S2. Related to Figure 1. TFF2-MSA improves the anti-PD-1 efficacy in other immunocompetent GC mouse models. A.** Representative H&E staining image of syngeneic tumors established from subcutaneously injected PC organoids. Scale bar: 800µm. **B.** Subcutaneous PC tumor growth curve in Hdc-GFP mice that received treatment as indicated (n=10 per group). **C.** Representative H&E staining of Mist1-CreERT; Cdh1$^{flox/flox}$; LSL-RHOA$^{Y42C/Y42C}$; Hdc-GFP mice stomach at 8 months after tamoxifen induction. Scale bar, 200 µm. **D.** Experimental scheme of the autochthonous Mist1-CreERT; Cdh1$^{flox/flox}$; LSL-RHOA$^{Y42C/Y42C}$; Hdc-GFP model. **E.** Representative images of invaded signet ring cells at 10 months after tamoxifen induction in the vehicle-treated Mist1-CreERT; Cdh1$^{flox/flox}$; LSL-RHOA$^{Y42C/Y42C}$; Hdc-GFP mice. Scale bar, 40 µm. **F.** Representative macroscopic pictures of the abnormal stomach (linitis plastica) and the splenomegaly of the Mist1-CreERT; Cdh1$^{flox/flox}$; LSL-RHOA$^{Y42C/Y42C}$; Hdc-GFP mice at 10 months post tamoxifen induction following the indicated treatments. Quantification of stomach weight is shown (n=10 per group)**. G.** Representative macroscopic images showing stomach wall thickening in Mist1-CreERT; Cdh1$^{flox/flox}$; LSL-RHOA$^{Y42C/Y42C}$; Hdc-GFP mice at 10 months post tamoxifen induction following the indicated treatments (n=10 per group). **H.** Representative H&E staining of autochthonous stomach tumors in Mist1-CreERT; Cdh1$^{flox/flox}$; LSL-RHOA$^{Y42C/Y42C}$; Hdc-GFP mice subjected to the indicated treatments (n=10 per group). The tumor area is colored green. Scale bar, 2mm. **I.** Quantification of the ratio of tumor area in the whole stomach area in the H&E staining in F. (n=10 per group). **J.** Spleen weight of the tumor-free mice and autochthonous GC-bearing Mist1-CreERT; Cdh1$^{flox/flox}$; LSL-RHOA$^{Y42C/Y42C}$; Hdc-GFP mice that received the indicated therapies (n=5 or 10 per group). Mean ± SEM are shown, and P values are calculated by one-way ANOVA (**F, I, J**), or two-way ANOVA (**B**). $*P < 0.05$, $**P < 0.01$, $***P < 0.001$, $****P < 0.0001$, ns, not significant.

Fig. S3

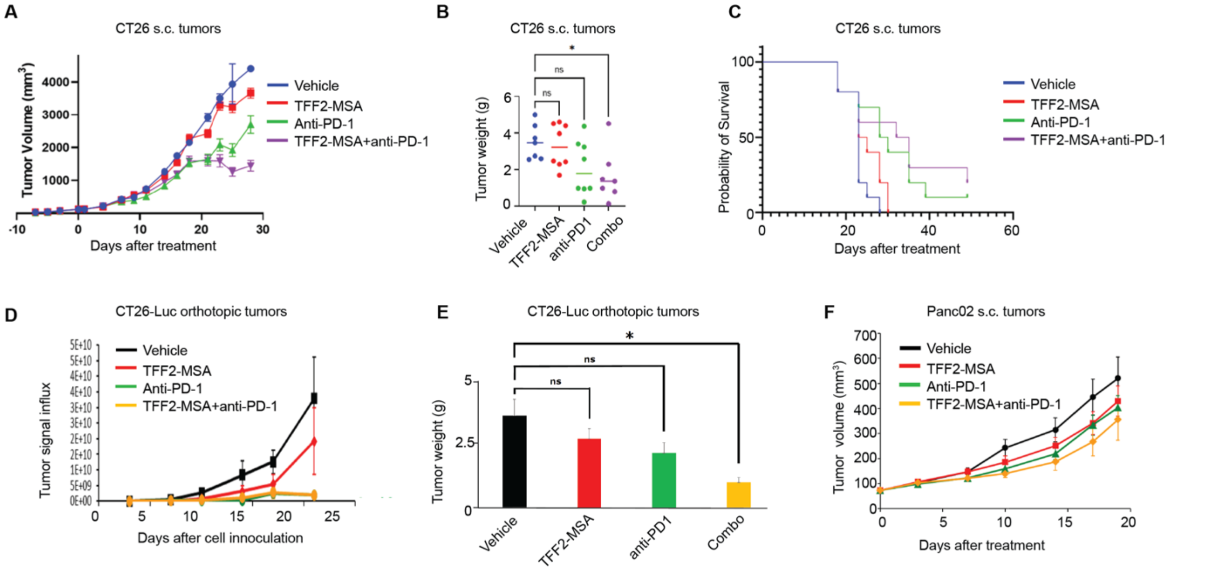


**Fig. S3. Related to Figure 1. TFF2-MSA improves the efficacy of anti-PD-1 in immunocompetent mouse models of colorectal and pancreatic cancer. A**. Subcutaneous CT26 tumor growth curve in C57BL/6J mice that received treatment of vehicle, anti-PD-1, TFF2-MSA, or their combination (n=7-8 per group). **B.** CT26 subcutaneous tumor weight at day 23 post cell inoculation (n=7-8 per group). **C.** Kaplan-Meier curve of CT26 subcutaneous tumor-bearing mouse survival in response to the indicated treatments (n= 7-8 per group). **D.** Quantification of luminescence showing stomach tumors of orthotopically injected luciferase-expressing CT26 cells subjected to the indicated treatments (n=5 per group). **E.** Tumor weight from orthotopically injected luciferase-expressing CT26 cells at day 22 post cell inoculation (n=5 per group). **F.** Subcutaneous Panc02 tumor growth curve in C57BL/6J mice that received treatment of vehicle, anti-PD-1, TFF2-MSA, or their combination (n=7-8 per group). Mean ± SEM are shown, and P values are calculated by one-way ANOVA (**B, E**). *P < 0.05, **P < 0.01, ***P < 0.001, ****P < 0.0001, ns, not significant.

**Fig. S4**

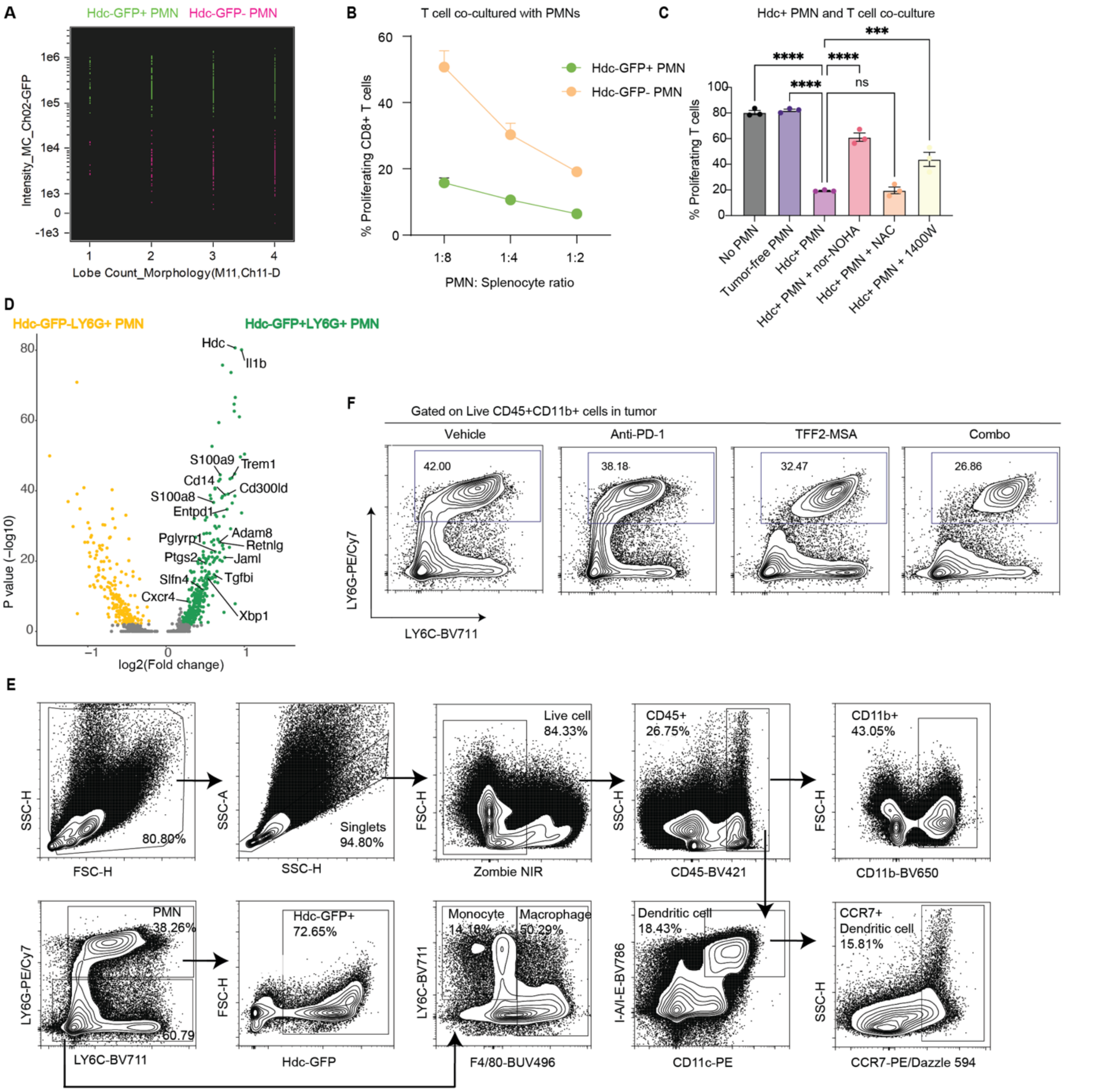


**Fig. S4. Related to Figure 2. Hdc-GFP$^+$ and Hdc-GFP$^-$ subsets of tumor PMNs. A.** Distribution of Hdc-GFP$^+$ (green) and Hdc-GFP$^-$ (pink) PMNs by the number of identified nuclear lobes.**B.** Evaluation of CD8$^+$ T cell proliferation in coculture with Hdc-GFP$^+$ and Hdc-GFP$^-$ tumor PMNs at the indicated PMN to splenocytes ratios. **C.** CD8$^+$ T cell proliferation stimulated with aCD3/CD28 and cocultured with tumor Hdc-GFP$^+$ PMN-MDSCs from the indicated groups at a PMN to splenocytes ratio of 1:8. PMNs from tumor-free mice spleen were used for comparison. Putative inhibitors of PMN-MDSC immunosuppression were added into the coculture: nor-NOHA (Nω-Hydroxy-nor-l-arginine), arginase inhibitor at 200μM. NAC (N-acetyl-L-Cysteine), ROS inhibitor at 1mM. 1400W, iNOS inhibitor at 1μM. **D.** Volcano plot for differentially expressed genes in the Hdc-GFP+ versus Hdc-GFP- PMNs from vehicle-treated ACKP tumors detected by scRNA-seq ($log_2FC \geq 0.2$ and adjusted p value$\leq$ 0.01). The x axis shows the calculated fold change of genes upregulated (green) or downregulated (yellow) in Hdc-GFP+ PMNs versus Hdc-GFP- PMNs. The y axis indicates the calculated adjusted p value. **E.** Gating strategy for myeloid cells in mice. **F.** Representative flow cytometry plots of LY6G+LY6C-/low PMNs in ACKP tumor with the indicated treatments. Mean ± SEM are shown, and P values are calculated by one-way ANOVA (**C**). ***P < 0.001, ****P < 0.0001, ns, not significant.

**Fig. S5**

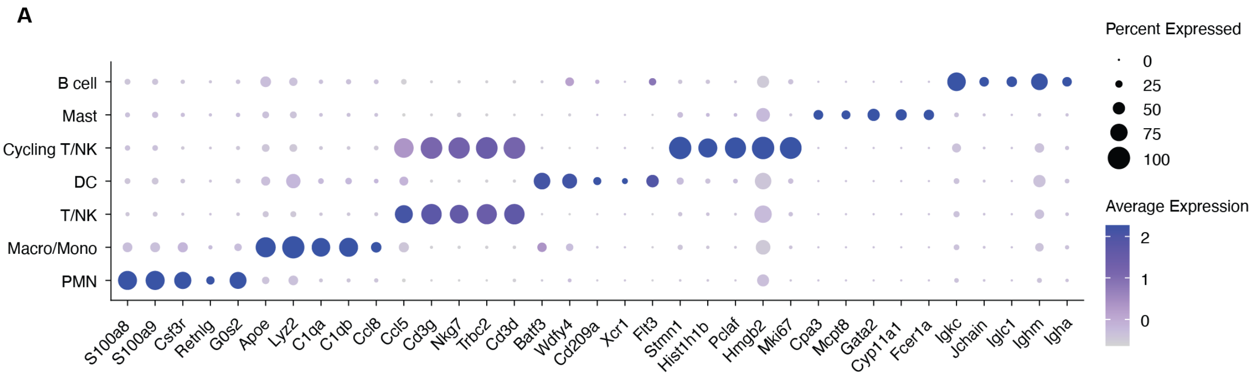


**Fig. S5. Related to Figure 2. ScRNA-seq analysis of CD45+ cells isolated from ACKP tumors. A.** Gene expression of lineage-defining markers from scRNA-seq analysis of CD45+

cell in the TME. The y axis shows cell clusters, the x axis shows characteristic genes in the clusters.

### Fig. S6

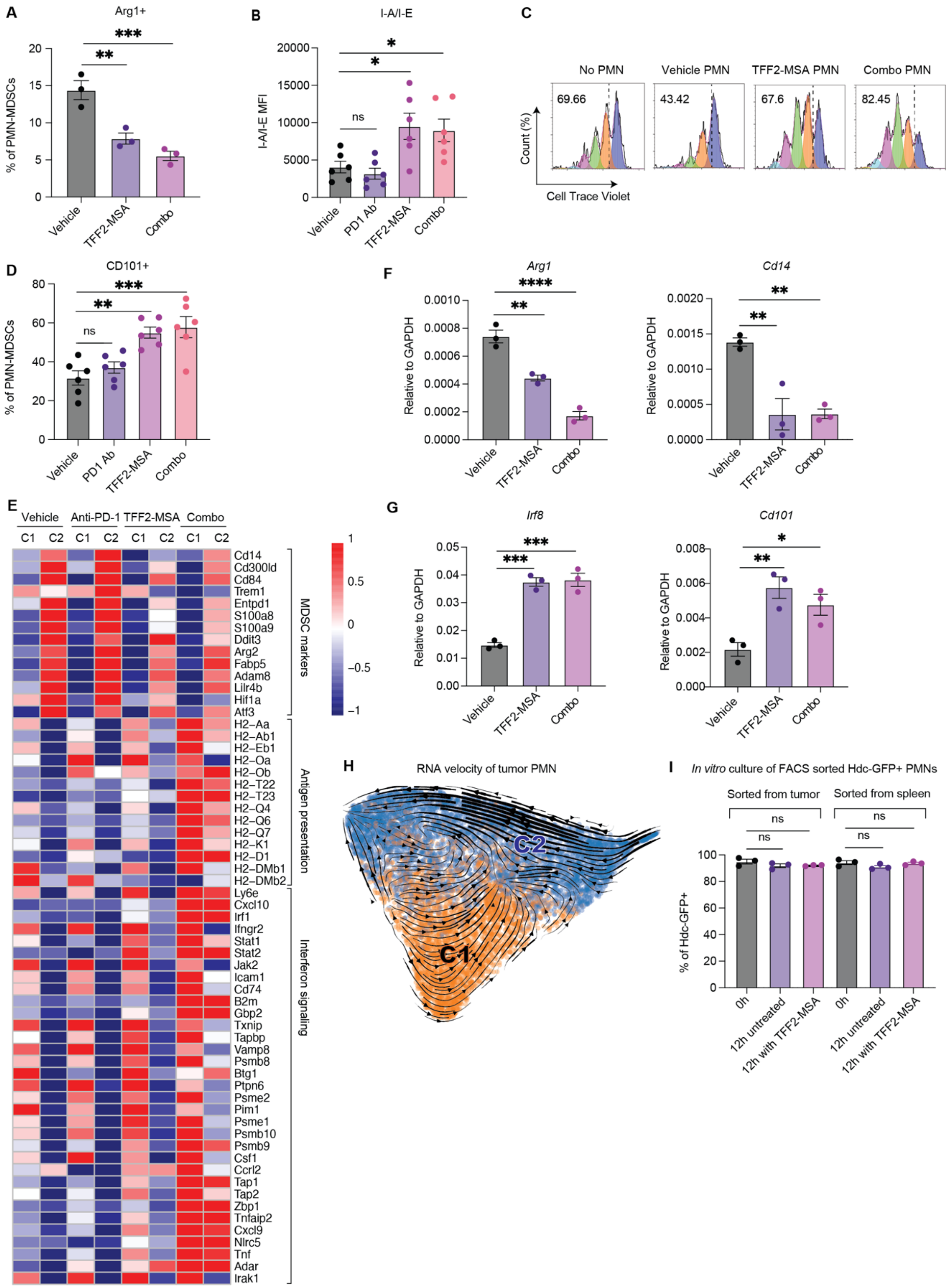
A
Arg1+
% of PMN-MDSCs
Vehicle
TFF2-MSA
Combo
B
I-A/I-E
I-A/I-E MFI
ns
Vehicle
PD1 Ab
TFF2-MSA
Combo
C
No PMN
Vehicle PMN
TFF2-MSA PMN
Combo PMN
69.66
43.42
67.6
82.45
Count (%)
Cell Trace Violet
D
CD101+
% of PMN-MDSCs
F
Arg1
Cd14
Relative to GAPDH
E
Vehicle
Anti-PD-1
TFF2-MSA
Combo
C1
C2
MDSC markers
Antigen presentation
Interferon signaling
G
Irf8
Cd101
H
RNA velocity of tumor PMN
C1
C2
I
In vitro culture of FACS sorted Hdc-GFP+ PMNs
Sorted from tumor
Sorted from spleen
% of Hdc-GFP+
0h
12h untreated
12h with TFF2-MSA

**Fig.S6. Related to Figure 2. TFF-MSA inhibited immunosuppression of tumor PMN-MDSCs. A.** Frequency of Arginase $1^+$ cells within tumor Hdc-$GFP^+$ PMN-MDSCs from the indicated treatment groups (n=3 per group). **B.** Expression of I-A/I-E on tumor PMN-MDSCs from the indicated treatment groups (n=3 per group), expressed as mean fluorescence intensity (MFI). **C.** Representative flow cytometry plots showing T cell proliferation in coculture with PMNs isolated from vehicle, TFF2-MSA, TFF2-MSA plus anti-PD-1 treated ACKP tumors. Coculture is performed at a PMN to splenocyte ratio of 1:8, and T cell without PMN coculture was used as control. **D.** Frequency of $CD101^+$ cells within tumor PMN-MDSCs from the indicated treatment groups (n=3 per group). **E.** Heatmap showing manually selected gene expressions in C1 and C2 PMN subsets from the indicated treatment conditions. **F, G.** RT-qPCR showing expressions of key genes in positive (*Arg1*, *Cd14*) or negative indicators (*Irf8*, *Cd101*) of PMN-MDSCs in tumor Hdc-$GFP^+$ PMN-MDSCs isolated from the indicated treatment groups (n=3 per group). **H.** RNA velocity trajectory analysis of C1 and C2 subsets in PMN. **I.** Changes of Hdc-$GFP^+$ percentage of sorted Hdc-$GFP^+$ PMNs from tumor and matched spleen following *in vitro* culture with or without TFF2-MSA for 12 hours. Mean ± SEM are shown, and P values are calculated by one-way ANOVA (**A, B, D, F, G, I**). *P < 0.05, **P < 0.01, ***P < 0.001, ****P < 0.0001, ns, not significant.

### Fig. S7

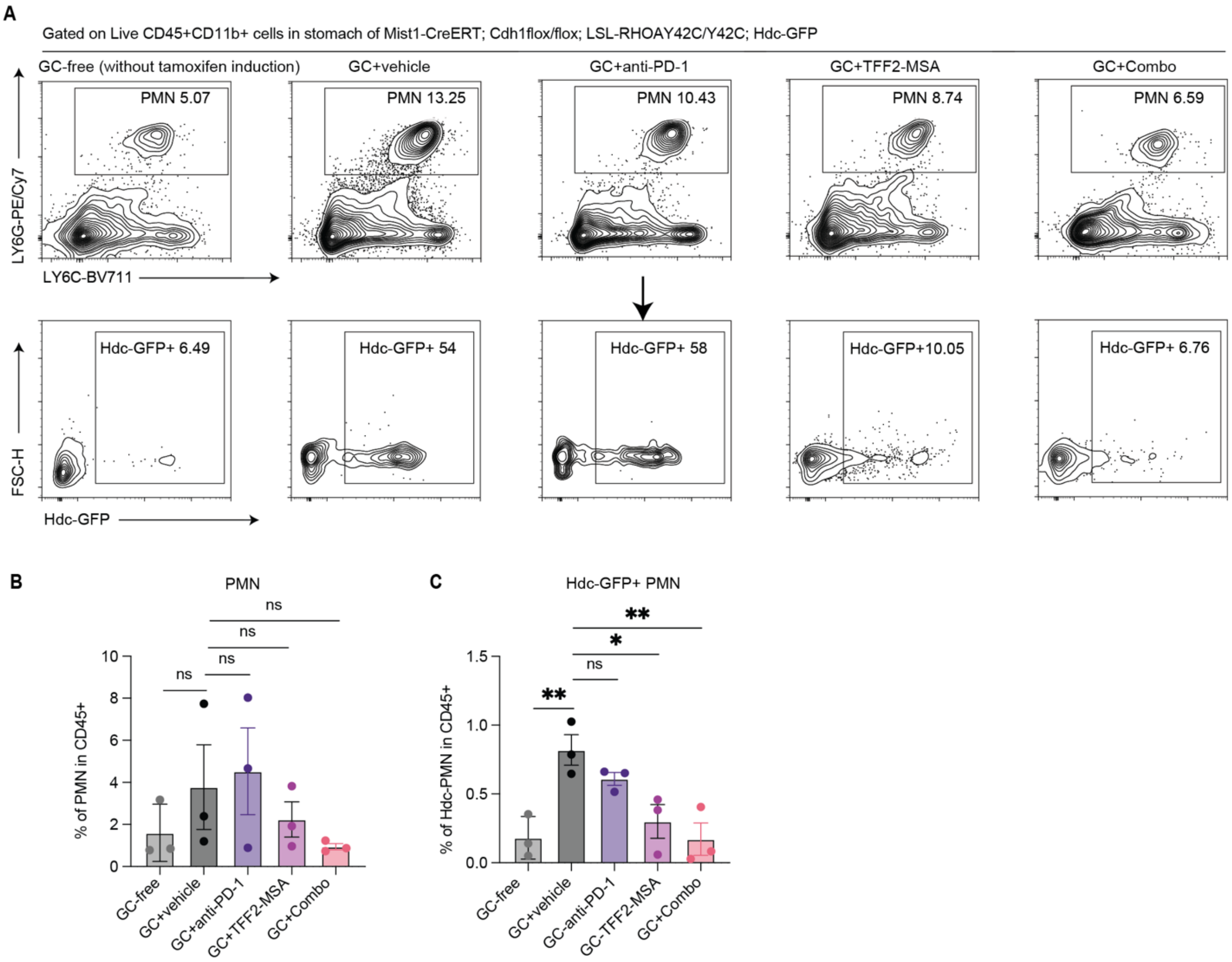


**Fig. S7. Related to Figure 2. TFF-MSA selectively reduced Hdc-GFP+ PMN-MDSCs in the autochthonous GCs of Mist1-CreERT; Cdh1$^{flox/flox}$; LSL-RHOA$^{Y42C/Y42C}$; Hdc-GFP model. A, B, C.** Representative flow cytometry plots of LY6G$^{+}$LY6C$^{-/low}$ PMNs and Hdc-GFP$^{+}$ PMN-MDSCs in the stomach of Mist1-CreERT; Cdh1$^{flox/flox}$; LSL-RHOA$^{Y42C/Y42C}$; Hdc-GFP mice bearing autochthonous GCs is shown in A. Mice of the same genotype and age without tamoxifen induction were used as tumor-free controls (n=3 per group). Quantifications of LY6G$^{+}$LY6C$^{-/low}$ PMNs and Hdc-GFP$^{+}$ PMN-MDSCs in the stomach were shown respectively in B and C. Mean ± SEM are shown, and P values are calculated by one-way ANOVA (**B, C**). *P < 0.05, **P < 0.01, ns, not significant.

**Fig. S8**

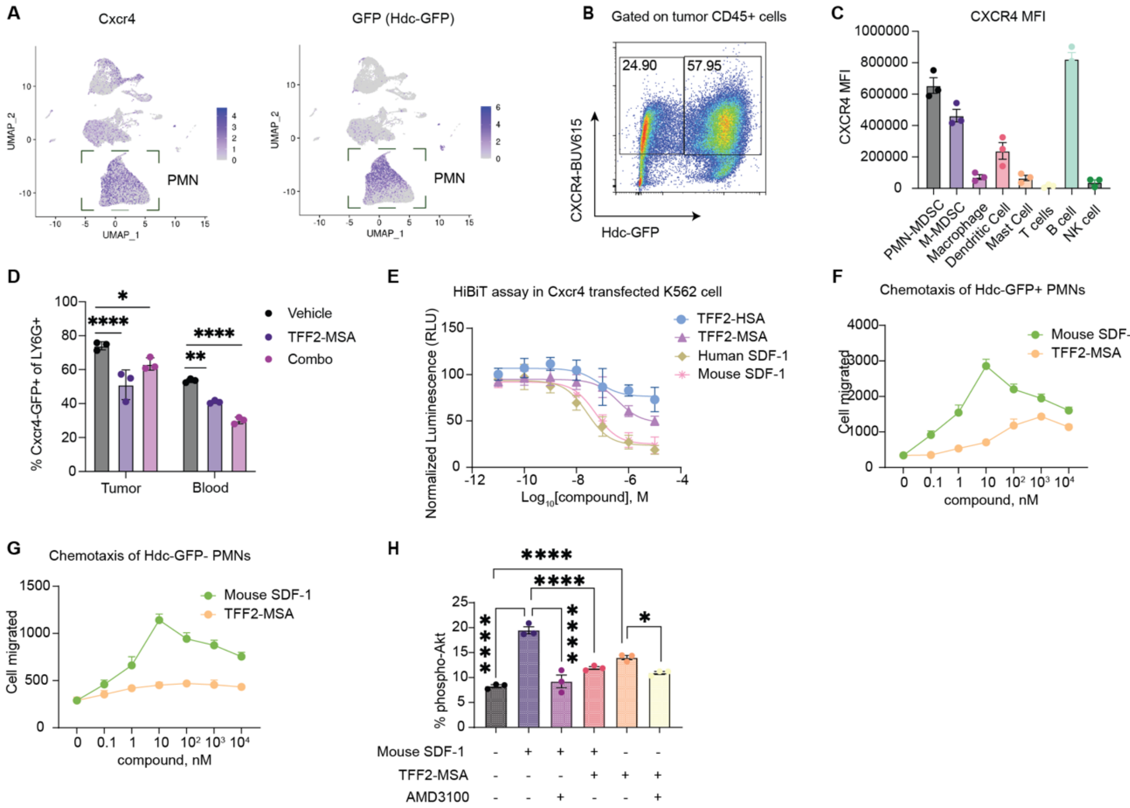


**Fig. S8. Related to Figure 2. TFF-MSA targets CXCR4 on PMN-MDSCs as a partial agonist. A.** UMAP depicting the distribution of CXCR4 and Hdc-GFP expression in total CD45$^+$ cells from ACKP tumors. The dashed box highlights their expression in PMNs defined **Fig. 2F**. **B.** Representative flow cytometry plots of CXCR4 and Hdc-GFP expression gated on CD45$^+$ cells from subcutaneous ACKP tumors. **C.** Representative flow cytometry histograms showing Cxcr4-GFP expression in immune cell populations from the subcutaneous ACKP tumor (n=3 per group). **D.** Quantification of Cxcr4-GFP$^+$ frequency in LY6G$^+$ PMNs from tumor and blood of ACKP tumor-bearing Cxcr4-GFP mice receiving the indicated therapies. Tumor-free mice were used as a control (n=3 per group). **E.** HiBiT assay in human K562 cells overexpressing a HiBiT-tagged CXCR4 construct showing a concentration-dependent internalization of CXCR4 in response to 30 minutes of the indicated agonists or partial agonists at indicated concentrations. The background-subtracted luminescence was normalized to untreated cells. TFF2-HSA is the modified human version of TFF2-MSA. HSA, human serum albumin. **F-G.** Chemotaxis assay showing increasing concentrations of SDF-1 or TFF2-MSA induced migration of splenic LY6G$^+$ PMNs in transwell plates (5 µm pore size). Migrated Hdc-GFP$^+$ (**F**) and Hdc-GFP$^-$ (**G**) cell number after 5 hours of treatment were evaluated by flow cytometry respectively. Data are pooled from 3 independent biological replicates. **H.** Frequency of phospho-$Akt^+$ in splenic $LY6G^+$ PMNs with the indicated treatment of agonist SDF-1 (12.5nM), partial agonist TFF2-MSA (1μM), CXCR4 antagonist AMD3100 (5μM), or their combination acquired by flow cytometry (n=3 per group). Mean ± SEM are shown, and P values are calculated by one-way ANOVA (**D, H**). *P < 0.05, **P < 0.01, ***P < 0.001, ****P < 0.0001, ns, not significant.

### Fig. S9

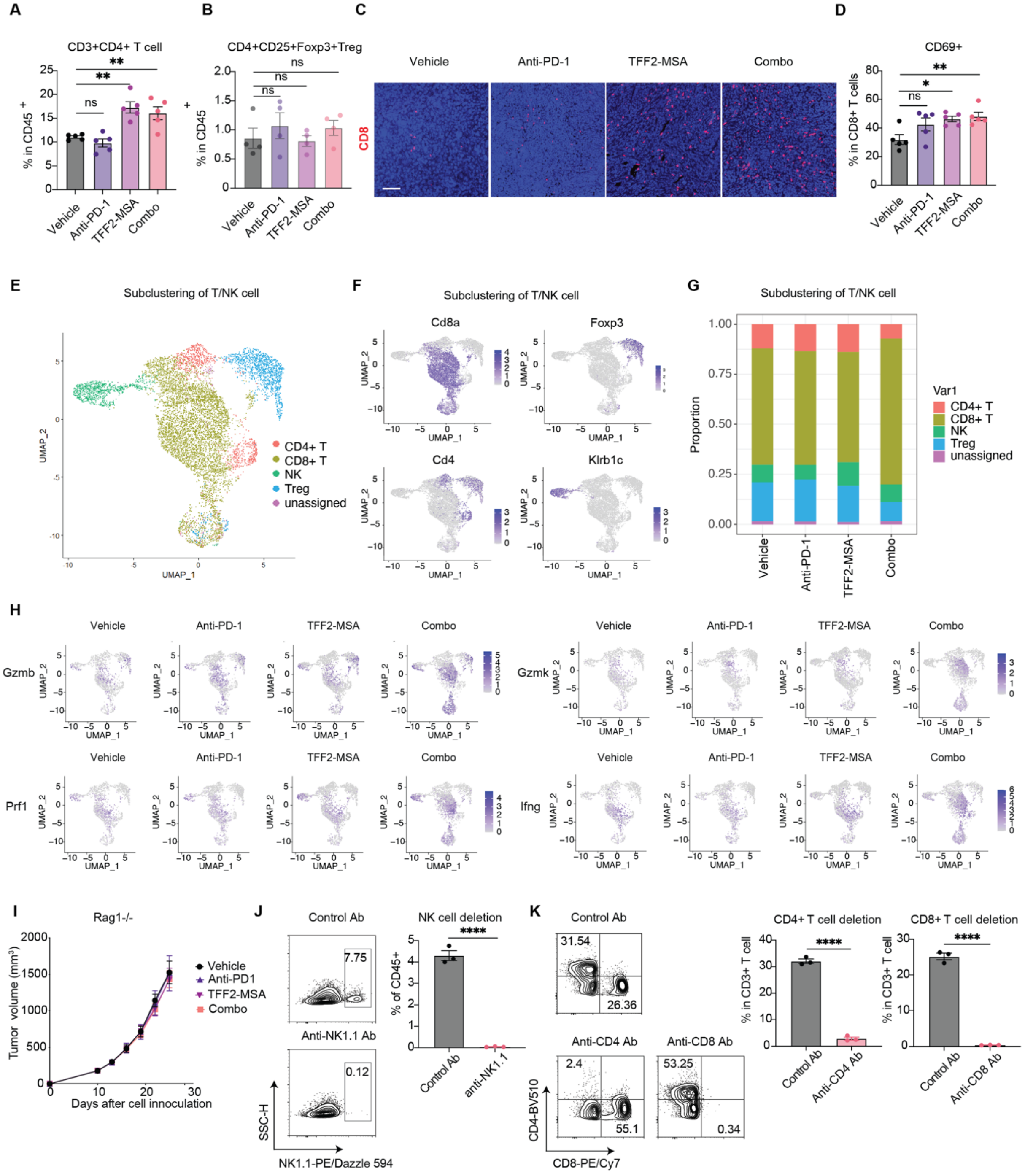
A
CD3+CD4+ T cell
% in CD45+
ns
**
Vehicle
Anti-PD-1
TFF2-MSA
Combo
B
CD4+CD25+Foxp3+Treg
% in CD45+
ns
C
Vehicle
Anti-PD-1
TFF2-MSA
Combo
CD8
D
CD69+
% in CD8+ T cells
ns
*
**
E
Subclustering of T/NK cell
UMAP_1
UMAP_2
CD4+ T
CD8+ T
NK
Treg
unassigned
F
Subclustering of T/NK cell
Cd8a
Foxp3
Cd4
Klrb1c
G
Subclustering of T/NK cell
Proportion
Var1
H
Gzmb
Gzmk
Prf1
Ifng
I
Rag1-/-
Tumor volume (mm3)
Days after cell innoculation
Vehicle
Anti-PD1
TFF2-MSA
Combo
J
Control Ab
7.75
Anti-NK1.1 Ab
0.12
SSC-H
NK1.1-PE/Dazzle 594
NK cell deletion
% of CD45+
****
Control Ab
anti-NK1.1
K
Control Ab
31.54
26.36
Anti-CD4 Ab
2.4
55.1
Anti-CD8 Ab
53.25
0.34
CD4-BV510
CD8-PE/Cy7
CD4+ T cell deletion
% in CD3+ T cell
Anti-CD4 Ab
CD8+ T cell deletion
Anti-CD8 Ab

**Fig. S9. Related to Figure 3. Robust anti-tumor $CD8^+$ T cell response induced by the combination of TFF2-MSA with anti-PD-1 immunotherapy. A.** Analysis of $CD4^+$ T cells in ACKP tumors with the indicated treatments (n=5 per group). **B.** Analysis of $CD4^+CD25^+FOXP3^+$ Tregs in ACKP tumors with the indicated treatments (n=4 per group). **C.** Representative enlarged images showing CD8 immunostaining at the center of ACKP tumors with the indicated therapies. Scale bars, 10 μm. **D.** CD69 expression in the tumor $CD8^+$ T cells with the indicated therapies (n=5 per group). **E.** UMAP depicting the $CD4^+$, $CD8^+$ T cells, Tregs, NK cell, and undefined clusters within the T/NK cells defined in **Fig. 2F** from the ACKP tumors. **F.** UMAP showing the expression of characteristic markers in T/NK cells used to define $CD4^+$ and $CD8^+$ T cells in **G.** Stacked bar graphs depicting the frequency of each cluster defined in **E** within T/NK population from ACKP tumors of the indicated treatment groups. **H.** UMAP showing the expression of granzyme B (*Gzmb*), perforin 1 (*Prf1*), granzyme K (*Gzmk*), and interferon γ (*Ifng*) in T/NK cells from the indicated treatment groups. **I.** Growth curve of subcutaneously implanted ACKP tumor in $Rag1^{-/-}$ mice subjected to the indicated treatments starting 10 days post cell inoculation (n= 10 per group). **J.** Representative flow cytometry plots (left) and quantification (right) of $NK1.1^+$ cells with the administration of control antibody or anti-NK1.1 antibody (n=3 per group). **K.** Representative flow cytometry plots (left) and quantification (right) of $CD4^+$ and $CD8^+$ T cells with the administration of control antibody, anti-CD4 or anti-CD8 antibody (n=3 per group). Mean ± SEM are shown, and P values are calculated by one-way ANOVA (**A, B, D**). or two-sided unpaired Student's t-test (**J, K**). $^*P < 0.05$, $^{**}P < 0.01$, $^{***}P < 0.001$, $^{****}P < 0.0001$, ns, not significant.

### Fig. S10

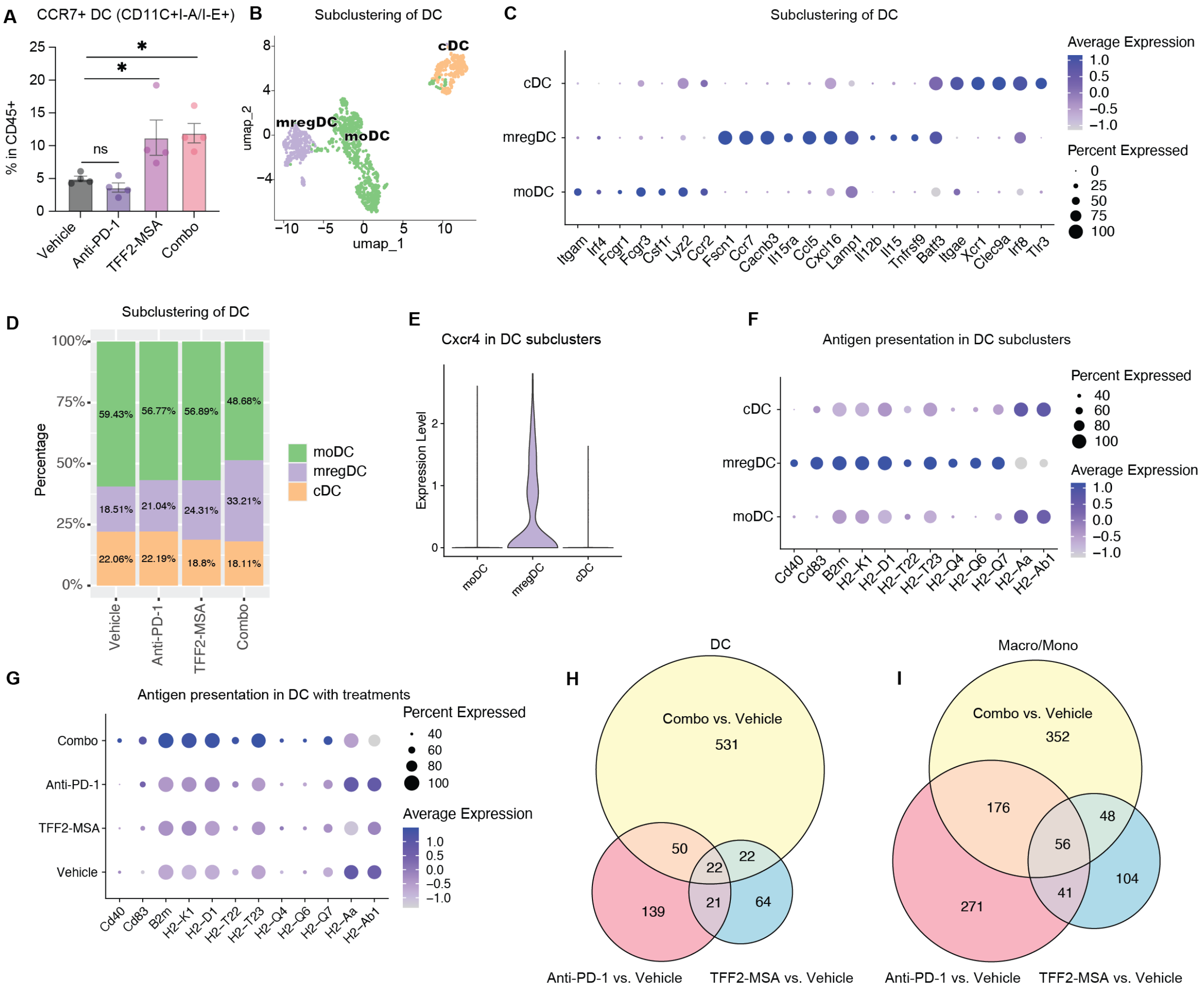


**Fig. S10. Related to Figure 3. Changes of dendritic cells upon the TFF2-MSA and/or anti-PD-1 treatments. A.** Percentage of CCR7$^{+}$ DCs (defined as CD11C$^{+}$I-A/I-E$^{+}$) in CD45$^{+}$ cell of ACKP tumors from the indicated treatment groups (n=4 per group). DC, dendritic cell. **B.** UMAP visualization of DC subclustering. DC. Dendritic cell. moDC, monocyte-derived DC. mregDC, mature immunoregulatory DC. cDC, conventional DC. **C.** Dot plot showing expression of markers used in DC subclustering in each subset. **D.** Stacked bar graphs depicting the frequency of DC subclusters in each indicated treatment group. **E.** Violin plot showing CXCR4 expression in each identified DC subcluster. **F.** Dot plot showing expression of antigen presentation molecules in each identified DC subcluster. **G.** Dot plot showing expression of antigen presentation molecules in DCs from each indicated treatment group. **H.** Venn diagram depicting the differentially expressed genes in tumor DCs with indicated treatment versus vehicle-treated ACKP tumor-bearing mice. Numbers in the Venn diagram indicate the number of differentially expressed genes. **I.** Venn diagram depicting the differentially expressed genes in tumor macrophage/monocytes with indicated treatment versus vehicle-treated ACKP tumor-bearing mice. Numbers in the Venn diagram indicate the number of differentially expressed genes. Mean ± SEM are shown, and P values are calculated by one-way ANOVA (**A**). *P < 0.05, ns, not significant.

### Fig. S11

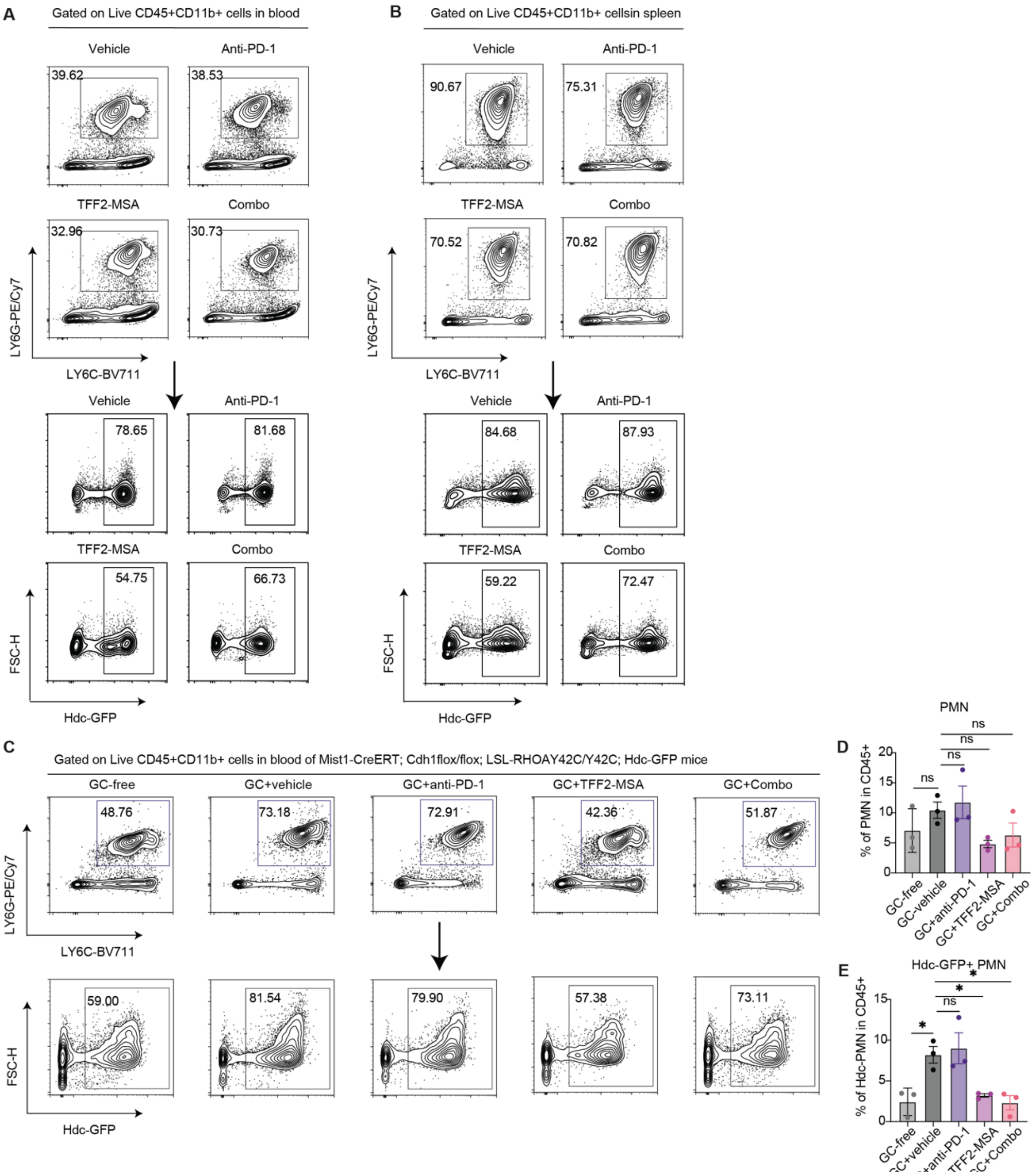
A
Gated on Live CD45+CD11b+ cells in blood
Vehicle
Anti-PD-1
39.62
38.53
TFF2-MSA
Combo
32.96
30.73
LY6G-PE/Cy7
LY6C-BV711
78.65
81.68
54.75
66.73
FSC-H
Hdc-GFP
B
Gated on Live CD45+CD11b+ cellsin spleen
90.67
75.31
70.52
70.82
84.68
87.93
59.22
72.47
C
Gated on Live CD45+CD11b+ cells in blood of Mist1-CreERT; Cdh1flox/flox; LSL-RHOAY42C/Y42C; Hdc-GFP mice
GC-free
GC+vehicle
GC+anti-PD-1
GC+TFF2-MSA
GC+Combo
48.76
73.18
72.91
42.36
51.87
59.00
81.54
79.90
57.38
73.11
D
PMN
% of PMN in CD45+
ns
E
Hdc-GFP+ PMN
% of Hdc-PMN in CD45+
*

**Fig. S11. Related to Figure 4. TFF2-MSA reduced PMN-MDSCs in the blood and spleen of ACKP tumor-bearing mice. A-B.** Representative flow cytometry plots of $LY6G^{+}LY6C^{-/low}$ PMN and Hdc-$GFP^{+}$ PMN-MDSCs in the blood (A) or spleen (B) of the ACKP tumor model with the indicated treatments. **C, D, E.** Representative flow cytometry plots of $LY6G^{+}LY6C^{-/low}$ PMN and Hdc-$GFP^{+}$ PMN-MDSCs in the blood of Mist1-CreERT; $Cdh1^{flox/flox}$; LSL-$RHOA^{Y42C/Y42C}$; Hdc-GFP model bearing autochthonous GCs with the indicated treatments (C). Mice of the same genotype and age without tamoxifen induction were used as tumor-free controls (n=3 per group). Quantifications of $LY6G^{+}LY6C^{-/low}$ PMN and Hdc-$GFP^{+}$ PMN-MDSCs are shown respectively in D and E. Mean ± SEM are shown, and P values are calculated by one-way ANOVA (**D, E**). *$P < 0.05$, ns, not significant.

**Fig. S12**

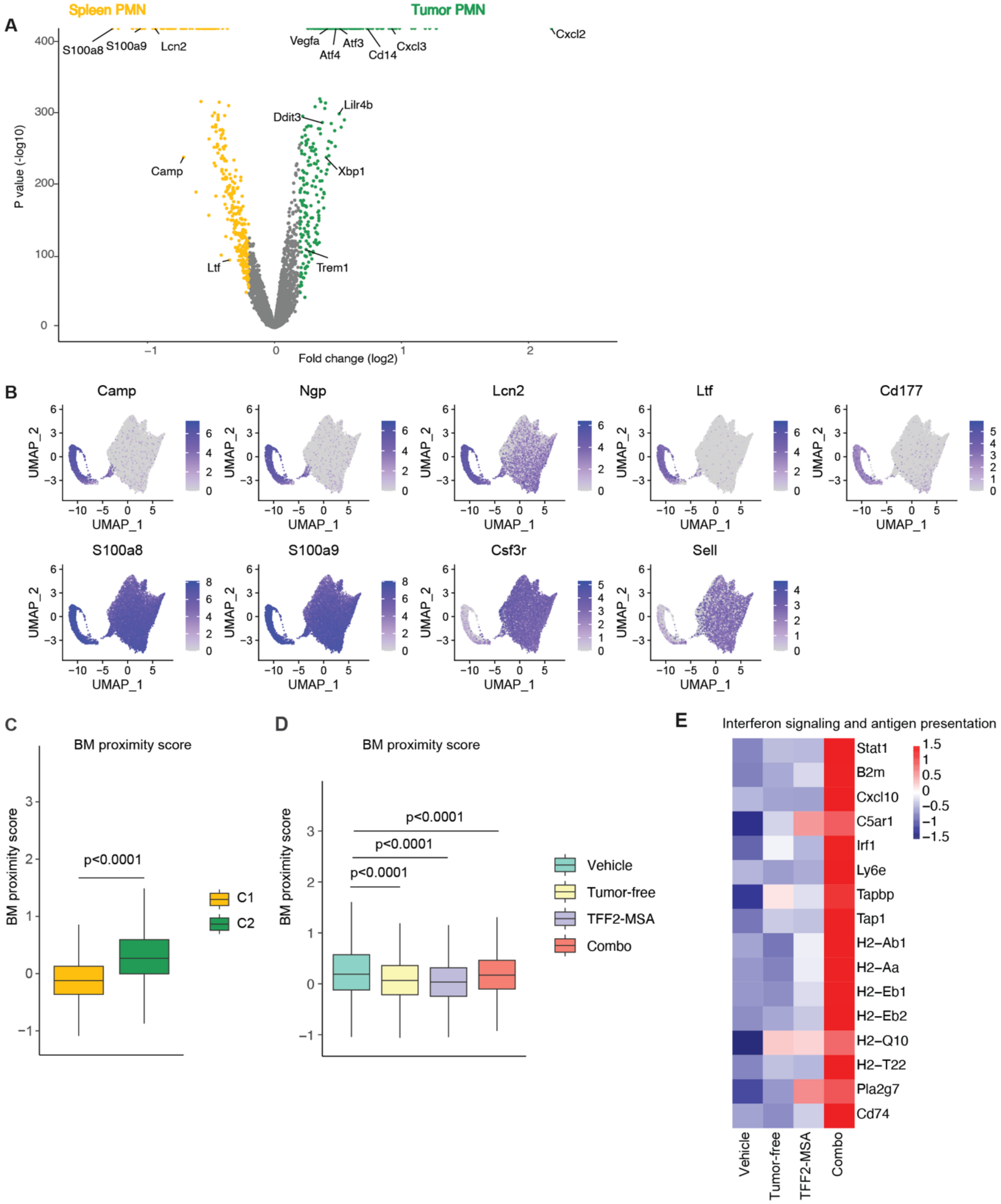


**Fig. S12. Related to Figure 4. TFF2-MSA modulated spleen PMN of ACKP tumor-bearing mice. A.** Volcano plot for differentially expressed genes ($log_2FC \geq 0.2$ and adjusted p value$\leq$ 0.01) in the spleen versus tumor PMNs from vehicle-treated ACKP tumors detected by scRNA-seq. The x axis shows the calculated fold change (FC) of genes upregulated (yellow) or downregulated (green) in spleen PMNs versus tumor PMNs. The y axis indicates the calculated adjusted p value. **B.** UMAP visualization of expression of selected genes enriched in C2 (*Camp, Ngp, Lcn2, Ltf, Cd177, S100a8, S100a9*) or C1 (*Csf3r, Sell*) of spleen PMNs identified in **Fig. 4G**. **C.** Expression of BM proximity score[1] in 2 subclusters of spleen PMNs. BM, bone marrow. **D.** Expression of BM proximity score[1] in spleen PMNs from the indicated treatment groups. BM, bone marrow. **E.** Heatmap showing manually selected gene expressions in the indicated treatment conditions. Mean ± SEM are shown, and P values are calculated by Wilcoxon test (**C, D**).

### Fig. S13

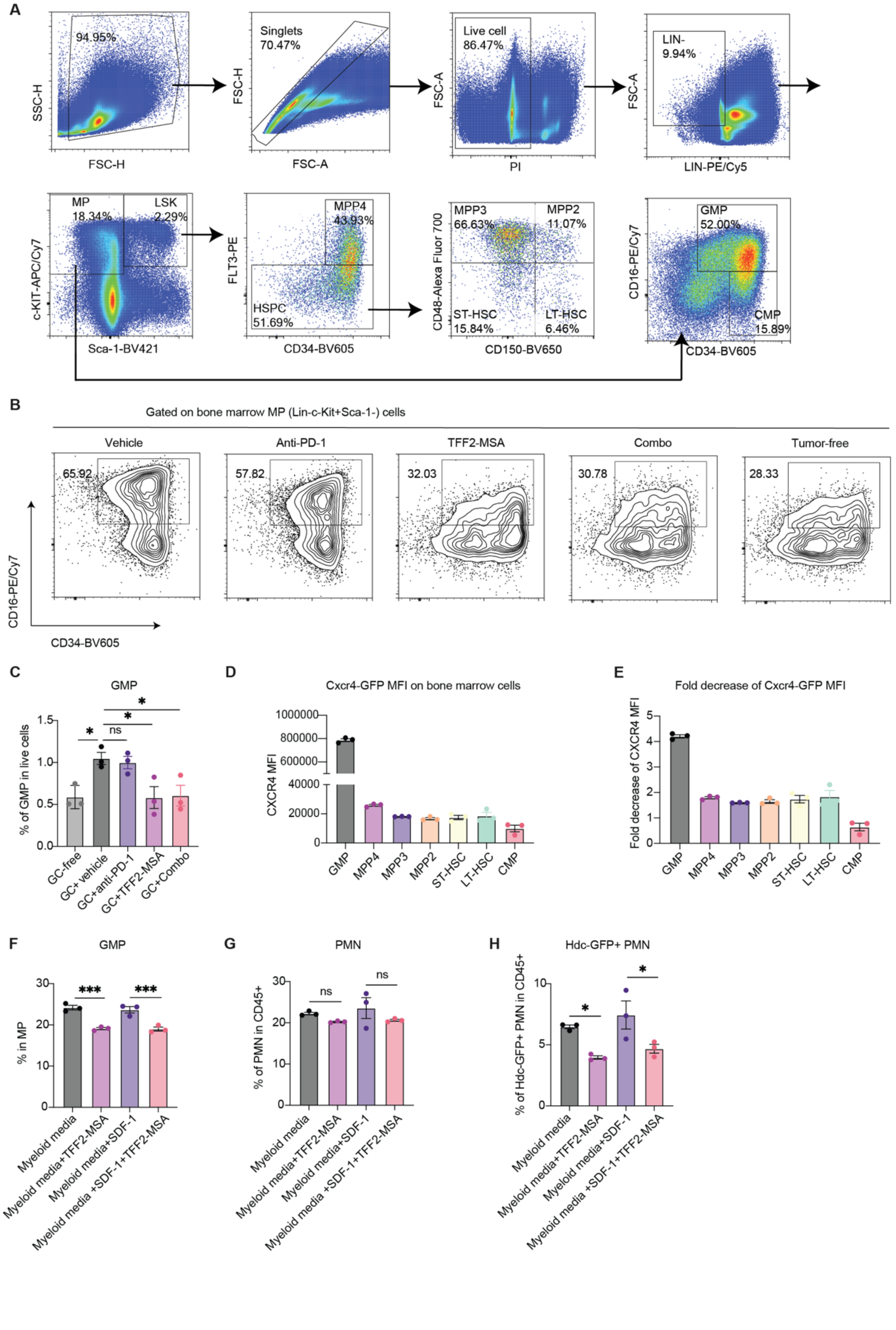

A
94.95%
SSC-H
FSC-H
Singlets
70.47%
FSC-H
FSC-A
Live cell
86.47%
FSC-A
PI
LIN-
9.94%
FSC-A
LIN-PE/Cy5
MP
18.34%
LSK
2.29%
c-KIT-APC/Cy7
Sca-1-BV421
MPP4
43.93%
HSPC
51.69%
FLT3-PE
CD34-BV605
MPP3
66.63%
MPP2
11.07%
ST-HSC
15.84%
LT-HSC
6.46%
CD48-Alexa Fluor 700
CD150-BV650
GMP
52.00%
CMP
15.89%
CD16-PE/Cy7
CD34-BV605
B
Gated on bone marrow MP (Lin-c-Kit+Sca-1-) cells
Vehicle
Anti-PD-1
TFF2-MSA
Combo
Tumor-free
65.92
57.82
32.03
30.78
28.33
CD16-PE/Cy7
CD34-BV605
C
GMP
% of GMP in live cells
GC-free
GC+ vehicle
GC+anti-PD-1
GC+TFF2-MSA
GC+Combo
ns
D
Cxcr4-GFP MFI on bone marrow cells
CXCR4 MFI
GMP
MPP4
MPP3
MPP2
ST-HSC
LT-HSC
CMP
E
Fold decrease of Cxcr4-GFP MFI
Fold decrease of CXCR4 MFI
GMP
MPP4
MPP3
MPP2
ST-HSC
LT-HSC
CMP
F
GMP
% in MP
***
***
Myeloid media
Myeloid media+TFF2-MSA
Myeloid media+SDF-1
Myeloid media +SDF-1+TFF2-MSA
G
PMN
% of PMN in CD45+
ns
ns
Myeloid media
Myeloid media+TFF2-MSA
Myeloid media+SDF-1
Myeloid media +SDF-1+TFF2-MSA
H
Hdc-GFP+ PMN
% of Hdc-GFP+ PMN in CD45+
Myeloid media
Myeloid media+TFF2-MSA
Myeloid media+SDF-1
Myeloid media +SDF-1+TFF2-MSA

**Fig. S13. Related to Figure 4. TFF2-MSA reduced cancer-driven myelopoiesis. A.** Gating strategy for stem cell and progenitors in the bone marrow of tumor-bearing mice. **B.** Representative flow cytometry images of GMPs in the bone marrow of tumor-free mice and ACKP tumor-bearing mice following treatments (n=3). MP, myeloid progenitor ($Lin^-C^-kit^+Sca$-$1^-$). **C.** Quantification of GMPs in the bone marrow of Mist1-CreERT; $Cdh1^{flox/flox}$; LSL-$RHOA^{Y42C/Y42C}$; Hdc-GFP mice with autochthonous GCs following treatments. Mice of the same genotype and age without tamoxifen induction were used as tumor-free controls (n=3 per group). **D.** Quantification of Cxcr4-GFP MFI of bone marrow progenitor and stem cells from Cxcr4-GFP mice (n=3 per group). **E.** Fold decrease of Cxcr4-GFP MFI of bone marrow progenitor and stem cells from Cxcr4-GFP mice (n=3 per group). **F.** Quantification of GMP percentage within MP cells after culture of sorted MP cells with the indicated treatments for 5 days (n=3 per group). MP, myeloid progenitor ($Lin^-c$-$Kit^+Sca$-$1^-$). **G.** Quantification of PMN percentage within $CD45^+$ cells after culture of sorted MP cells with the indicated treatments for 5 days (n=3 per group). **H.** Quantification of Hdc-$GFP^+$ PMN percentage within $CD45^+$ cells after culture of sorted MP cells with the indicated treatments for 5 days (n=3 per group). Mean ± SEM are shown, and P values are calculated by one-way ANOVA (**C, F, G, H**). *P < 0.05, ***P < 0.001, ns, not significant.

Fig. S14

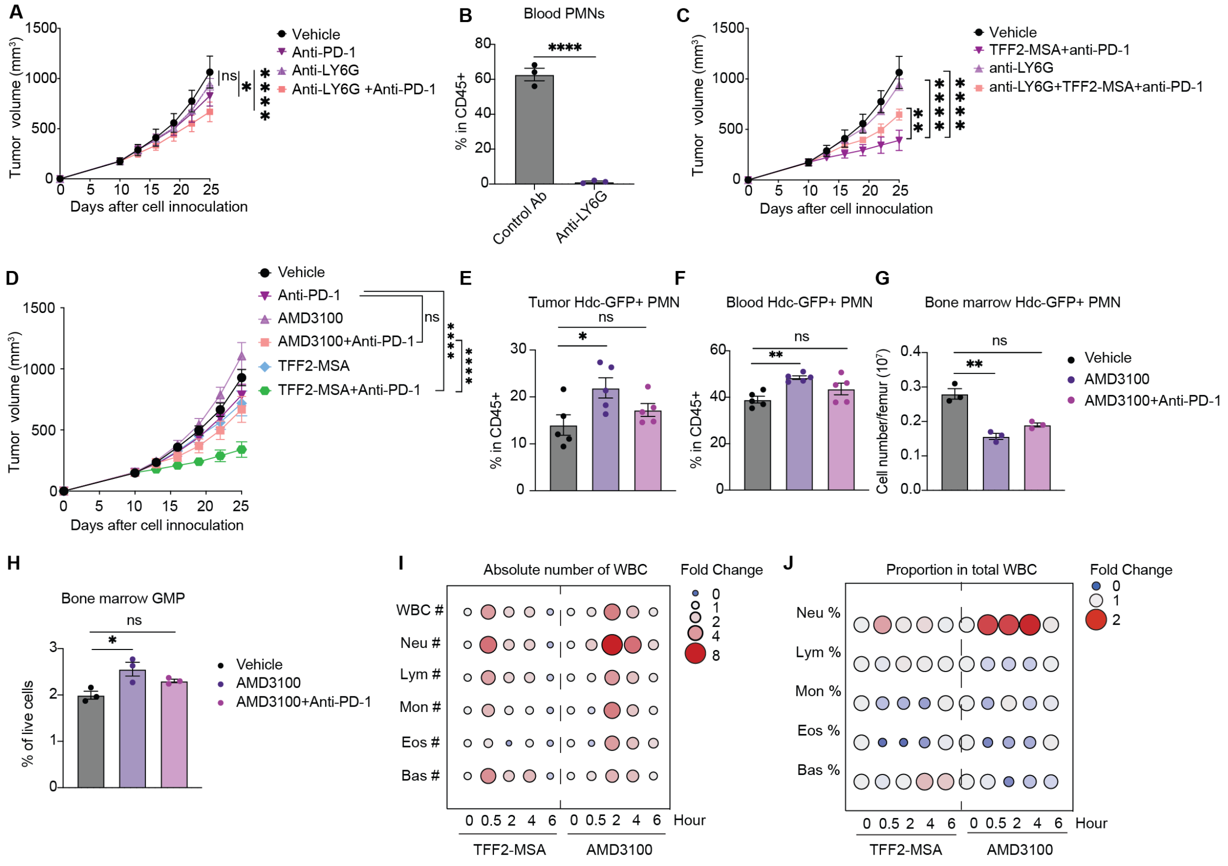


**Fig. S14. Related to Figure 4. TFF2-MSA outperforms total PMN deletion or CXCR4 antagonist in improving anti-PD-1 efficacy. A.** Tumor growth curve of subcutaneous ACKP tumors subjected to the indicated treatments of anti-LY6G and/or anti-PD-1 antibody (n=6 per group). **B.** Quantification of PMN-MDSCs in blood after administration of anti-LY6G antibodies (n=3 per group). **C.** Tumor growth curve of subcutaneous ACKP tumors subjected to the indicated treatments of anti-LY6G and/or TFF2-MSA plus anti-PD-1 combination (n=6 per group). **D.** Tumor growth curve of subcutaneous ACKP tumors subjected to the indicated treatments (n=6 per group). **E, F, G.** Quantification of Hdc-GFP+ PMNs in the tumor, blood, and bone marrow of the subcutaneous ACKP tumors subjected to the indicated treatments (n=3 per group). **H.** Quantification of GMP in the bone marrow of ACKP model with indicated treatments (n=3 per group). **I, J.** Automatic blood test showing absolute number and proportion of each white blood cell (WBC) type within the total WBC after 0-6 hours of intraperitoneal administration of TFF2-MSA (22.5mg/kg) and AMD3100 (10mg/kg) (n=5 per group). Neu, neutrophils. Lym, lymphocytes. Mon, monocytes. Eos, eosinophils. Bas, basophils. Mean ± SEM are shown, and P values are calculated by two-way ANOVA (**A, B, D**). one-way ANOVA (**E, F, G, H**) or two-sided unpaired Student's t-test (**C**). *P < 0.05, **P < 0.01, ***P < 0.001, ****P < 0.0001, ns, not significant.

**Fig. S15**

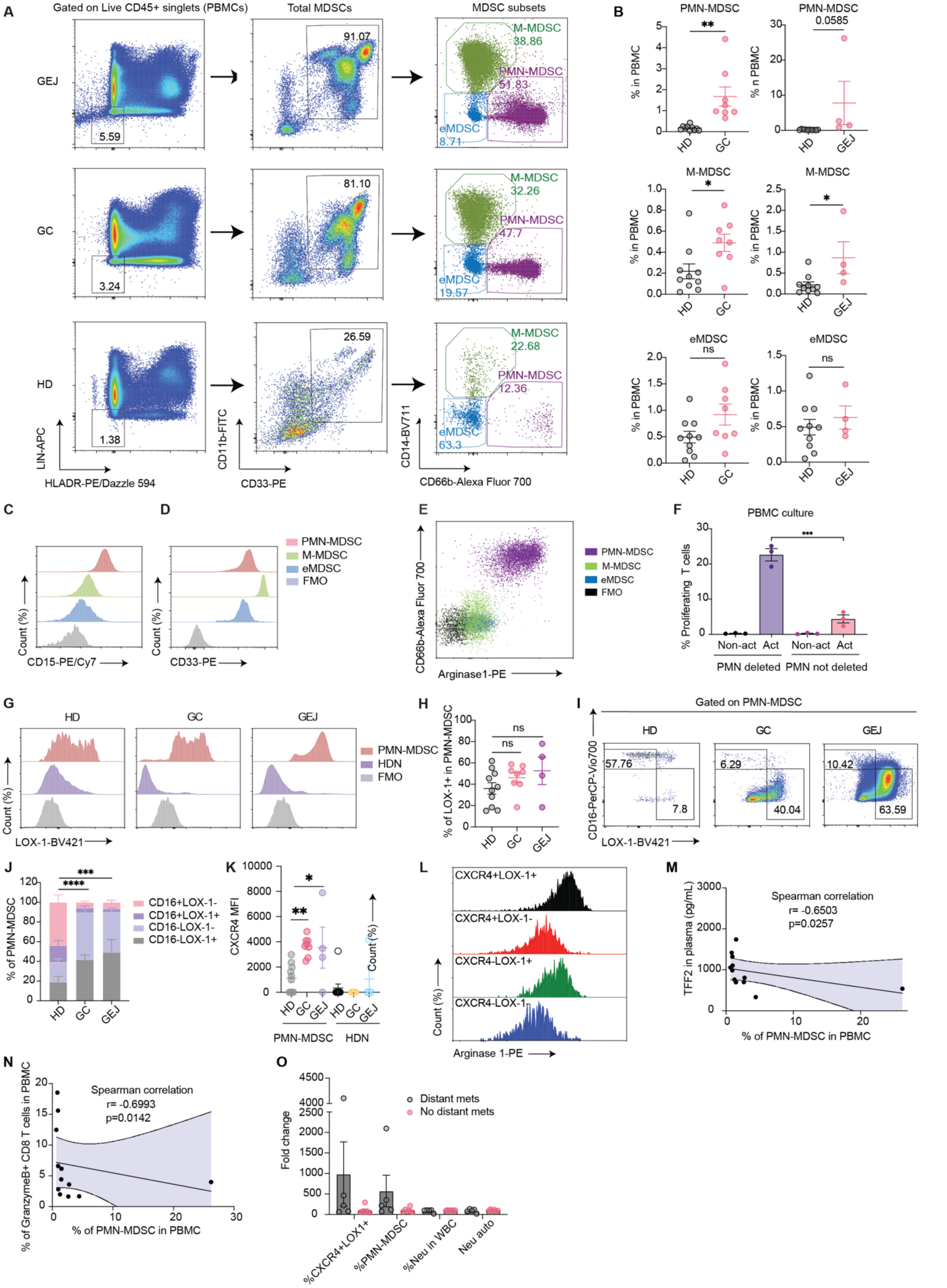

A
Gated on Live CD45+ singlets (PBMCs)
Total MDSCs
MDSC subsets
GEJ
GC
HD
5.59
3.24
1.38
91.07
81.10
26.59
M-MDSC 38.86
PMN-MDSC 51.83
eMDSC 8.71
M-MDSC 32.26
PMN-MDSC 47.7
eMDSC 19.57
M-MDSC 22.68
PMN-MDSC 12.36
eMDSC 63.3
LIN-APC
HLADR-PE/Dazzle 594
CD11b-FITC
CD33-PE
CD14-BV711
CD66b-Alexa Fluor 700
B
PMN-MDSC
M-MDSC
eMDSC
% in PBMC
% n PBMC
0.0585
ns
HD
GC
GEJ
C
D
Count (%)
CD15-PE/Cy7
CD33-PE
PMN-MDSC
M-MDSC
eMDSC
FMO
E
CD66b-Alexa Fluor 700
Arginase1-PE
F
PBMC culture
% Proliferating T cells
Non-act
Act
PMN deleted
PMN not deleted
G
HDN
LOX-1-BV421
H
% of LOX-1+ in PMN-MDSC
I
Gated on PMN-MDSC
CD16-PerCP-Vio700
57.76
7.8
6.29
40.04
10.42
63.59
J
% of PMN-MDSC
CD16+LOX-1-
CD16+LOX-1+
CD16-LOX-1-
CD16-LOX-1+
K
CXCR4 MFI
L
CXCR4+LOX1+
CXCR4+LOX-1-
CXCR4-LOX-1+
CXCR4-LOX-1-
Arginase 1-PE
M
Spearman correlation
r= -0.6503
p=0.0257
TFF2 in plasma (pg/mL)
% of PMN-MDSC in PBMC
N
Spearman correlation
r= -0.6993
p=0.0142
% of GranzymeB+ CD8 T cells in PBMC
O
Distant mets
No distant mets
Fold change
%CXCR4+LOX1+
%PMN-MDSC
%Neu in WBC
Neu auto

**Fig. S15. Related to Figure 5. $CXCR4^+LOX\text{-}1^+$ PMN-MDSCs significantly expanded and negatively correlated with serum TFF2 and $CD8^+$ T cell abundance in GC patients. A.** Gating strategy and representative flow cytometry plots of total MDSC and its three subsets (PMN-MDSC in purple, M-MDSC in green, eMDSC in blue) in the indicated human PBMCs. HD, healthy donor, n=10. GC, gastric cancer, n=8. GEJ, gastroesophageal junction cancer, n=4. **B.** Quantification of PMN-MDSC, M-MDSC and eMDSC in PBMCs from the blood of HD, GC and GEJ (n=10, 8, 4 respectively). **C. D.** Representative histogram of CD15 (C) and CD33 (D) expression on PMN-MDSCs, M-MDSCs and eMDSCs, shown as the normalized count percentage of total population. FMO, fluorescence minus one, serves as gating control. **E.** Representative flow cytometry plots (left) showing Arginase 1 expression in $CD66b^+$PMN-MDSC rather than $CD66b^-$ M-MDSC or eMDSC from the same GC PBMC sample. FMO, fluorescence minus one, serves as a gating control (n=3). **F.** Cell trace violet–labeled PBMC or CD66b-depleted PBMC from GC patients were activated (Act) with immobilized anti-CD3/CD28 and proliferation was determined after 72 hours (n=3 per group). Non-activated (Non-act) PBMCs were used as control. **G.** Representative histogram of LOX-1 expression on PMN-MDSCs and HDN, shown as the normalized count percentage of the total population. FMO, fluorescence minus one, serves as gating control. **H.** Percentage of $LOX\text{-}1^+$ cells in PMN-MDSCs in HD, GC, and GEJ patients (n=10, 8, 4 respectively). **I.** Representative flow cytometry plots of $CD16^+LOX\text{-}1^-$ and $CD16^-LOX\text{-}1^+$ PMN-MDSCs in HD, GC and GEJ patients (n=10, 8, 4 respectively). **J.** Stacked bar graphs depicting the frequency of CD16 and LOX-1 expressed subsets within PMN-MDSCs in HD, GC and GEJ patients (n=10, 8, 4 respectively). Statistical significance indicates the difference of $CD16^+LOX\text{-}1^-$ frequency in HD, GC and GEJ PMN-MDSCs. **K.** Quantification of CXCR4 MFI in PMN-MDSC and HDN populations from HD, GC and GEJ cases (n=10 for HD PMN-MDSC and HDN, n=8 for GC PMN-MDSC and HDN, n=4 for GEJ PMN-MDSC and HDN). MFI, mean fluorescence intensity. **L.** Representative histogram of Arginase 1 expression on subsets of PMN-MDSCs in the blood of GC patients, shown as the normalized count percentage of the total population (n=3). **M.** Correlation of TFF2 serum levels with PMN-MDSC frequency in human PBMCs (n=12 including GC and GEJ). **N.** Correlation of Granzyme $B^+$ $CD8^+$ T cell with PMN-MDSC frequency in GC PBMCs (n=12 including GC and GEJ). **O.** Fold changes of the percentages of $CXCR4^+LOX\text{-}1^+$ PMN-MDSCs in PBMCs, total PMN-MDSCs in PBMCs, neutrophils in WBCs, and automated neutrophil counts in GC and

GEJ patients with distant metastasis compared to those without distant metastasis. Data is presented as mean ± SEM, and P values are calculated by one-way ANOVA (**F, H, J, K**) or two-sided unpaired Student's t-test (**B**). For correlation analysis, The Spearman correlation coefficient was calculated and indicated (**M, N**). *P < 0.05, **P < 0.01, ***P < 0.001, ****P < 0.0001, ns, not significant.

**Fig. S16**

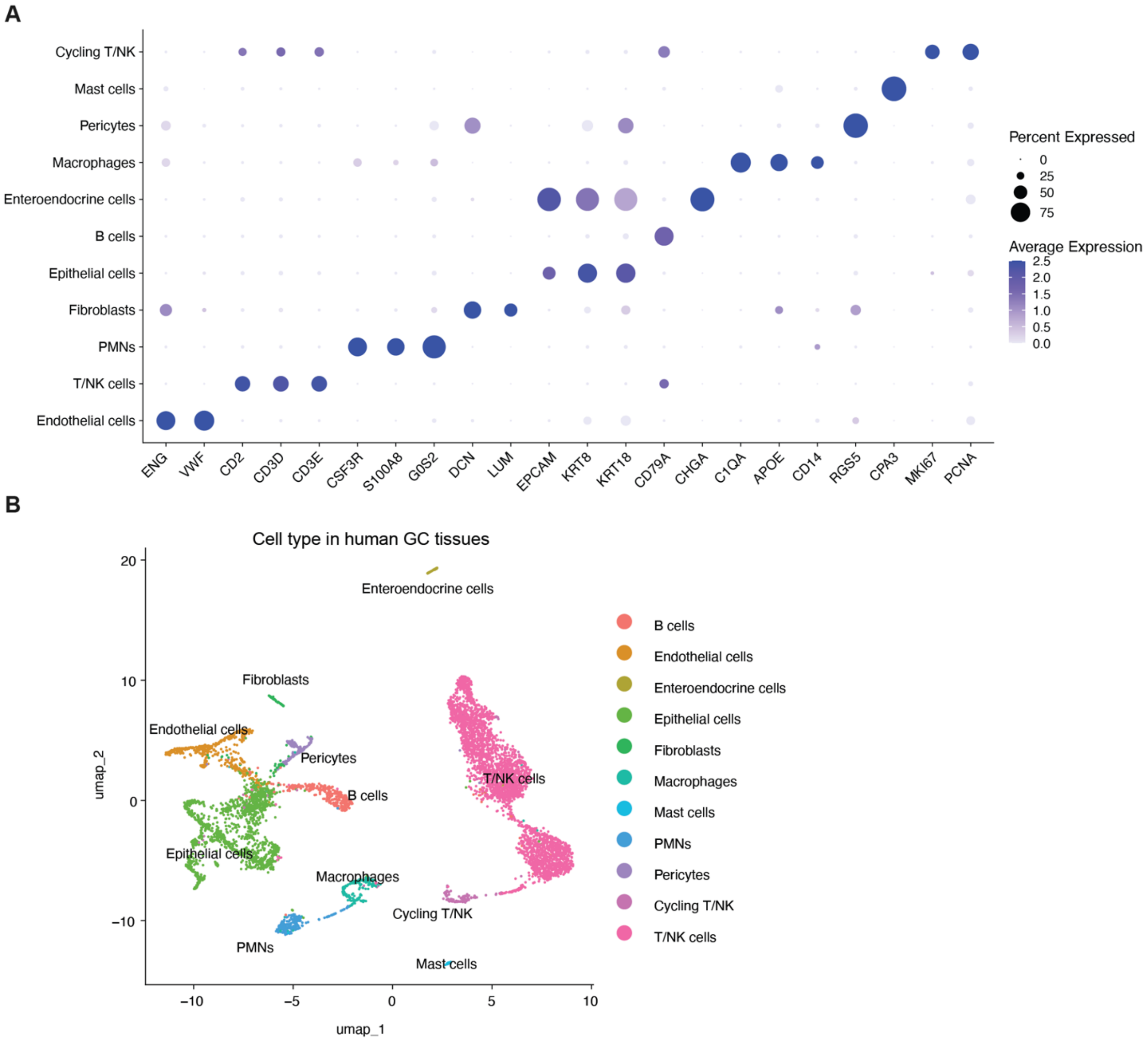


**Fig. S16. Related to Figure 5. PMN analysis in human GC tissues.**

**A.** Gene expression of lineage-defining markers from scRNA-seq analysis of human GC tissues (n=2). The y axis shows cell clusters, the x axis shows characteristic genes in the clusters. **B.** Reclassification of cell types in human GC samples (n=2).

Fig. S17

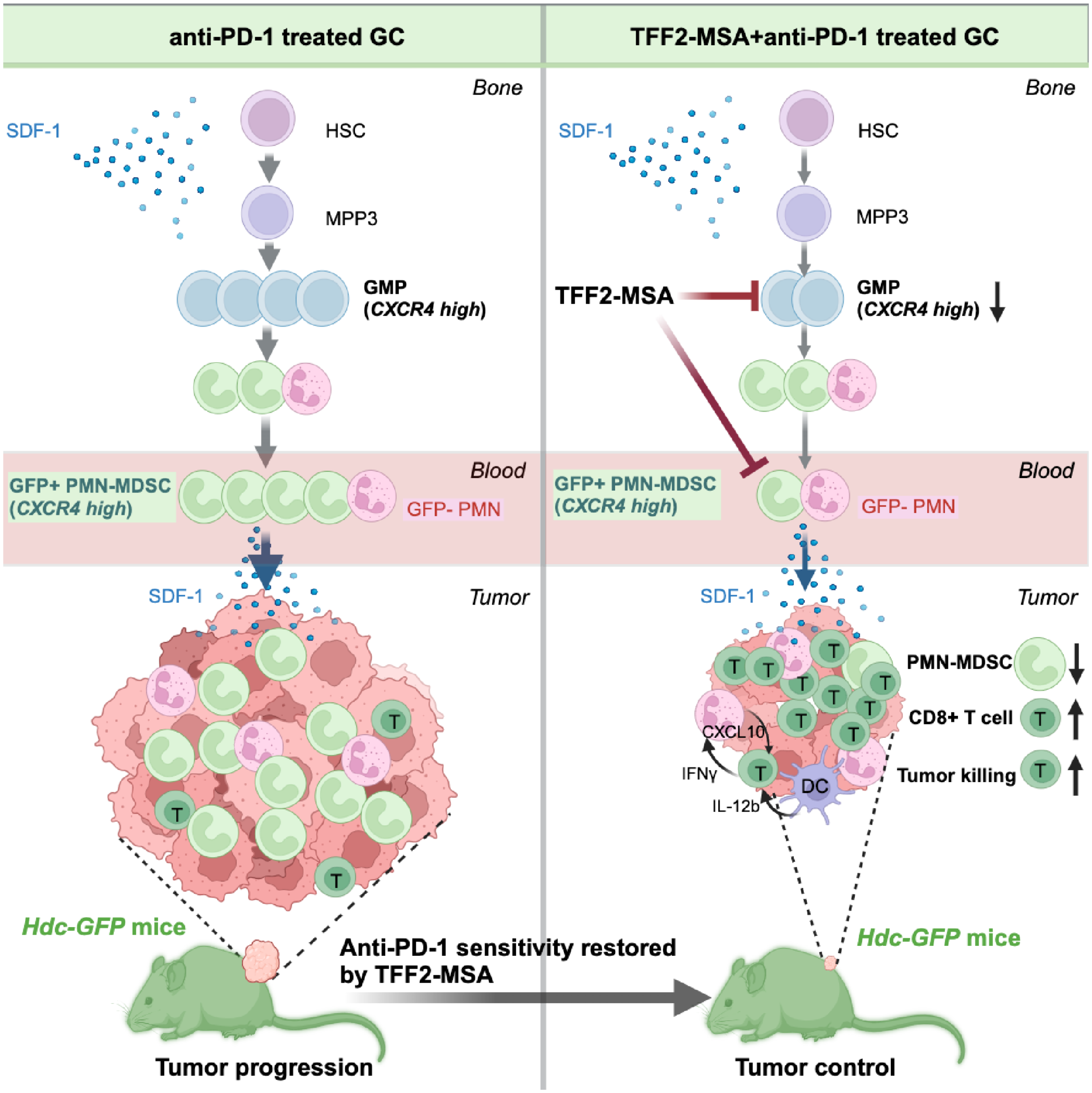


**Fig. S17. Schematic of TFF2-MSA mechanism in effective control of anti-PD-1 resistant GCs.**

Hematopoietic stem cells (HSCs) reside in a specialized niche within the bone marrow, where they receive abundant SDF-1 signals secreted from stromal cells. These cells are responsible for producing blood cells of all lineages through tightly regulated processes. However, in the presence of solid tumors, the balance in bone marrow outputs is disrupted, shifting hematopoiesis toward myelopoiesis. As a result, HSCs preferentially differentiate into myeloid-biased multipotent progenitor 3 (MPP3), which subsequently gives rise to myeloid-committed granulocyte-macrophage progenitors (GMPs), fueling the production of tumor-demanded PMN-MDSCs. Elevated SDF-1 level which is common in solid tumors, can also be an inflammatory signal. In Hdc-GFP transgenic mice bearing syngeneic tumors, Hdc-$GFP^+$ PMN-MDSCs are prematurely released into the bloodstream and recruited to the tumor microenvironment following the SDF-1 gradient and other signals. Their accumulation exerts potent immunosuppressive effects on T cells, thereby promoting tumor progression and resistance to anti-PD-1 monotherapy. TFF2-MSA inhibits both the tumor recruitment of Hdc-$GFP^+$ PMN-MDSCs and the expansion of GMPs in the bone marrow, thereby reducing PMN-MDSC levels and immunosuppression. This leads to an increased number of intratumoral $CD8^+$ T cells. When TFF2-MSA is combined with anti-PD-1 therapy, cytotoxic subsets of $CD8^+$ T cells synergistically expand and mediate effective tumor killing, possibly in cooperation with myeloid cells via an IFNγ-CXCL10-IL-12b feedforward loop. In contrast to CXCR4 antagonist AMD3100, TFF2-MSA acts as a CXCR4 partial agonist, allowing for its selective targeting of GMPs and Hdc-$GFP^+$ PMN-MDSCs which express high levels of CXCR4.
